# Multimodal definition and characterization of the tail of the ventral tegmental area through *Sox14*

**DOI:** 10.64898/2026.11.27.769527

**Authors:** Benjamin M. Muller, Eva Clerc, Pierre-Alexis Derrien, Margot Villechalane, Lucie Mazé, Gwennaëlle Fenollosa, Pierre Hener, Stéphanie Le Gras, Stéphane Doridot, Fanny Faivre, Cristina Sandu, Anil Joshi, Chiara Ebner, Tom Pianezzola, Pierre-Eric Lutz, Emmanuel Darcq, Didier De Saint Jan, Michel Barrot, Jennifer Kaufling

## Abstract

The tail of the ventral tegmental area, or rostromedial tegmental nucleus, is the main inhibitory control of dopamine neurons, but it has lacked a selective marker allowing genetic access. Here we combined bulk and spatial transcriptomics, in situ hybridization, electrophysiology, fiber photometry, viral tracing and behavior to define the structure and characterize its main neuronal population. Expression of Sox14, enriched in the structure in the adult rat and mouse, outlined its full extent, and in the mouse was present in a majority of its GABAergic neurons and more poorly in neighboring regions. In the Allen whole mouse brain atlas, the transcriptomic cell types of these neurons formed a continuous volume along the whole nucleus and yielded additional markers of its subpopulations. Using Sox14-Cre mice, we show that driving these neurons evokes fast, mu-opioid receptor sensitive, GABAA-mediated currents in ventral tegmental dopamine neurons, and in the intact brain rapidly lowers accumbal dopamine. Brain-wide tracing showed that these Sox14 neurons collect input broadly and project narrowly, mainly onto the dopaminergic midbrain. Stimulating them inhibited locomotion and favored the development of place avoidance. Sox14 therefore provides an anatomo-molecular definition of this nucleus and the genetic access needed to dissect its contribution to behavior.

## INTRODUCTION

Scientific progress in brain function and in neuroanatomy is often interdependent. While forebrain and cortical regions are mostly well described, some midbrain to hindbrain parts remain less characterized. Earlier this century, a structure was newly defined at the mesencephalopontine transition in the rat by Jhou and colleagues^1,2^ and by our research group^3,4^. Named RMTg for rostromedial tegmental nucleus or tVTA for tail of the ventral tegmental area, this structure is now considered as the main inhibitory control of dopamine systems^5,6^.

In the rat^2–4,7^ (but not in mice), the tVTA can be visualized by its Fos response to psychostimulants. It was described in this species as a bilateral cluster of GABA neurons rostrally inserted within the posterior VTA, then extending caudally beyond VTA borders and shifting dorsally to be embedded in the superior cerebellar peduncle. While many of its inputs were described to arise from regions mostly similar to VTA ones^2,4^, its outputs appeared more restricted^2,8^. Notably, the tVTA is a main projection of the lateral habenula (LH), this pathway contributing to aversive stimuli processing^9^, and sends dense projections to the VTA and substantia nigra (SNc) dopamine neurons. Electrophysiological studies demonstrated that the tVTA can relay LH inputs to the VTA^10^, exerts inhibitory control on dopamine systems^11,12^ and mediates dopamine response to opiates^13–15^. These features make it a structure of interest for its potential role in various behavioral responses and psychopathologies. Accordingly, the tVTA has been proposed to be implicated in aversive stimuli processing^1^, punishment learning^16,17^, reward prediction error^10^ and risk/reward decision making^18^, dopamine response to drugs of abuse^19^, motor functions^12,20^, motivation to exert effort^21^, despair behavior^22^, pain and analgesia^23,24^, palatable food consumption^25^, and sleep-wake regulation^26,27^. However, up to now, studies were partly limited by both the barrier of species and the precise definition of this brain structure.

Defining the tVTA and its boundaries across species indeed faced the lack of selective anatomo-molecular markers, which partly explains that the tVTA/RMTg is not yet present in brain atlases^28–30^. Besides connectivity features, the tVTA is known to be a GABAergic structure^2–4^, expressing the mu-opioid receptor^2,15^, FoxP1 transcription factor^31,32^, nociceptin neuropeptide^31^ and somatostatin^2^. On the other hand, neurodevelopmental studies have also highlighted molecular markers of the lineage of mesencephalopontine GABAergic subpopulations^32^, but the presence of these developmental markers at adulthood is unclear. In the present study, we combined whole-tissue transcriptomic in rat, RNAscope in rat and mouse, spatial transcriptomic in mouse and data mining of the Allen Brain Bank mouse datasets, to propose a converging anatomo-molecular definition of the rodent tVTA. We then build on these data to use a transgenic mouse line to characterize through brain connectivity, *ex vivo* and *in vivo* electrophysiology, photometry, and behavior the main GABAergic cell population of the tVTA.

## RESULTS

### A transcriptomic screen identifies *Sox14* in the adult tVTA

To identify genes with enriched expression in the tVTA, we labeled the structure by retrograde transport of microbeads injected into the VTA of adult rats, and compared RNAs of the laser-microdissected tVTA against whole brain (**Fig.1a**). The principal component analysis separated the tVTA and whole-brain libraries on its first component (74.9% of the variance, **Supplementary Fig.1**), and 11,624 of the 24,393 tested transcripts were enriched or impoverished at an adjusted *P* below 0.05 (**Fig.1b**). Seven of the twenty most enriched transcripts corresponded to validated markers of monoaminergic neurons (**Fig.1c**; **Supplementary Table1**). The tVTA being predominantly GABAergic^2,4^, these transcripts most likely originated from contamination of the dissected area by neighboring structures, namely the VTA and the raphe nuclei. Myelin-related transcripts were also enriched, eleven validated oligodendrocyte genes (**Supplementary Table1**) accounting for 5.2 to 7.2% of the tVTA libraries against 2.1% of the whole-brain ones, consistent with the peduncular fibers crossing the structure^2^.

**Figure 1.**
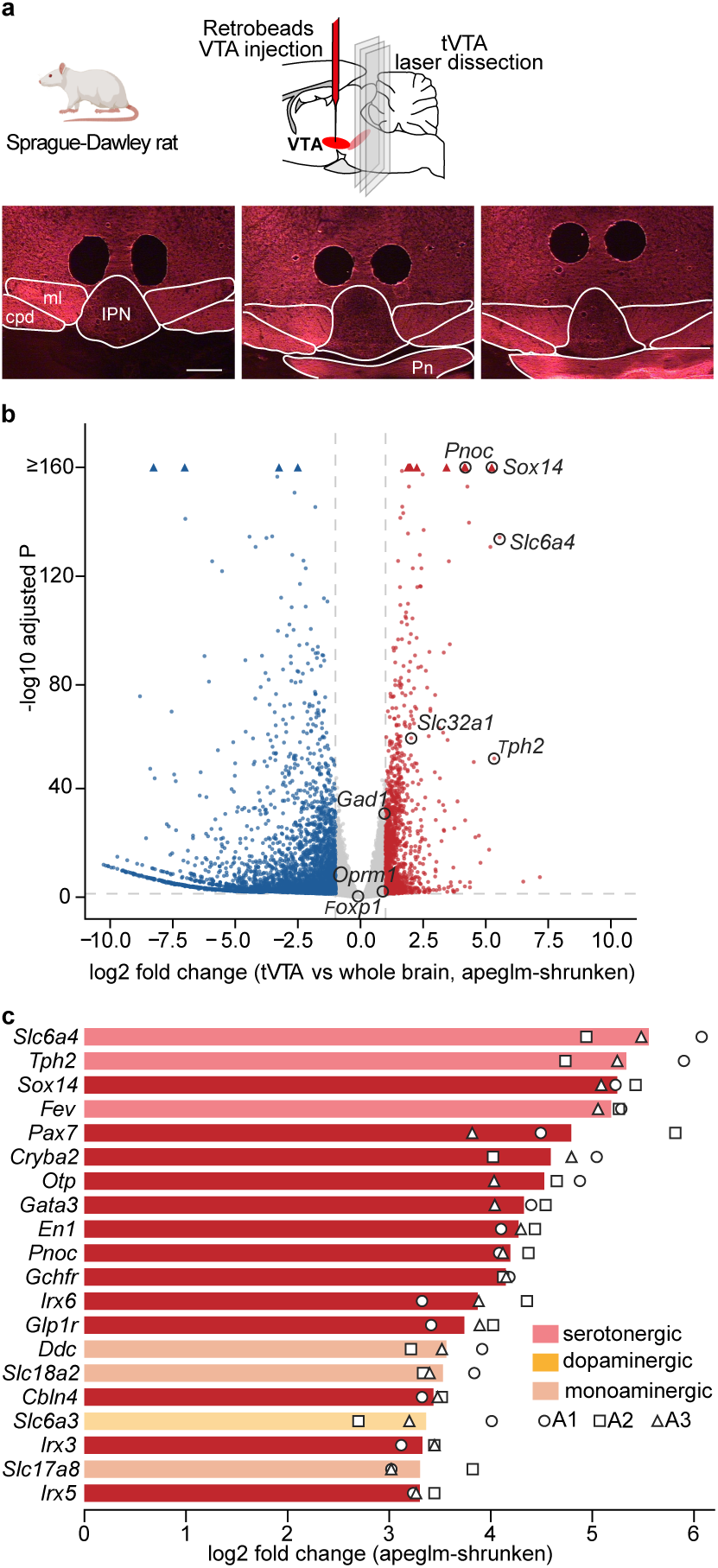
RNA sequencing of the laser-microdissected rat tVTA against whole brain identifies *Sox14* among the most enriched transcripts. **a** Red retrograde microbeads were injected bilaterally into the VTA to label tVTA neurons; one week later the tVTA identified under fluorescence on coronal sections was laser-microdissected. Pictures from three sections of a rat brain, anterior to posterior, illustrating the microdissected area (black holes). **b** Volcano plot of the 24,393 tested genes in the tVTA *versus* the whole brain. Red: enriched in the tVTA; blue: depleted; gray: not significant; dashed lines: adjusted *P* < 0.05 and |log2 fold change| > 1; top row, triangles (with circles when labeled): the thirteen genes above the capped ordinate. Labeled: *Sox14*, the established tVTA markers *Pnoc*, *Foxp1*, *Oprm1*, the GABAergic markers *Slc32a1* and *Gad1* and the serotonergic markers *Slc6a4* and *Tph2*. 1,275 genes were enriched and 3,580 depleted beyond both thresholds (adjusted *P* < 0.05 and |log2 fold change| > 1). *Sox14* was among the most enriched and most significant transcripts, whereas *Foxp1* lay at the center of the distribution and *Oprm1* and *Gad1* at the fold-change threshold. c The twenty most enriched transcripts, ordered by shrunken log2 fold change, among the 863 of the 1,275 enriched transcripts that were protein-coding, detected in all three tVTA libraries and expressed at a mean of at least 5 transcripts *per* million (**Supplementary Table 2**). Bars: shrunken log2 fold change; symbols: log2 ratio of each tVTA library (A1, A2 and A3); pale colors: the serotonergic, dopaminergic and shared monoaminergic markers taken *a priori* from the literature as dissection contamination (**Supplementary Table 1**); dark red: candidate tVTA-enriched transcripts. Seven of the twenty most enriched transcripts were markers of monoaminergic neurons, and *Sox14* was the most enriched of the remaining ones. Group sizes: 3 rats for the tVTA, 3 rats for the whole brain; one library *per* rat. Differential expression assessed by two-sided DESeq2 Wald test with Benjamini-Hochberg correction, unit being the rat. Scale bar: 500 µm (**a**). cpd, cerebral peduncle; IPN, interpeduncular nucleus; ml, medial lemniscus; Pn, pontine nuclei.

Of the established tVTA markers, only *Pnoc*^31^, encoding prepronociceptin, was among the twenty most enriched genes (**Fig.1c**). Three others lay outside the enriched set: *Foxp1* expression in the tVTA did not differ from whole brain (log2 fold change -0.10, adjusted *P* 0.49), while *Oprm1* and *Gad1* fell just below the twofold threshold (0.89 and 0.96), in accordance with their wide expression elsewhere in the brain.

With the monoaminergic genes set aside, *Sox14*, encoding a developmental transcription factor of a GABAergic neuron lineage^33^, was the first enriched transcript of the screen, at 5.24 in log2 fold change and 34.9 against 1.3 transcripts *per* million, and the fifth by adjusted *P* (**Supplementary Table2**). The tVTA *versus* whole-brain contrast reports regional enrichment rather than cell-type selectivity, but identified *Sox14* as a candidate selective marker of tVTA neurons in the adult brain, a hypothesis we tested by *in situ hybridization* and spatial transcriptomics.

### *Sox14* is an anatomo-molecular marker of the tVTA

Transcriptomic analysis identified *Sox14* as selectively enriched in the adult tVTA (**Fig.1**). To validate these findings, we next performed duplex chromogenic *in situ* hybridization (RNAscope ISH) for *Sox14* and *Gad1* on rodent coronal sections of the midbrain and rostral pons. *Sox14* and *Gad1* co-labeled a compact bilateral group of neurons that outlined the tVTA over its full rostro-caudal extent in both the mouse (**Fig.2a**) and the rat (**Supplementary Fig.2a**). Unlike *Gad1*, *Sox14* labeling was not detected in the VTA or the SNc and was very low in raphe nuclei. Outside the tVTA, *Sox14* also labeled a glutamatergic population of the central subnucleus of the interpeduncular nucleus^32^ and a *Gad1*+ population of the superior colliculus^34^, both clearly distinct from the tVTA (**Fig.2a**). Three-dimensional reconstruction revealed a continuous bilateral horn, widening caudo-dorsally, over at least 840µm rostro-caudal extent in the mouse and over 1.2mm in the rat (**Fig.2b**; **Supplementary Fig.2b**). The tVTA displayed a similar position and overall shape in both species, the two horns lying closer together in the mouse. *Sox14* therefore provided a native molecular delineation of a structure previously lacking a precise anatomical marker, particularly in the mouse.

**Figure 2.**
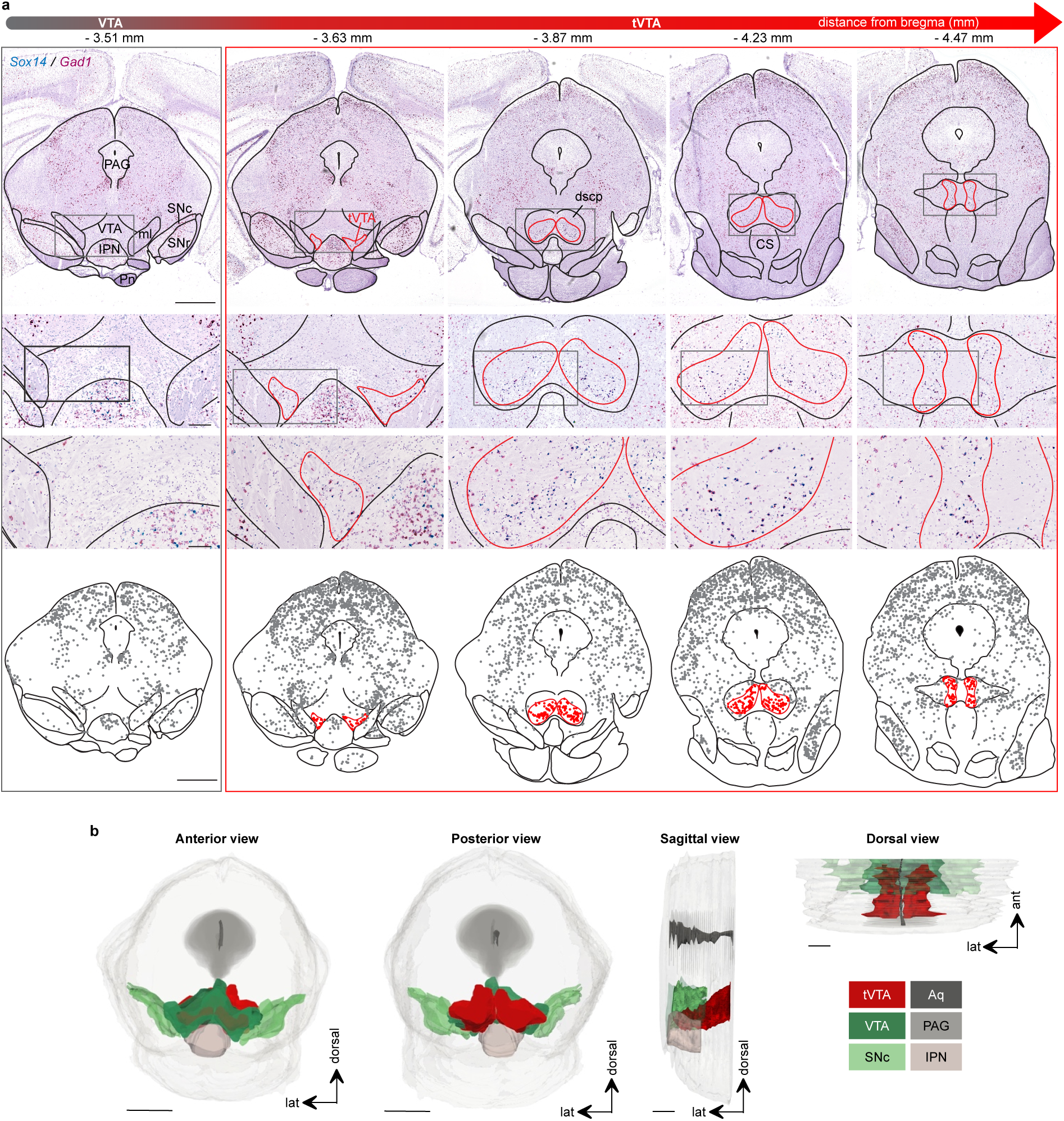
*Sox14*/*Gad1 in situ* hybridization delineates the mouse tVTA as a bilateral horn that widens caudally. **a** Serial coronal sections along the mesopontine rostro-caudal axis, processed for duplex chromogenic RNAscope *in situ* hybridization against *Sox14* (blue) and *Gad1* (red) mRNA, with hematoxylin counterstain. The leftmost column is a reference level through the VTA, rostral to the tVTA (-3.51 mm from bregma); the four columns framed in red span the tVTA, from -3.63 to -4.47 mm from bregma. From top to bottom: whole coronal sections; the boxed VTA/tVTA region at two increasing magnifications; and the corresponding delineation of the tVTA (red outline) relative to the VTA and adjacent structures. The *Sox14*/*Gad1* double-labeled cell cluster defined the tVTA from its rostral pole, embedded within the caudal VTA, to its caudal end in the pons. Rostrally the tVTA appeared as a small cluster within the caudal VTA, above the interpeduncular nucleus (IPN) and medial to the medial lemniscus (ml). It shifted progressively dorsally along its caudal extension, following the decussation of the superior cerebellar peduncle (dscp), and ended between the dorsal raphe dorsally and the median raphe (CS) ventrally. Its cross-sectional area increased caudally. **b** Three-dimensional reconstruction of a mouse tVTA (red) generated from serial *Sox14*/*Gad1* sections (one 14 µm section every four; *n* = 1 mouse), covering the rostral 840 µm of the structure (-3.63 to -4.47 mm from bregma). The reconstruction is shown in anterior, posterior, sagittal and dorsal views and positioned relative to the VTA (green), substantia nigra pars compacta (SNc, light green), periaqueductal gray (PAG), cerebral aqueduct (Aq) and interpeduncular nucleus (IPN). Scale bars: 1 mm (**a**, top and bottom rows), 200 µm and 100 µm (**a**, upper and lower magnification rows), 1 mm (**b**, anterior and posterior views) and 250 µm (**b**, sagittal and dorsal views). Pn, pontine nuclei; SNr, substantia nigra pars reticulata.

We next characterized *Sox14*+ cells at mid rostro-caudal levels of the mouse tVTA by multiplex fluorescent RNAscope ISH (Fig.3). *Sox14*+ cells were neurons, 99.5±0.1% of them co-expressing NeuN protein, and they accounted for 87.6±1.5% of the NeuN+ neurons (**Fig.3a**). They were essentially non-dopaminergic, 0.4±0.2% expressing tyrosine hydroxylase (*Th*; **Fig.3b**), and overwhelmingly GABAergic, 96.8±0.7% expressing *Gad1* and 94.7±0.9% *Gad2* (**Fig.3c, d**), the two isoforms being largely co-expressed (Supplementary Fig.3a). Most also expressed the genes for the mu-opioid receptor (*Oprm1*, 73.2±2.5%), prepronociceptin (*Pnoc*, 86.5±1.7%) and somatostatin receptor 1 (*Sstr1*, 79.6±1.2%) (Fig.3e-g). *Oprm1* was almost confined to GABAergic cells, 96.9±1.6% of *Oprm1*+ cells co-expressing *Gad1* (Supplementary Fig.3b). *Sox14* was present in 70.3±2.6% of the *Gad1*+ neurons at these levels, leaving a minor GABAergic subpopulation without *Sox14*.

**Figure 3.**
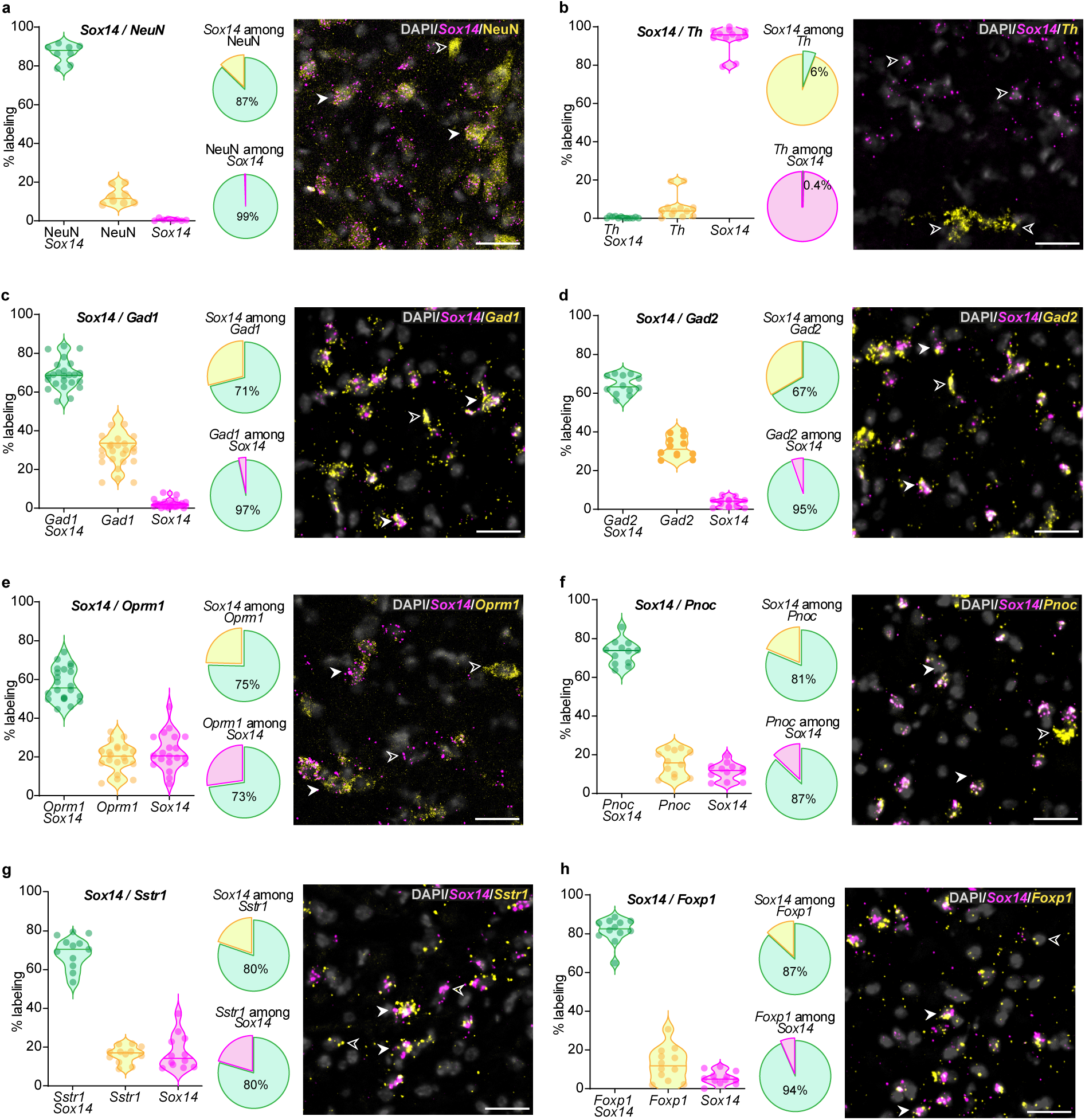
*Sox14*+ tVTA neurons are GABAergic and non-dopaminergic, and most express *Oprm1*, *Pnoc* and *Sstr1*. Markers were analyzed by RNAscope at mid rostro-caudal levels of the mouse tVTA. For each panel: left, proportion of double-labeled, marker-only and *Sox14*-only cells *per* section (one dot *per* section; lines, median and quartiles); middle, pie charts of pooled co-expression across sections (marker among *Sox14*+, and *Sox14* among marker+); right, representative confocal image (*Sox14*, magenta; marker, yellow; DAPI, gray), in which filled arrows indicate double-labeled cells and open arrows single-labeled cells. Co-expression is given below as mean ± s.e.m. *per* section; the values across animals are displayed in the Results. **a** *Sox14*/NeuN: the neuronal marker NeuN was detected by immunohistochemistry in 99.4±0.2% of *Sox14*+ neurons, and *Sox14* in 87.1±1.6% of NeuN+ neurons (*n* = 8 sections, 3 mice, 1,695 cells). **b** *Sox14*/*Th*: *Th* encoding tyrosine hydroxylase was detected in 0.4±0.2% of *Sox14*+ neurons (*n* = 12 sections, 3 mice, 1,590 cells). **c** *Sox14*/*Gad1*: *Gad1* encoding a glutamate decarboxylase was detected in 96.9±0.6% of *Sox14*+ neurons, and *Sox14* in 70.5±1.6% of *Gad1*+ neurons (*n* = 24 sections, 6 mice, 3,536 cells). **d** *Sox14*/*Gad2*: *Gad2* encoding a glutamate decarboxylase was detected in 94.7±0.9% of *Sox14*+ neurons, and *Sox14* in 66.6±1.5% of *Gad2*+ neurons (*n* = 12 sections, 3 mice, 1,621 cells). **e** *Sox14*/*Oprm1*: *Oprm1* encoding the mu-opioid receptor was detected in 72.1±2.4% of *Sox14*+ neurons, and *Sox14* in 74.6±1.9% of *Oprm1*+ neurons (*n* = 20 sections, 6 mice, 2,966 cells). **f** *Sox14*/*Pnoc*: *Pnoc* encoding prepronociceptin was detected in 86.5±1.4% of *Sox14*+ neurons, and *Sox14* in 82.3±2.0% of *Pnoc*+ neurons (*n* = 12 sections, 3 mice, 1,533 cells). **g** *Sox14*/*Sstr1*: *Sstr1* encoding the somatostatin receptor type 1 was detected in 79.5±2.8% of *Sox14*+ neurons, and *Sox14* in 80.5±1.2% of *Sstr1*+ neurons (*n* = 12 sections, 3 mice, 1,539 cells). **h** *Sox14*/*Foxp1*: *Foxp1* was detected in 93.7±1.0% of *Sox14*+ neurons, and *Sox14* in 86.7±2.5% of *Foxp1*+ neurons (*n* = 12 sections, 3 mice, 1,459 cells). *Sox14* labeled most but not all *Gad1*+ neurons at these levels (**c**), leaving a GABAergic subpopulation without *Sox14*. Scale bar: 30 µm.

We also compared *Sox14* with *Foxp1*, encoding another transcription factor proposed as tVTA marker^31,32^. *Foxp1* transcript was detected in 93.7±0.5% of *Sox14*+ neurons (Fig.3h; Supplementary Fig.3c), but FOXP1 protein in only 57.0±5.1% of them (Supplementary Fig.3d). Foxp1 labeling of the tVTA therefore depends on whether transcript or protein is probed.

In the rat, all *Sox14*+ neurons expressed *Gad1* (Supplementary Fig.4a), none expressed *Th* (Supplementary Fig.4b), and a large majority expressed *Oprm1* (83.2±3.1%; Supplementary Fig.4c), which was again almost confined to GABAergic neurons (**Supplementary Fig.4d**). FOXP1 protein was detected in 84.8±3.1% of *Sox14*+ neurons (**Supplementary Fig.4e**). The neurochemical identity of *Sox14*+ tVTA neurons was therefore conserved between rodent species.

### Spatial transcriptomics delineates the mouse tVTA

To validate and expand these findings, we performed Xenium spatial transcriptomics (5,006-gene panel plus 100 custom genes; Supplementary Table5) on five coronal sections through the core of the structure (3 mice). The sections were superimposed in a common frame anchored on the raphe nuclei, and putative tVTA border was traced using two independent strategies (**Fig.4a**). Route A mapped the GABAergic neurons expressing the genes enriched in the rat bulk (*Sox14*, aminergic and oligodendrocyte markers excluded), and route B those expressing three published markers of the adult rodent tVTA^2,31^, *Oprm1*, *Pnoc* and *Foxp1*. The two territories largely overlapped (Dice coefficient 0.74), and their consensus, trimmed of the dorsal raphe nucleus (DR), defined a final tVTA territory (**Supplementary Fig.5a**, **b**). This territory remained stable when one mouse or section was left out or when delineation settings were varied **(Supplementary Table6).** Moreover, the tVTA territory did not overlap other territories drawn by a similar procedure from 13 unrelated neuronal populations **(Supplementary Fig.5b**; **Supplementary Table6).** The contour drawn from *Sox14* alone enclosed 91% of the final territory (**Fig.4a**), thus confirming the converging anatomo-molecular definition of the structure.

**Figure 4.**
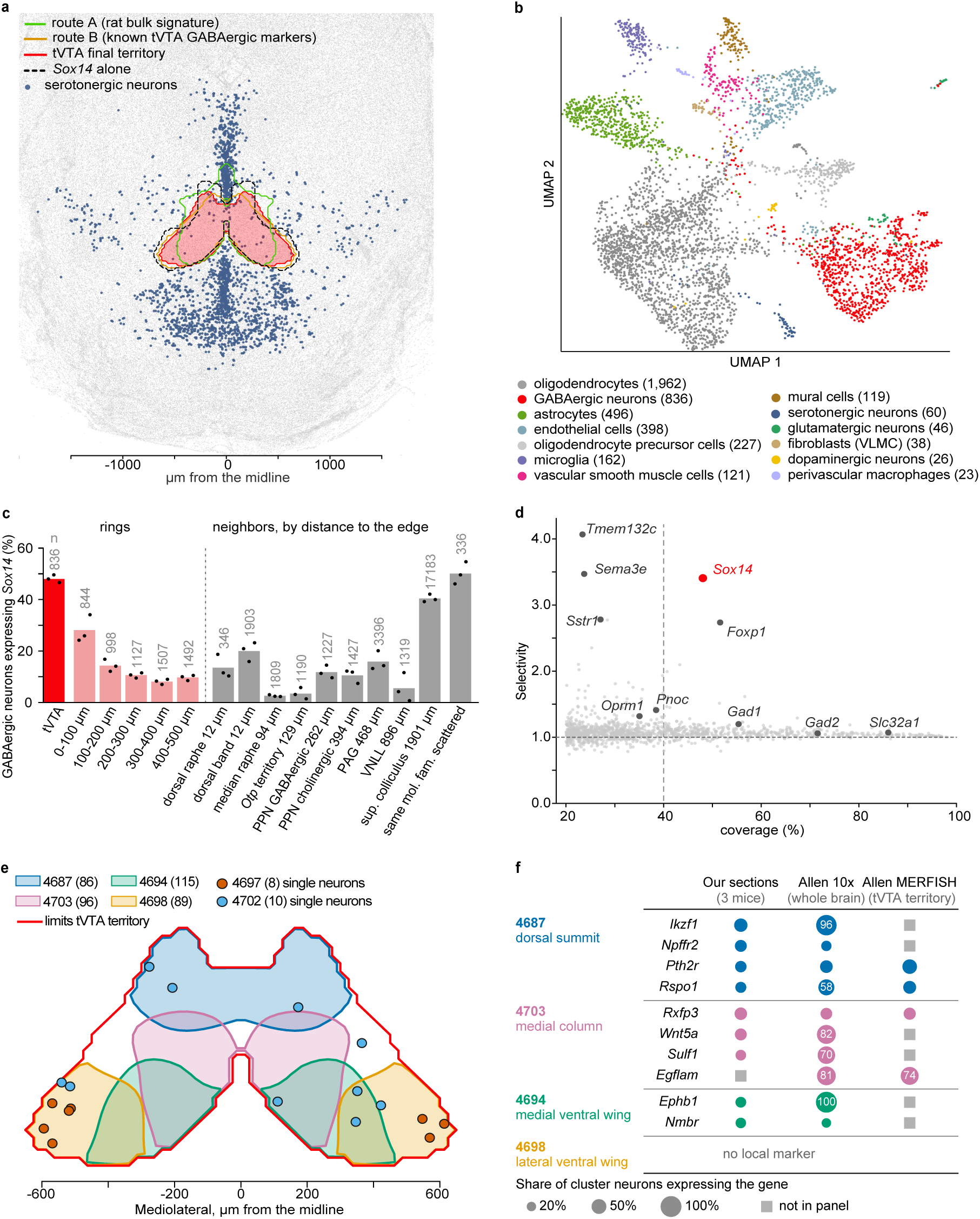
Spatial transcriptomics delineates the mouse tVTA. **a** Mouse tVTA territory (pooled; 5 sections, 3 mice) defined based on rat bulk transcriptomics, on GABAergic neurons with known tVTA transcripts^2,31^ *Oprm1*, *Pnoc* and *Foxp1*, or on *Sox14* expression, in a frame anchored on the raphe nuclei. The two routes defined the tVTA final territory, which was largely matched by the *Sox14* contour. **b** UMAP of cell classes within the tVTA final territory (0.283 mm² *per* hemisphere; 4,514 cells; *n* of cells in brackets). Oligodendrocytes represent 43% of the cells and neurons a fifth, most of them GABAergic (red). **c** Proportion of GABAergic neurons expressing *Sox14* in the tVTA (red), in five 100 µm rings around its edge (pink), and in ten neighboring populations ordered by distance from tVTA edge (gray). Bars, mean of three mice; dots, individual mice; *n*, neurons. *Sox14* fell from 48% in the tVTA to 28% in the first ring, and was at least twice less frequent in eight of the ten neighbors, the superior colliculus and the scattered neurons of the same molecular family excepted. **d** Proportion of tVTA GABAergic neurons expressing genes detected in at least 20% of them (coverage), relative to enrichment of these genes in the tVTA *versus* a 500 µm ring around its edge (selectivity). Gray, all genes; red, *Sox14*. *Sox14* had the highest ratio of the genes expressed in at least 40% of the GABAergic neurons; only two rarer genes, *Tmem132c* and *Sema3e*, exceeded it. **e** Zones of highest concentration of the four largest Allen clusters of the tVTA (subclass 268; red line, tVTA edge). Dots, neurons of the two small clusters, 4697 and 4702. Overlapping zones show intermingled clusters, ordered from a dorsal summit to lateral ventral wings. **f** Coverage of local markers of the four main tVTA Allen clusters. Circle area, proportion of the cluster neurons expressing the genes in Xenium sections (three mice), in the Allen 10x cells (whole brain) and in the Allen MERFISH atlas (tVTA territory, one brain); values written above 50%. Each marker covered a quarter to two fifths of its cluster on Xenium sections.

**Figure 5.**
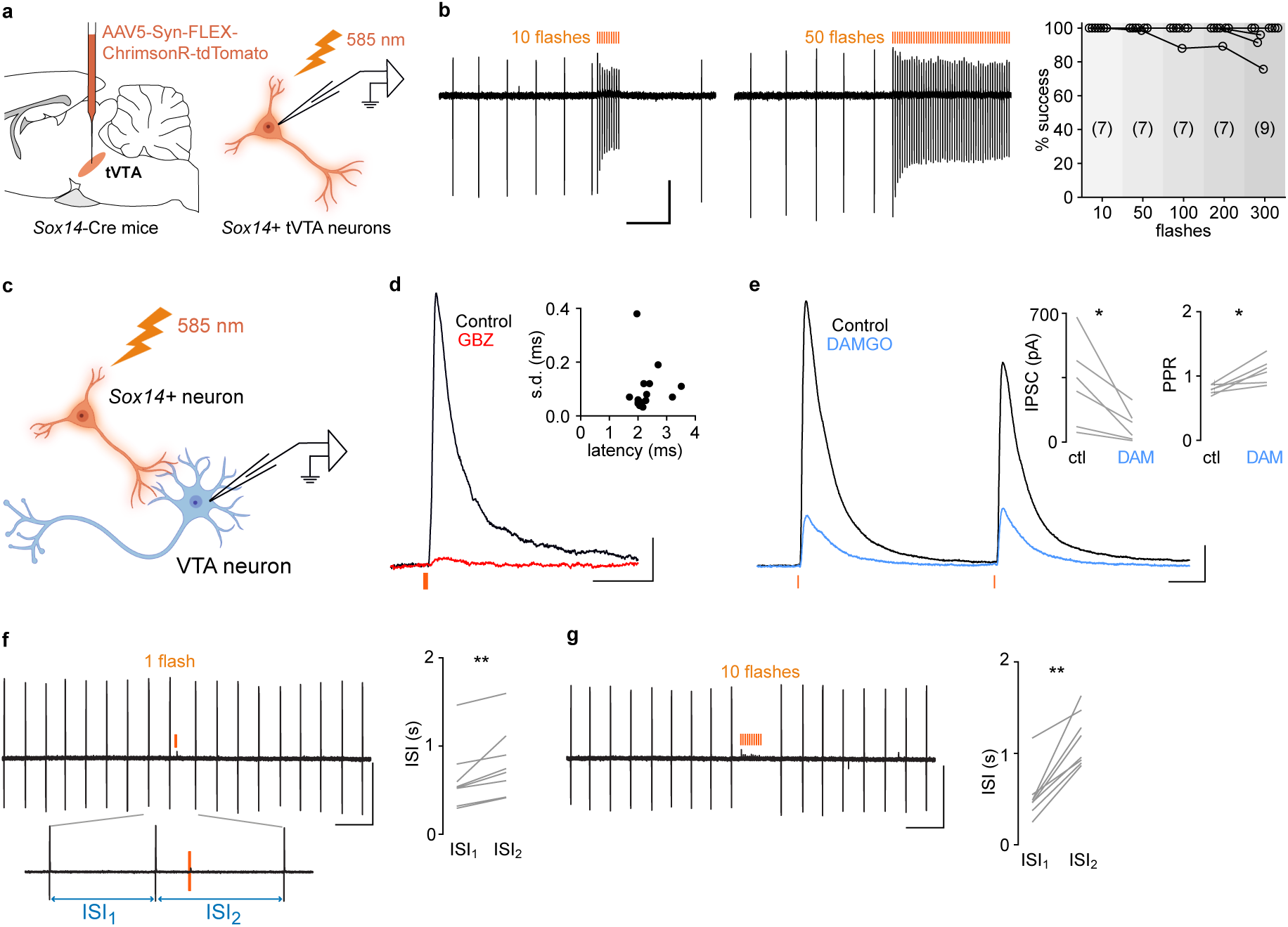
Photostimulation of *Sox14*+ tVTA neurons evokes a fast, mu-opioid-sensitive GABA_A_ IPSC that delays the firing rate of VTA dopamine neurons. **a** Schematic of virus delivery for optogenetic stimulation of *Sox14*+ tVTA neurons and their loose cell-attached or whole-cell recordings. **b** Representative *Sox14*+ tVTA neuron with a regular spontaneous firing and its responses to a train of 10 or 50 flashes every 30 ms. Right: fidelity (% of success) of the evoked spiking against train length, with 10 to 300 flashes at 30 to 40 Hz. Number of neurons in brackets. Fidelity stayed above 90% in eight of the nine neurons tested with 300 flashes. **c** Schematic of synaptic experiments. *Sox14*+ tVTA fibers were photostimulated with 1ms flashes while VTA neurons were recorded in the same horizontal slice. **d** Light-evoked inhibitory postsynaptic current (IPSC) in a representative VTA neuron. The IPSC was blocked by gabazine (GBZ, 5 µM, red) in the three neurons tested (to 4.2±0.7% of control). Inset: latencies of the IPSCs against their s.d. across sweeps, one dot *per* neuron (n = 16). e Pair of IPSCs evoked by two flashes 50 ms apart in a representative VTA neuron under control aCSF or in the presence of DAMGO (1 µM). Insets: first IPSC amplitude (left) and paired-pulse ratio (PPR, right) in the six neurons tested, before (ctl) and after DAMGO (DAM). f Discharge of a putative VTA dopamine neuron stimulated with a single flash. Below: zoom on the two ISI that were compared, the one preceding the flash (ISI1) and the next one that contained it (ISI2). Right: ISI2 and ISI1 in the eight neurons tested. g Same neuron, stimulated with ten flashes at 20 Hz. Detailed statistics are provided in Supplementary Table 11a. \**P* < 0.05, \*\**P* < 0.01. Scale bars: 50 pA, 0.5 s (b); 20 pA, 20 ms (d); 100 pA, 10 ms (e); 100 pA, 1 s (f, g).

Among the 4,514 cells analyzed in this tVTA territory (**Fig.4b**; **Supplementary Table7**), nearly half were oligodendrocytes (1,962, 43%), in line with the myelin enrichment of the bulk screen and the peduncular fibers crossing the structure, and a fifth were neurons (968, 21%), 836 of them being GABAergic (86%). The tVTA is thus a GABAergic nucleus embedded in white matter. The serotonergic and dopaminergic neurons present at the dorsal edge of the territory (**Supplementary Fig.5c**) likely explained the monoaminergic contamination of the tVTA bulk dissection. *Sox14* was detected in 48% of the territory GABAergic neurons (404 neurons; 47-50% *per* mouse), and its presence dropped rapidly beyond the edge of the territory, down to 14% within 200µm (**Fig.4c**; **Supplementary Fig.6**; **Supplementary Table8**). *Sox14* co-detection in *Gad1*+ neurons was lower in the Xenium approach (56-63% *per* mouse; **Supplementary Table8**) than by RNAscope, which may reflect the overall low abundance of *Sox14* transcripts with the Xenium approach using thinner sections (1.99 copies *per* expressing neuron). The Xenium values for *Sox14* as for markers of **Fig.3** could therefore be underestimations (**Supplementary Fig.6c; Supplementary Table8**).

**Figure 6.**
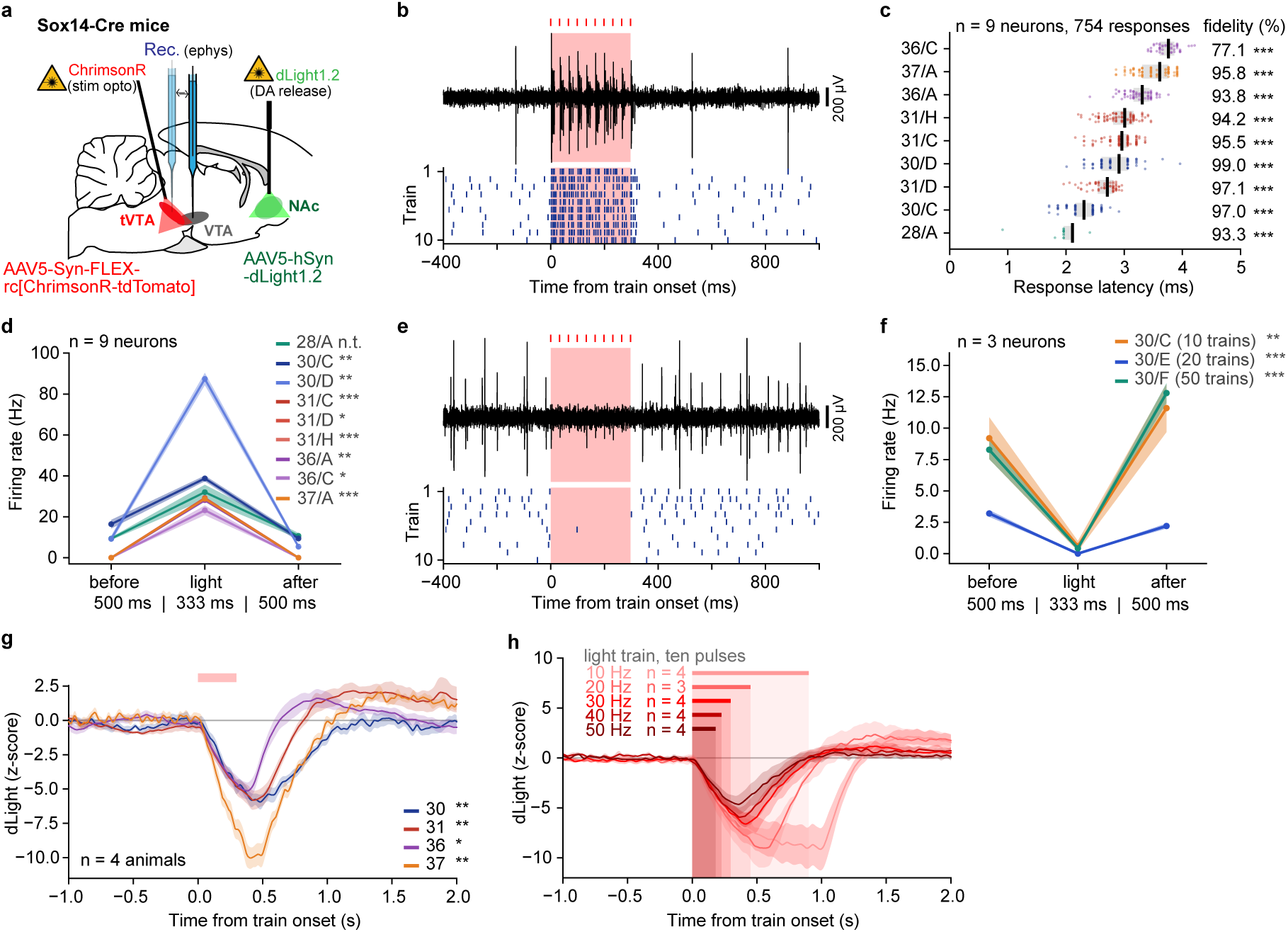
Photostimulation of *Sox14*+ tVTA neurons drives them reliably, silences VTA putative dopaminergic neurons and suppresses NAc dopamine release. **a** Schematic of virus delivery for expression of ChrimsonR in *Sox14*+ tVTA neurons and dLight1.2 in the nucleus accumbens (NAc) of urethane-anesthetized Sox14-Cre mice; single-unit recording in the tVTA and the VTA, fiber photometry in the NAc. **b** Representative photoidentified tVTA neuron. Top: raw signal during one train (5 ms pulse, 30 Hz, 10 pulses); bottom: raster of the 10 trains. **c** Response latency of the nine tVTA neurons, from five mice; one dot *per* response, band: the interquartile range, bar: the median. 754 of 800 pulses answered in 80 trains. Exact binomial *per* neuron. **d** Firing rate of the nine tVTA neurons before (500 ms), during (333 ms) and after (500 ms) the light. Wilcoxon *per* neuron over its trains, comparing the light window (stars) and the after window to the before window. **e** Representative VTA putative dopamine (DA) neuron (same animal as in b) inhibited by tVTA light stimulation. f Firing rate of the three VTA putative DA neurons, same windows and tests as in **d**, 80 trains. g Extracellular DA response at 30 Hz in the four animals, the animal of **b/e** among them. Wilcoxon *per* animal over its ten trials, mean z from 0 to +1 s against the matched -1 to 0 s window; group trough -6.63 z. h DA response to ten pulses at each frequency, mean and s.e.m. (four mice, three for 20 Hz); bars, stimulation train duration. The response end varied with frequency (repeated measures ANOVA over 10, 30, 50 Hz, *F*_(2,6)_ = 54.0, corrected *P* = 0.0026). In (**d**), (**f**) and (**g**) each line is one neuron or animal, mean over its trains or trials, band the s.e.m. Pink in (**b**), (**e**) and (g): light on; one color *per* animal, one tint *per* neuron for an animal. * *P* < 0.05, ** *P* < 0.01, *** *P* < 0.001; n.s., not significant; n.t., not testable. Statistics are provided in **Supplementary Table 11b**.

We also compared the tVTA with the GABAergic neurons of ten neighboring populations delineated over the same sections (**Supplementary Fig.6d**, **7**). *Sox14* was more than twice as frequent in the tVTA as in eight of these other populations, including in dorsal and median raphe nuclei (13% and 3% of GABAergic neurons, respectively). The two exceptions were the superior colliculus, 1.9 mm away (40%), and scattered GABAergic neurons of the same molecular family as the tVTA (50%; **Fig.4c**; **Supplementary Table8**). Within the Xenium panel, *Sox14* appeared as the most selective **gene** expressed by a large share of tVTA neurons (**Fig.4d**; **Supplementary Fig.6b**).

Molecular subpopulations of tVTA GABAergic neurons were then investigated using the Allen whole mouse brain taxonomy^35^. Nearly half of these neurons (404/836) belonged to six clusters of the subclass 268 of GABAergic neurons (**Supplementary Table9**). **The** four largest **clusters** were intermingled but topographically ordered, as a dorsal summit (cluster 4687), a medial column (4703) and a medial and a lateral ventral wing (4694 and 4698; **Fig.4e**) within the tVTA territory. Three of the four clusters showed local markers, each expressed by 26-41% of the neurons of its cluster (**Supplementary Table9**), *Ikzf1*, *Npffr2* and *Pth2r* for the summit, *Wnt5a* for the column and *Ephb1* for the medial wing. The same search in the Allen MERFISH atlas^35^ also retrieved *Pth2r* and *Rspo1* for the summit, and in both the Allen single-cell and MERFISH datasets it added *Egflam*, absent from our panel, for the column (**Fig.4f**; **Supplementary Table9**). *Sox14* was expressed by 59-82% of the neurons in each of the four clusters, and *Pnoc* was the only classical tVTA marker to differ between them, enriched in the column (78%) and rare in the summit (21%; **Supplementary Fig.8**; **Supplementary Table9**). *Sox14* thus marks the tVTA as a whole, not one of its parts.

As the Xenium experiment sampled only the core of the tVTA, we then used the Allen MERFISH atlas^35^ to follow the six GABAergic tVTA clusters along the rostro-caudal axis. They formed a compact territory over seven sections, 1.14mm long and of 0.78mm³, **beginning** where the VTA dopaminergic neurons faded and **running** below the DR, with the same shape (a dorsal summit above two ventral wings) along its extent (**Supplementary Fig.9a**, **b**; **Supplementary Table9**). Although *Sox14* is locally enriched in the tVTA, it should be noted that this structure contains only 2% of the cells of whole brain *Sox14*-rich clusters in the atlas (**Supplementary Fig.9c**).

Together, *in situ* hybridization and spatial transcriptomics delineate the mouse tVTA as a GABAergic nucleus lying between the VTA and the DR, and establish *Sox14* as a selective marker of its neurons within the mesopontine tegmentum. Combined with local viral injection, the Sox14-Cre mouse line^34^ therefore opens genetic access to the tVTA, which we next exploited.

### *Sox14*+ tVTA neurons inhibit VTA dopamine neurons in slices

We injected a Cre-dependent AAV into the tVTA of Sox14-Cre mice to express the excitatory opsin ChrimsonR fused to tdTomato in *Sox14*+ neurons, and recorded them in acute slices (**Fig.5a**). In loose cell-attached (LCA) recordings, over half of the twenty-one recorded neurons were spontaneously active, seven regularly (interspike interval coefficient of variation 0.13-0.22) and five irregularly (0.5-2.5), at rates from <1 to 11Hz. They all responded to a brief flash with a fast-occurring action potential, and trains of 10-300 flashes at 30-40Hz entrained firing with high fidelity (**Fig.5b**). In whole-cell configuration, depolarizing steps evoked irregular spiking in eight neurons and a short burst of spikes at step onset in five (**Supplementary Fig.10a**). Most of the nine neurobiotin-filled neurons displayed a fusiform soma (**Supplementary Fig.10b**).

The tVTA being known to control dopamine systems^2,4^, we then photostimulated *Sox14*+ tVTA fibers in horizontal slices while recording VTA neurons (**Fig.5c**). A single flash evoked a fast outward inhibitory postsynaptic current (IPSC) in twenty-two of the forty-three neurons tested. The IPSC was blocked by gabazine in the three neurons tested (**Fig.5d**) and reversed near the chloride equilibrium potential (**Supplementary Fig.10c**), identifying it as GABA_A_ receptor-mediated. IPSCs were large, with a fast onset latency (2.3±0.1ms) and a jitter below 0.4ms in every neuron, consistent with monosynaptic transmission (**Fig.5d**; **Supplementary Table10**). During trains of ten flashes at 20Hz, the IPSC depressed over the first three pulses and then held, indicating a robust connection (**Supplementary Fig.10d**). As tVTA neurons mediate opioid inhibition of dopamine neurons^13,14^, we then tested the mu-opioid receptor agonist DAMGO (1µM), which reduced the IPSC to 26±7% of control and increased the paired-pulse ratio from 0.83±0.03 to 1.15±0.11 (**Fig.5e**; **Supplementary Table11a**), consistent with a presynaptic action. Most responsive neurons were putative dopamine neurons^36,37^, with a long action potential and a slow depolarization delaying the first spike (**Supplementary Fig.10f**; **Supplementary Table10**), and eight of the eleven neurons tested for tyrosine hydroxylase (TH) were positive (**Supplementary Fig.10e**). *Sox14*+ tVTA neurons thus contact VTA dopamine neurons through fast GABA_A_ synapses that can be controlled by presynaptic mu-opioid receptors.

To test the impact of this input on the pacemaker activity of putative dopamine neurons, we recorded them in LCA and compared the interspike interval preceding the photostimulation (ISI1) with the next one containing it (ISI2). A single flash on *Sox14*+ fibers lengthened ISI2 in all eight neurons, from 633±131 to 810±139ms (**Fig.5f**), and ten flashes at 20Hz lengthened it from 572±100 to 1207±106ms (**Fig.5g**; **Supplementary Table11a**). *Sox14*+ tVTA neurons thus delay the next spike of VTA dopamine neurons, a connection we next tested in the intact brain.

### *Sox14*+ tVTA neurons control the VTA-NAc dopamine system *in vivo*

In Sox14-Cre mice we expressed ChrimsonR in *Sox14*+ tVTA neurons and the dopamine sensor dLight1.2 in the nucleus accumbens (NAc), and recorded single units and accumbal dopamine under urethane anesthesia (**Fig.6a**).

Nine tVTA neurons, photoidentified, responded to trains of stimulation (10×5ms pulses, 30Hz) with a median latency of 2.96ms, a median fidelity of 95.5% and a median jitter of 0.240ms (**Fig.6b**, **c**; **Supplementary Table12**). Each neuron responded above the chance level set by its own spontaneous activity (**Supplementary Table11b**), such a jitter being expected from a spike directly driven by the opsin rather than relayed through the network. Under light, eight of the neurons fired at about one spike *per* pulse, and the ninth close to three (**Fig.6d**; **Supplementary Table13**). Six of the nine neurons emitted no spike within the 500ms before or after any train, indicating that their discharge was generated by the light rather than modulated by it (**Supplementary Table13**). This drive held at 50Hz and over longer trains. The three neurons tested at 50Hz answered 138 of the 140 pulses delivered, which did not differ from their 30Hz response (**Supplementary Table13**). When the 30Hz train was extended to thirty pulses, nearly 1s, three neurons kept a mean fidelity of 87.6 to 99.7%, one of them declining from 97 to 74% (**Supplementary Fig.11a**; **Supplementary Table13**). Sox14-Cre therefore provides direct control of identified tVTA neurons, including at the 30Hz 1s trains used in the behavioral experiments.

Further complementing the slice recordings (**Fig.5**), we recorded *in vivo* single units in the VTA of one animal, where three neurons met the classic criteria for putative dopamine neurons^38–40^(**Supplementary Table12**). The 30Hz tVTA photostimulation silenced them (**Fig.6e**, **f**). Over the first 333ms of the train, firing dropped by 93.5, 100 and 94.9% from its 500ms baseline, stopped altogether in 71 of the 80 trains, and resumed in the following 500ms (**Supplementary Tables11b**, **13**). The monosynaptic GABAergic connection characterized in slices therefore suffices, in the intact brain, to silence VTA dopamine neurons.

The NAc is the main output of VTA dopamine neurons^41^. In four of the mice, accumbal extracellular dopamine was also monitored by fiber photometry, before the unit recordings within the same session (**Fig.6a**). The 30Hz stimulation decreased extracellular dopamine in every animal (**Fig.6g**; **Supplementary Table11b**), with an onset of 67±7ms. Recruiting *Sox14*+ tVTA neurons therefore lowers accumbal dopamine release within a tenth of a second.

To characterize this decrease further, we delivered ten pulses at five frequencies (10, 20, 30, 40 and 50Hz), so that the train, from first to last pulse, lasted 900ms at 10Hz and 180ms at 50Hz. Frequency and train duration therefore covaried, and their effects could not be separated. The onset of the decrease did not vary detectably with frequency. The decrease then deepened throughout the stimulation, slowing down only during the longest trains. In all but one recording, it kept deepening for 74 to 216ms after the light stopped (**Fig.6h**; **Supplementary Fig.11b**; **Supplementary Table11b**). The overall response outlasted the light in all 19 recordings and the absence of 405nm isosbestic response excluded an optical artifact (**Supplementary Fig.11c, d**; **Supplementary Table11b**). Driving *Sox14*+ tVTA neurons therefore does not merely trigger the decrease in accumbal extracellular dopamine, it sets how long it lasts.

*Sox14* thus gives selective access to the tVTA in the intact brain, from the spikes of identified neurons to accumbal dopamine release. We next asked where these neurons draw their input from and where else they send it.

### A genetically defined whole-brain connectome of the tVTA

To map the connectivity of *Sox14*+ tVTA neurons, we combined two Cre-dependent tracing strategies in Sox14-Cre mice. Inputs were mapped by retrograde monosynaptically restricted tracing with a glycoprotein-deleted, EnvA-pseudotyped rabies virus^42–44^(2 mice, 502 and 312 starter cells**; Fig.7a**; **Supplementary Fig.12a-d**). Outputs were mapped with a Cre-dependent AAV expressing EGFP (4 mice; **Fig.7b**; **Supplementary Fig.12e-h**), for which 90.25±0.81% of transduced somata lay within the tVTA. All injections were ipsilaterally dominant, with regional densities consistent between animals in both datasets (**Supplementary Fig.12d, h, Fig.13a-e**; **Supplementary Table14**).

**Figure 7.**
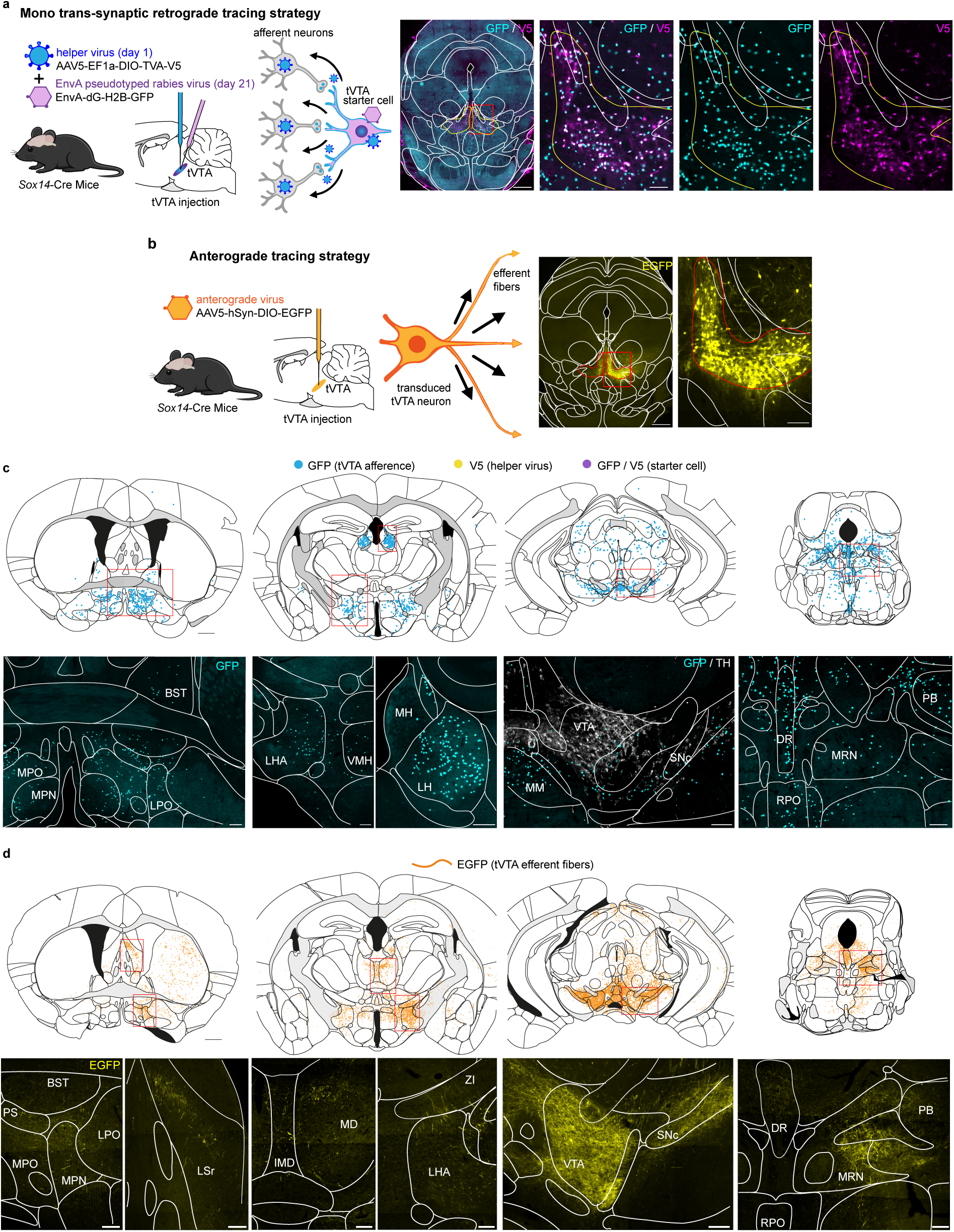
Genetically restricted whole-brain input-output mapping of *Sox14*+ tVTA neurons. **a** Monosynaptic retrograde rabies tracing. A Cre-dependent helper AAV carrying the avian receptor TVA and the rabies glycoprotein (AAV5-EF1a-DIO-TVA-V5) was injected into the tVTA of Sox14-Cre mice, followed 21 days later by an EnvA-pseudotyped, glycoprotein-deleted rabies virus expressing nuclear GFP (EnvA-dG-H2B-GFP). Primary infection was restricted to *Sox14*+ cells co-expressing TVA; starter cells were GFP+/V5+ double positive and transferred the virus to their direct presynaptic partners. Right, injection site at low and high magnification: merge and single channels for GFP (rabies, cyan) and V5 (helper, magenta). **b** Anterograde tracing. A Cre-dependent AAV (AAV5-hSyn-DIO-EGFP) was injected into the tVTA of the same line, labeling *Sox14*+ somata and their axonal projections. Right, injection site at low and high magnification (EGFP, yellow). **c** Examples of afferent labeling at four rostro-caudal levels. Top, two-dimensional reconstruction on coronal atlas templates; cyan, GFP+ input neurons. Bottom, one or two photomicrographs at these same levels; tyrosine hydroxylase immunolabeling (gray) delineates the dopaminergic midbrain. **d** Examples of efferent labeling. Top, two-dimensional reconstruction at the same four levels as in (**c**); orange, EGFP+ fibers. Bottom, one or two photomicrographs at these same levels; yellow, EGFP+ fibers. Red rectangles on the templates indicate the location of the photomicrographs. Rabies tracing, n = 2 mice; anterograde tracing, n = 4 mice; each reconstruction shows one representative animal, #8683 in (**c**) and #6200 in (**d**). Abbreviations: BST, bed nuclei of the stria terminalis; DR, dorsal nucleus raphe; IMD, intermediodorsal nucleus of the thalamus; LH, lateral habenula; LHA, lateral hypothalamic area; LPO, lateral preoptic area; LSr, lateral septal nucleus, rostral part; MD, mediodorsal nucleus of the thalamus; MH, medial habenula; MM, medial mammillary nucleus; MPN, medial preoptic nucleus; MPO, medial preoptic area; MRN, midbrain reticular nucleus; PB, parabrachial nucleus; PS, parastrial nucleus; RPO, nucleus raphe pontis; SNc, substantia nigra, compact part; VMH, ventromedial hypothalamic nucleus; VTA, ventral tegmental area; ZI, zona incerta. Scale bars: 500 µm (low-magnification views in **a** and **b**, templates in **c** and **d**) and 100 µm (all other photomicrographs).

Across the two datasets, 84 regions provided input to *Sox14*+ tVTA neurons and 110 received output, 124 regions in total (**Fig.7c**, **d**, **8a**; **Supplementary Figs.14**, **15**; **Supplementary Table15**). The five densest input regions accounted for 39.3% of the summed afferent density, and the five densest output regions for 61.4% of the summed efferent signal, outputs being consistently more concentrated overall (Gini coefficients 0.757 against 0.653 for inputs). The strongest afferents included the LH, interpeduncular nucleus (IPN), DR and superior central nucleus raphe (CS), whereas outputs converged on the VTA complex (VTA-cx) and SNc, which together accounted for 49.0% of the summed efferent signal. *Sox14*+ tVTA neurons therefore collect broadly and project narrowly.

Rabies labeling was also present within the tVTA itself. Helper-negative GFP+ neurons reached 195.2±33.5 cells/mm^2^ on the injected side (**Fig.8a**), denser than all but seven of the afferent regions. *Sox14*+ tVTA neurons therefore receive an intrinsic input from the tVTA itself, suggested qualitatively for GABAergic neurons^45^ but quantified here for the first time.

**Figure 8.**
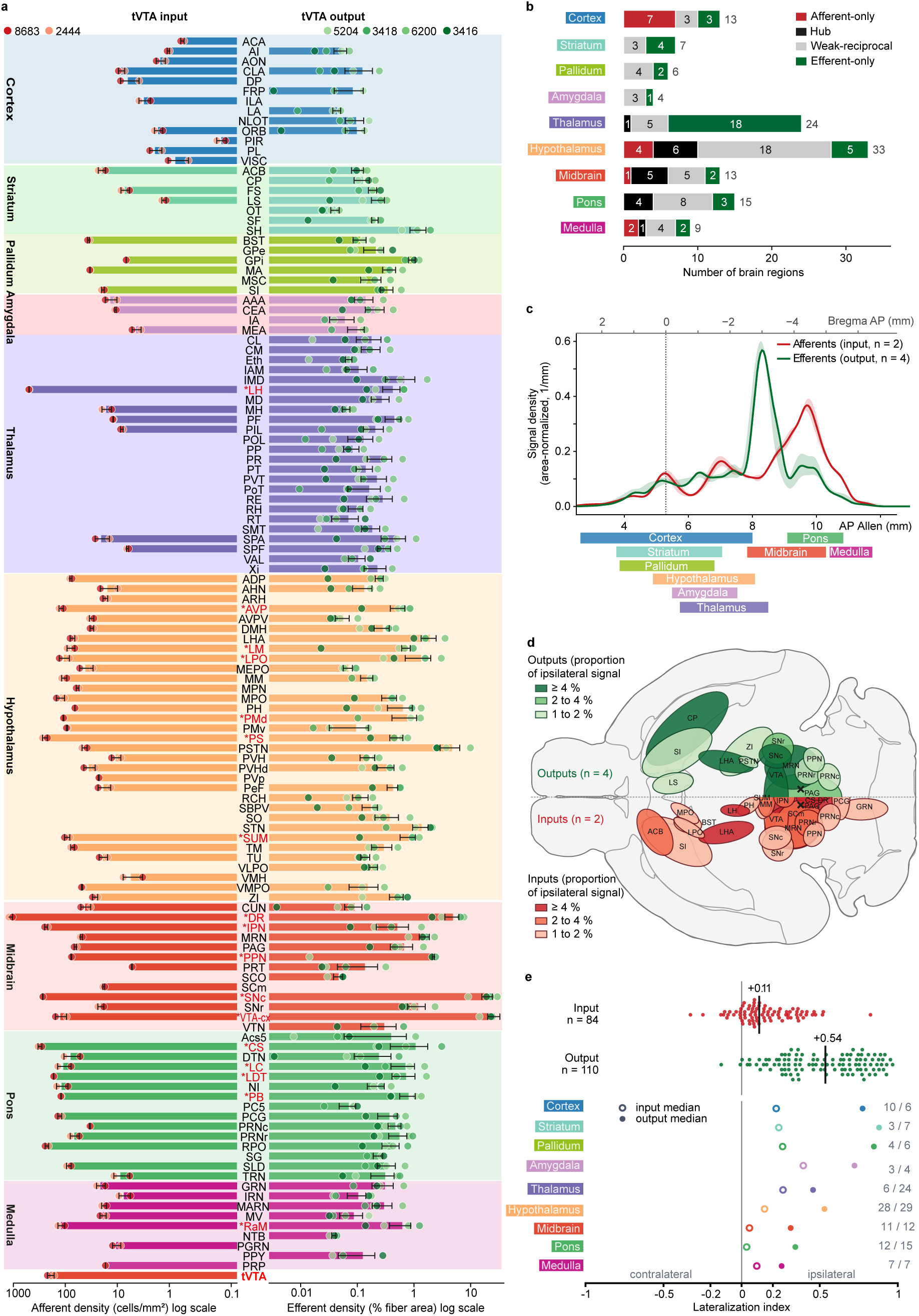
Whole-brain repertoire, reciprocity, rostro-caudal organization and hemispheric lateralization of the *Sox14*+ tVTA connectome. a Ipsilateral mirror-plot of the connectome. Left, inputs (labeled-cell density, cells/mm², log scale, n = 2 mice), and right, outputs (fiber area, % of region area, log scale, n = 4 mice), for the 124 detected regions in nine anatomical super-groups, rostro-caudal order. Bars, mean ± s.e.m.; dots, individual animals; red asterisks, the 17 reciprocal hubs. Bottom row, intra-tVTA afferents (tVTA helper-negative GFP+ cells), not counted among the 124 regions. Nomenclature, Allen Mouse Brain Common Coordinate Framework (CCFv3); full region names, Supplementary Table 15. RaM, medullary raphe nuclei (magnus, obscurus and pallidus merged). b Connectivity composition *per* super-group. Stacked bars, number of regions *per* class; numbers on bars, count *per* class and, at bar end, super-group total. Input/output preference of the 54 unidirectional regions was assessed by exact binomial test (Supplementary Table 11c). Class of each region, Supplementary Tables 15 and 16. c Rostro-caudal profile of ipsilateral input (red) and output (green); Allen CCFv3 below, approximate distance to bregma above. Density profiles were normalized to their whole-brain signal. Line, mean; band, s.e.m. Inputs peak at AP 9.70 mm and outputs at 8.30 mm. Bars below, anatomical extent of each super-group, not of the labeling; colors as in (**a**). The tVTA itself is excluded from both profiles. **d** Main connectome on a horizontal brain drawing, outputs (green) and inputs (red); color depth, proportion of the ipsilateral signal carried by each region, mean of mice; cross, tVTA. **e** Lateralization index, LI = (ipsilateral - contralateral) / (ipsilateral + contralateral), computed *per* animal, then averaged *per* region. Top, one dot *per* region, input (red, 84 regions) and output (green, 110 regions); bar and number, median. Bottom, median *per* super-group, open symbol input, filled symbol output, colors as in (**a**). At right, numbers of indexed input/output regions. Each distribution was tested against zero by two-tailed Wilcoxon signed-rank, unit the region (inputs *P* = 3.2 × 10⁻¹⁰, outputs *P* = 1.2 × 10⁻¹⁹; **Supplementary Table 11c**).

Of the 124 regions, 70 were connected in both directions, including 17 reciprocal hubs that ranked in the top 30% for both input and output, while 14 provided input only and 40 received output only (**Fig.8b**; **Supplementary Tables14**, **15**). Output-only regions thus outnumbered input-only ones **(Supplementary Table11c).** Of the nine anatomical super-groups, the cortex and the thalamus carried this imbalance in opposite directions: seven of the thirteen cortical regions provided only input to the tVTA and none was a hub, whereas 18 of the 24 thalamic regions received only tVTA output, and both showed the lowest reciprocity of the connectome. The pons, midbrain and hypothalamus held 15 of the 17 hubs. *Sox14*+ tVTA neurons therefore exchange with the brainstem and the hypothalamus, but form principally a one-way connection with the cortex, from which they receive, and with the thalamus, to which they project.

The input profile was trimodal (**Fig.8c**). Its highest and broadest peak laid in the caudal midbrain and rostral pons, carried by the DR, CS, midbrain reticular nucleus (MRN) and periaqueductal gray (PAG). A second component, about half as high, was carried by the LH and lateral hypothalamic area (LHA), and a third, the most anterior, by preoptic regions. On the other hand, the output profile had a single narrow peak, centered on the dopaminergic midbrain, and a small pontine component reaching just over a quarter of its height. Inputs reached the tVTA from three levels of the neuraxis, whereas its projection was concentrated in the midbrain immediately rostral to the nucleus.

Both connectivity distributions were ipsilaterally biased (**Supplementary Fig.16a-d**; **Supplementary Tables11c**, **17**), more strongly for outputs and including in the nine super-groups. Indeed, the median lateralization index of the regions connected to the tVTA was +0.11 for inputs and+0.54 for outputs (**Fig.8e**). The input bias was limited, 53 of the 84 input regions remaining bilateral, with only three strongly ipsilateral, the internal globus pallidus (GPi), the zona incerta (ZI) and the visceral area (VISC). Of the 110 output regions, 14 were bilateral and 59 strongly ipsilateral. Overall, no region was strongly contralateral in either dataset. However, a contralateral connection confined to a region subdivision, such as the input from the intermediate layers of the anterior part of the superior colliculus^2,4^, was diluted in the regional average but remained visible on the section-by-section cartographies (**Supplementary Figs.14**, **15**) that followed the labeling rather than the atlas boundaries.

### tVTA stimulation inhibits locomotion and induces avoidance

Activation of LH terminals in the tVTA was reported to induce place avoidance in mice^9^ and in rats^21^. We first confirmed these findings by expressing ChrimsonR in the LH of C57BL/6J mice; 60Hz stimulation^9^ of these terminals led to real-time place avoidance (RTPA) (**Supplementary Fig.17a**, **b**).

**Figure 9.**
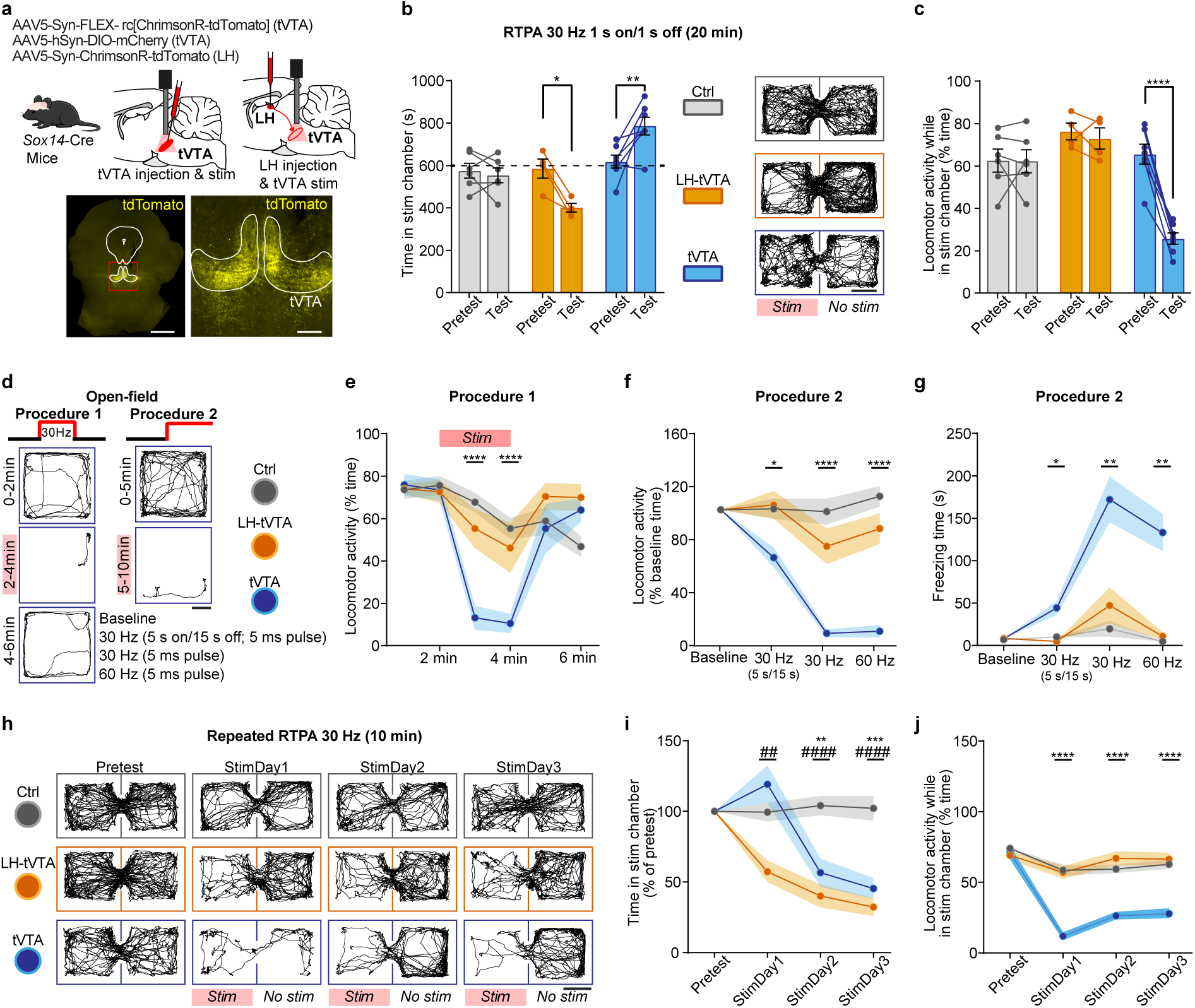
Optogenetic stimulation of *Sox14*+ tVTA neurons inhibits locomotor activity and induces avoidance behavior. **a** Top, schematic of virus delivery for optogenetic stimulation of *Sox14*+ tVTA neurons or of the LH-tVTA pathway. Bottom, representative images of tVTA injection sites. **b** Real-time place avoidance (RTPA). Left, unlike the control group, stimulation of LH-tVTA terminals reduced the time spent in the stimulation chamber, while direct stimulation of *Sox14+* tVTA neurons increased it (Time x Group, F_(2,14)_=14.41, *P*=0.0004). Right, representative tracks on test day. **c** Locomotor activity in the stimulation chamber was unchanged between pretest and test in the control and LH-tVTA groups, but decreased with tVTA stimulation (Time x Group, *F*_(2,14)_=20.75, *P*<0.0001). **d** Open-field procedures. Procedure 1: 30 Hz, 5 ms pulses for 2 min within a 6min test. Procedure 2: daily sessions (5 min off, 5 min on) differing in stimulation frequency and pattern. Tracks from a tVTA animal (3^rd^ day for procedure 2). **e** Stimulation of *Sox14*+ tVTA neurons decreased locomotor activity (Time x Group, *F*_(10,125)_=5.81, *P*<0.0001). **f** Stimulation of *Sox14*+ tVTA neurons induced a stimulation-dependent decrease in locomotor activity (Protocol x Group, *F*_(6,75)_=11.36, *P*<0.0001), associated with increased freezing (**g**; Protocol x Group, *F*_(6,75)_=12.94, *P*<0.0001). **h** Representative tracks over repeated RTPA days for each group. **i** While avoidance with stimulation was immediately present in the LH-tVTA group, it developed over days with stimulation of *Sox14*+ tVTA neurons (Day x Group, *F*_(6,72)_=16.55, *P*<0.0001). **j** On each day, stimulation of *Sox14*+ tVTA neurons decreased locomotor activity in the stimulation chamber (Day x Group, *F*_(6,72)_=11.31, *P*<0.0001). Group sizes (Ctrl, LH-tVTA, tVTA): 6, 4, 7 Sox14-Cre mice in (**b**, **c**); 11, 10, 7 in (**e-g**) and 11, 10, 6 in (**h-j**), with C57BL/6J and Sox14-Cre mice pooled in Ctrl and LH-tVTA groups. Graphs display mean ± s.e.m.; dots: individual animals. Detailed analyses of variance and post-hoc comparisons are provided in **Supplementary Table 11d**. Asterisks: pretest *versus* test in (**b**, **c**), Ctrl *versus* tVTA in (**e-j**); hashes: Ctrl versus LH-tVTA in (i). \**P*<0.05, \*\**P*<0.01, \*\*\**P*<0.001, \*\*\*\**P*<0.0001; same scale for the hashes. Scale bars in **a**: 1 mm (left) and 200 µm (right).

We next explored the consequences of direct optogenetic activation of *Sox14*+ tVTA neurons using Sox14-Cre mice (**Fig.9a**). At 30Hz (5ms pulses, 1s on / 1s off), stimulation of LH-tVTA terminals induced RTPA, whereas direct stimulation of *Sox14*+ tVTA neurons increased the time spent in the stimulation chamber (**Fig.9b**). This unexpected increase was accompanied by a marked decrease in locomotor activity (**Fig.9c**) and speed (**Supplementary Fig.18a**) during stimulation, while these parameters were unaffected in the unstimulated chamber (**Supplementary Fig.18b**, **c**) or by the stimulation of LH-tVTA terminals. This reduction occurred without significant freezing under this protocol (**Supplementary Fig.18d**). These results suggested that direct stimulation of *Sox14*+ tVTA neurons induced a locomotor effect that may mask potential aversive consequences.

To better characterize this locomotor effect, we used open-field procedures (**Fig.9d**). A 2-min stimulation of *Sox14*+ tVTA neurons strongly decreased locomotor activity, an effect immediately reversed when stopping the stimulation (**Fig.9e**). Comparing different protocols, LH-tVTA stimulation did not detectably reduce locomotor activity, whereas direct stimulation of *Sox14*+ tVTA neurons produced a locomotor inhibition that deepened with uninterrupted trains, accompanied by the appearance of freezing (**Fig.9f**, **g**; **Supplementary Fig.18e**, **f**). In a repeated and cued RTPA paradigm, we then tested the effect of daily sessions. Stimulation of *Sox14*+ tVTA neurons led to avoidance of the stimulation chamber on StimDay2 and StimDay3, similar to LH-tVTA stimulation (**Fig.9h**, **i**), while locomotor activity of the tVTA group remained inhibited in that chamber over days (**Fig.9j**). These data supported a dual action of *Sox14*+ tVTA neuron stimulation on locomotor activity and avoidance behavior, with aversive learning favoring the detection of the avoidance component over days.

In an independent experiment, we then tested conditioned avoidance to confirm this aversive learning, and asked whether these effects were mediated by the major projections of *Sox14*+ tVTA neurons to the mesencephalic dopaminergic system. To this end, an optical fiber was implanted above either the tVTA or the VTA/SNc area (**Fig.10a**). Over sessions, stimulation of *Sox14*+ tVTA neurons or of their terminals within the VTA/SNc area produced an increase in the time spent in the stimulation chamber on StimDay1, associated with decreased locomotor activity, followed by the development of avoidance behavior over days (**Fig.10b-d**). The day after the fourth session, a test without stimulation revealed conditioned avoidance in both groups, while locomotor activity had returned to baseline (**Fig.10b**, **c**). An open-field test done a week later then confirmed the immediate and reversible inhibition of locomotor activity with stimulation of either *Sox14*+ tVTA neurons or their terminals within the VTA/SNc area (**Fig.10e**).

**Figure 10.**
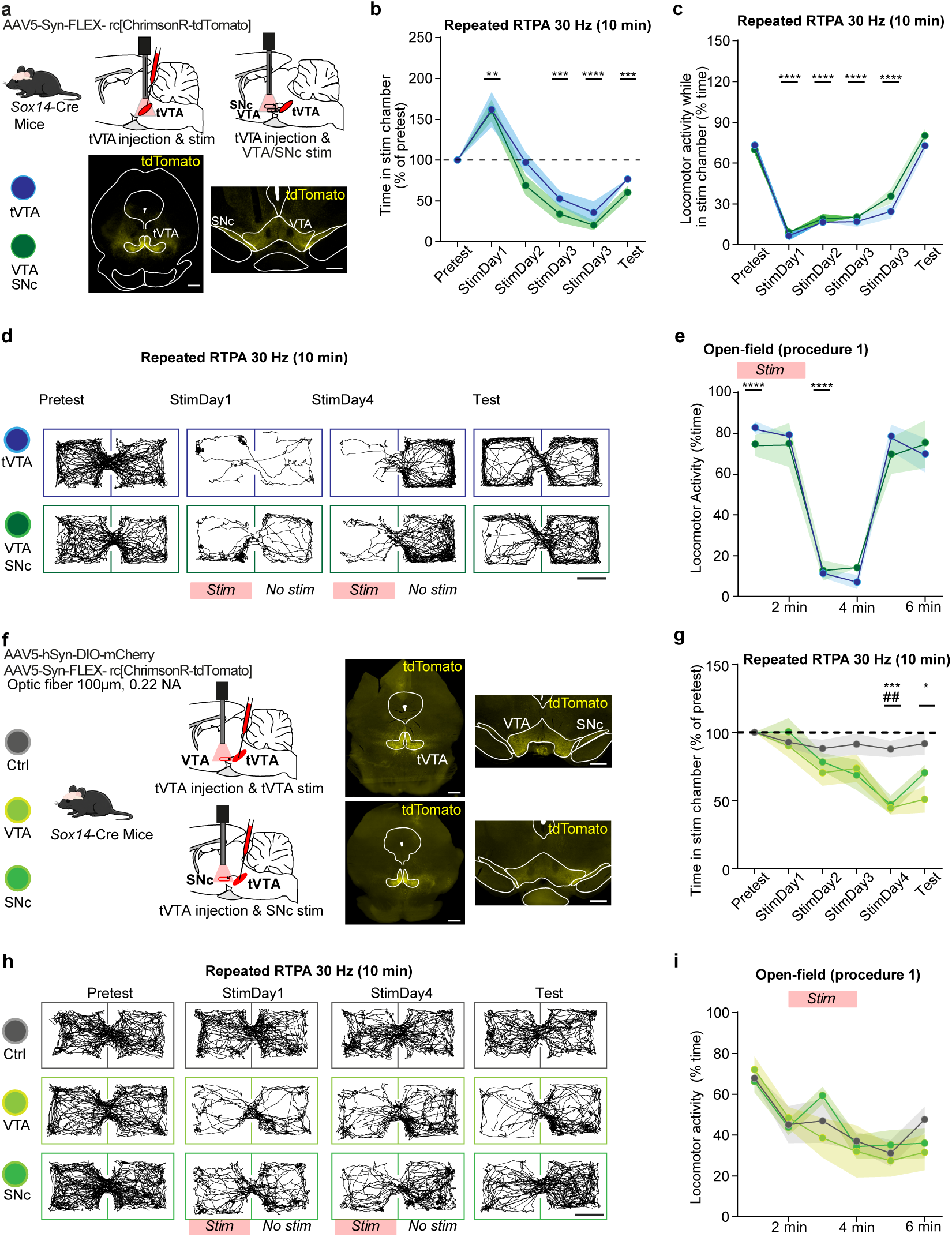
Optogenetic stimulation of *Sox14*+ tVTA neurons or their projections inhibits locomotor activity and favors aversive learning. **a** Left, schematic of virus delivery for optogenetic stimulation of *Sox14*+ tVTA neurons or of their terminals in the VTA/SNc. Right, top, representative image of the tVTA injection site in Sox14-Cre mice; bottom, terminal fibers in the VTA/SNc area. b, c Repeated real-time place avoidance (RTPA). Optogenetic stimulation of *Sox14*+ tVTA neurons and of their VTA/SNc projection (10 min, 30 Hz, 5 ms pulses) had a similar effect on b the time spent in the stimulation chamber (Day effect, *F*_(2.04,14.28)_=37.67, *P*<0.0001) and c the locomotor activity in that chamber (Day effect, *F*_(3.31,23.15)_=138.2, *P*<0.0001), avoidance developing over sessions and persisting on the test day without stimulation. d Representative tracks at pretest, StimDay1, StimDay4 and test, for each group. e Stimulation of *Sox14*+ tVTA neurons and of their VTA/SNc projection similarly decreased locomotor activity in an open-field (Time effect, *F*_(2.35,16.44)_=68.16, *P*<0.0001). f Left, schematic of virus delivery for optogenetic stimulation of terminals in either the VTA or the SNc. Right, representative images of the tVTA injection site in a control and in an opsin-expressing mouse, and of the terminal fields in the VTA and in the SNc. **g** Stimulation of terminals in the VTA or the SNc induced avoidance behavior after repeated sessions (StimDay x Group, *F*_(10,80)_=2.87, *P*=0.0042), and preferentially favored conditioned avoidance after stimulation of terminals in the VTA. **h** Representative tracks at pretest, StimDay1, StimDay4 and test, for each group. **i** Optogenetic stimulation of terminals in the VTA or the SNc did not affect locomotor activity in an open-field (Time x group, F_(10,80)_=0.82, *P*=0.61). Group sizes: tVTA 4 and VTA/SNc 5 in (**b-e**); Ctrl 6, VTA 6, SNc 7 in (**g-i**). Graphs display mean ± s.e.m. Detailed analyses of variance and post-hoc comparisons are provided in **Supplementary Table 11d**. Asterisks: *versus* pretest in (**b**, **c**), across time in (**e**), Ctrl *versus* VTA in (**g**); hashes: Ctrl *versus* SNc in (**g**). \**P*<0.05, \*\**P*<0.01, \*\*\**P*<0.001, \*\*\*\**P*<0.0001; same scale for the hashes. Scale bar in **a** and **f**: 1 mm (wide views) and 500 µm (magnifications).

To dissect the respective contributions of the VTA and SNc terminal fields of *Sox14*+ tVTA neurons, we used smaller injection volumes (66nL) and thinner optical fibers (100µm) implanted above either the VTA or the SNc, restricting the recruitment of tVTA projections (**Fig.10f**). The increased time in the stimulation chamber was no longer present on StimDay1 but avoidance behavior developed over days (**Fig.10g**, **h**), without change in freezing across sessions (**Supplementary Table11d**). RTPA developed whether the optical fiber was inserted above the VTA or the SNc, conditioned avoidance reaching significance against control only for VTA terminal stimulation. In the open-field test, terminal stimulation did not significantly affect locomotor activity (**Fig.10i**). Restricting recruitment therefore dissociated the two components of the behavior, avoidance persisting while locomotor activity was no longer affected.

## DISCUSSION

Over the years, the tVTA/RMTg has been shown to be critical in controlling dopaminergic systems and implicated in various behavioral functions, but 20 years after its discovery this brain structure is still not present in brain atlases. This absence, partly explained by the lack of detailed definition across species, is however limiting functional studies. Here we demonstrate that *Sox14*, expressed by most of the tVTA GABAergic neurons in the rodent brain, provides a molecular signature of the structure, allows to define its boundaries, and offers a genetic access to a nucleus that was previously difficult to selectively target using stereotaxic coordinates. Using this access to selectively map the whole-brain connectivity of *Sox14*+ tVTA GABAergic neurons, we show that inputs reach this discrete neuronal population from the whole neuraxis, while outputs of these *Sox14* neurons funneled onto the adjacent dopaminergic midbrain, thus recapitulating the connectivity of the structure itself. Driving these *Sox14* neurons controls dopamine neuron activity, accumbal dopamine levels, and motor and avoidance behavior.

*Sox14* is a transcription factor preserved in vertebrates across evolution, and that has been described in the embryonic mesencephalic GABAergic lineage leading to tVTA neurons^32^. Using classical *in situ* hybridization, its expression has however been initially reported as not detected in the adult tVTA neurons in mice^32^, likely for sensitivity reasons. Indeed, the bulk RNAseq analysis of transcripts enriched in the adult rat tVTA highlights its presence at adulthood, which we also confirmed in both rats and mice by using the RNAscope approach that can resolve individual transcripts, and with spatial transcriptomics analysis in mice.

From the spatial transcriptomics experiment, we did not identify a combination of two or three genes of the studied panel that was better, in terms of both coverage and selectivity, than *Sox14* to highlight the tVTA territory. Two rarer genes, *Tmem132c* and *Sema3e*, were more selective than *Sox14* within the panel, but covered fewer tVTA neurons. Conversely, *Pnoc* was detected in most *Sox14*+ tVTA neurons, but is also present in nearby areas, thus carrying limited contrast locally as also shown using the Pnoc-Cre transgenic line^31^. The FOXP1 protein, however, has been shown to mark the tVTA^31^ and it remains a tool of choice to visualize it on a slide; yet its transcript is largely present in the brain, including at mesopontine level, making unlikely a Foxp1-Cre line to restrict recombination to the targeted nucleus. While *Sox14* allowed to locally define and characterize the tVTA, its expression is not restricted either to this structure nor to GABAergic neurons. Indeed, the Allen whole mouse brain atlas^35^ show that *Sox14*-expressing cell types span the brainstem, cerebellum and diencephalon (but not the telencephalon), and that the tVTA represents only a part of those cells. Accordingly, on the Xenium sections, *Sox14* also labeled a glutamatergic population of the IPN^32^ and GABAergic neurons of the superior colliculus^34^. Its value as a marker thus lies in the local contrast of its expression within the ventral midbrain, not in brain-wide specificity. Nevertheless, this distinguishes Sox14-Cre mice from the pan-GABAergic transgenic lines used so far to reach the nucleus^46,47^, in which selectivity came from injection placement only rather than from gene-related markers of the structure. Overall, *Sox14* marks a developmental family and, by its position and by the combination of its clusters, separates the tVTA from most of the neighboring GABAergic populations compared. It thus allows proposing an anatomo-molecular definition for the structure.

The processed tVTA Xenium sections all came from a single rostro-caudal level. Merging them increased the number of cells analyzed, but the direct description remains limited to that level. Mapping the cellular clusters onto the Allen transcriptomic and spatial atlas of the mouse brain^35^ then extended, through data mining, the description to the whole structure. The territory traced this way displayed a shape similar to the one reconstructed from chromogenic *in situ* hybridization. Presently, the Allen common coordinate framework gives however no name to this position. The subclass that holds these clusters, CS-PRNr-DR En1 Sox2 Gaba, is named after two genes and after the raphe and reticular territories its cells were dissected from. Therefore, neurons of the tVTA were already present in the reference datasets, but under borrowed names. They were given no name of their own because the reticular formation has no distinct cytoarchitectural boundaries, and the nuclei embedded in it are delineated with difficulty on histological criteria alone^48^.

The tVTA clusters we identified are topographically ordered but intermingled, and their markers highlight neuropeptide signaling. The tVTA as a whole expresses the mu-opioid receptor^2^, but the dorsal summit is enriched in *Npffr2* and expresses *Pth2r*, respectively coding neuropeptide FF and TIP39 receptors. Interestingly, a NPFF2 agonist in the VTA can blunt the accumbal dopamine response to morphine^49^, and TIP39 has been implicated in nociceptive processing and affective behavior^50^, thus pointing to functions also related to the tVTA, the modulation of pain and opioid analgesia^23^ and opioid control of dopamine activity^13–15^. In link with such pain-related functions, the tVTA receives inputs from the parabrachial nucleus, medullary raphe and PAG. The medial column highlighted a different system, with the Allen datasets also pointing to *Egflam*, coding an extracellular matrix-like protein, as a marker. This medial column is enriched in *Rxfp3*, coding a relaxin-3 receptor, a peptide produced by the nucleus incertus^51^ that is also among inputs to *Sox14*+ tVTA neurons. This peptide contributes to arousal, stress responses, feeding and memory^51^, and the tVTA is itself known for regulating sleep^26,27^. The medial column could therefore be part of an ascending pathway contributing to arousal. However, none of these local markers is selective enough to offer genetic access to a single tVTA compartment.

A feature of tVTA neurons is the inhibition they impose on the dopamine neurons they contact, which has been established for more than a decade, in slices and in the intact brain^1,2,8,11–14^. This drive differs, in its short-term plasticity, from the one exerted by GABAergic interneurons intrinsic to the VTA^52^. Indeed, the drive from VTA interneurons loses strength during a train, while that from tVTA neurons is maintained. Interestingly, these characteristics, together with being mediated by GABA_A_^13,53^ and under presynaptic mu-opioid receptor influence^13,54^, also characterize *Sox14+* tVTA inputs to dopamine neurons. Together with the fact that *Sox14* is poorly detected in the VTA or SNc^35^, it clearly supports that VTA interneurons and *Sox14*+ tVTA ones are separate populations. In the intact brain, chemogenetic activation of tVTA GABAergic neurons has been shown to lower accumbal dopamine^23^. By using optogenetic, the present study demonstrates that driving *Sox14*+ tVTA neurons is sufficient *per se* to rapidly decrease accumbal dopamine, with a duration that depends on the stimulation frequency and lasts beyond stimulation end.

Aversion has been the most consistent behavioral consequence of driving tVTA neurons. It is induced by stimulating LH^9^ or dorsomedial prefrontal cortex^55^ afferents to the structure, as well as its terminals in the VTA^56^ and the SNc^22^. Locomotion, however, did not always display similar data. Pharmacological inactivation, ablation and chemogenetic activation of the tVTA itself can change motor output^12,20,23,47^, whereas the pathway-restricted manipulations that produce avoidance left general locomotion unchanged when it was measured^9,22,56^. These results may relate to the amount of affected tVTA neurons, and for how long. Indeed, each manipulation that altered motor output was permanent or sustained, and none of them was restricted to a single anatomical pathway. Our data support that reading. Stimulation of the habenular input to the tVTA reproduced the published pattern^9^, place avoidance with no change in locomotion. Driving the *Sox14*+ neurons themselves, over entire tVTA, produced both effects, a marked locomotor inhibition and avoidance over days of the stimulation chamber. Driving largely their midbrain terminals did the same, whereas restricting the recruitment to *Sox14*+ terminals in either the VTA or SNc spared the avoidance and did not affect locomotion. How much of the *Sox14*+ terminal field was illuminated thus appears important to the immediate impact on locomotion, while more discrete and repeated stimulations are enough to drive the acquisition of avoidance.

In rats, the tVTA connectome has been mapped^2,4,8^, with descriptive rather than quantified reporting. More recently, a quantitative map of the mouse tVTA afferents was produced, with 71 input regions identified using Vgat-Cre animals^45^. There is, however, no equivalent precedent for a quantitative map of tVTA efferents in mice; the only available description in this species being qualitative, secondary to a study on sleep^27^. A main add-on of the afferent/efferent connectome we provided here is to use a genetically defined starting population, highlighting the 84 input and 110 output regions of *Sox14*+ tVTA neurons. Qualitatively, the regions it highlights are similar to those already described for the whole structure in rats and mice. The afferent rankings of *Vgat* and *Sox14*-based maps disagree, in part because the study in Vgat-Cre mice ranked regions by the proportion of labeled neurons while we used a density, which favors small and compact regions^45^. Analysis by whole regions can also carry biases, since a locally dense cluster of labeling may be diluted in a larger region, as shown by the contralateral input from a subdivision of the superior colliculus. Both maps rest on a small number of animals. While there is agreement between animals, the rank of the regions connected to *Sox14*+ tVTA neurons remains qualitative. Our data show that inputs to *Sox14*+ tVTA neurons originate from the whole neuraxis, whereas outputs are concentrated in the midbrain immediately rostral to the tVTA, the VTA complex and the SNc accounting for 49.0% of the efferents. This funnel effect provides anatomical evidence for the view of the tVTA as main GABA brake on dopamine systems^5,6^. Raphe nuclei are densely connected to the tVTA in both directions, with the dorsal raphe ranking first among afferent and third among efferent regions, a projection also described in rats^2,57^. This influence is further supported by the 5-HT2C receptor transcript, detected in 48% of the *Sox14*+ tVTA neurons (*Htr2c*; **Supplementary Table 8d**), which depolarizes tVTA neurons in the rat^58^.

*Sox14* does not label every GABAergic neuron of the tVTA, yet those it labels reproduce the connectivity, the synaptic signature and the behavioral control that define the structure itself. Working on them is working on the tVTA. A nucleus that could until now only be approximately targeted can be now anatomo-molecularly defined, and recognized in the reference datasets where its neurons sat under borrowed names. The Sox14-Cre line, the quantitative connectome and the molecular and spatial definition of the territory give other groups a basis to work from, and turn the question away from where this nucleus lies toward what each of its parts does.

## METHODS

### Animals

Male Sprague Dawley rats (Rj:SD; Janvier Labs, France) were used for the transcriptomic (280 to 350 g) and RNAscope (200 to 250 g) experiments, and male C57BL/6J mice (RRID:IMSR_JAX:000664; Charles River, France) for the RNAscope (8 to 12 weeks, 20 to 35 g), Xenium (3 months) and behavioral (2 months) experiments. Heterozygous Sox14-Cre mice (Sox14<tm2.1(cre)Tmj>, MGI:5909921) on a C57BL/6J background were used for the *ex vivo* and *in vivo* recordings (3 to 10 months, over 20 g, both sexes), the connectome (3 to 6 months, over 20 g, both sexes) and the behavioral experiments (2 to 6 months, 20 to 40 g); the colony was bred in-house from breeders (Charles River, France) derived from sperm straws (gift from Prof. A. Delogu, King’s College London, UK)^34^, and every mouse was genotyped before inclusion. Animals were housed at the Chronobiotron facility (UAR3415, Strasbourg; agreement E67211838) under standard conditions: group-housed, 22 to 24°C, about 50% relative humidity, reversed 12 h light/dark cycle (lights off at 08:00), with environmental enrichment (wooden sticks, cotton squares and shredded nesting material) and food and water available *ad libitum*. All procedures followed European Directive 2010/63/EU, were approved by the regional ethics committee (CREMEAS) and authorized by the French Ministry of Higher Education, Research and Innovation under APAFIS #03691.02, #15234-2018052510531979, #37334-2022051213209078, #42625-2023041211468830 and #49994-2024062419221611. The RNAscope and Xenium experiments used tissue collected after deep anesthesia and killing, without any prior procedure, and therefore did not require project authorization (Directive 2010/63/EU, Article 3).

### Mouse stereotaxic surgery

Surgeries were performed under isoflurane anesthesia (Vetflurane®, Virbac, Carros, France; induction 4%, maintenance 1.5 to 2%, 0.5 L/min). After disinfection of the scalp with 0.5% chlorhexidine and 70% ethanol, a local anesthetic was delivered at the incision site (bupivacaine/lidocaine, 2 mg/kg each, 20 µL, subcutaneously (s.c.)) 5 min before the incision. Body temperature was maintained at 37°C with a thermostatically controlled heating pad, and eyes were protected with ophthalmic ointment (Ocry-gel®, Centravet,). Mice were placed in a stereotaxic frame (Kopf Instruments), the skull was exposed from lambda to bregma and leveled, and a circular craniotomy was drilled above each target. At the end of surgery, the scalp was sutured (Ethilon 5-0) and disinfected with 0.5% chlorhexidine, except in mice implanted with an optic fiber, where the headcap closed the incision. Meloxicam was administered (Meloxicam®, Biogaran, 10 mg/kg s.c., 50 µL/10 g), and mice recovered in a separate cage until fully awake before returning to their home cage. Analgesia was continued for 48 h (Inflacam®, Virbac, oral suspension, 10 mg/kg), and animals were monitored daily for at least 3 days until full recovery.

Intracerebral viral injections were performed at a rate of 66 nL/min using glass pipettes (30-40 µm tip diameter, Hirschmann®, Ringcaps® #9600105) connected to a manual pressure injector. Pipette lowering and retraction were controlled by a motorized manipulator (Scientifica, 1U RACK model) and monitored via a digital interface (LinLab2, v2.7.4, Scientifica). Pipettes were held in place for 10 min before and after each injection.

Animals for behavioral experiments underwent optic fiber implantation within the same surgical procedure. A stainless-steel screw (1.6 mm diameter) was secured to the skull contralateral to the fiber (anteroposterior (AP) -2 mm, mediolateral (ML) ±2 mm) as an anchor for the dental cement, and the optic fiber was lowered with the motorized manipulator and positioned 200 µm above the targeted structure. A first layer of adhesive dental cement (Super-Bond® Universal Kit, Sun Medical Co., Ltd.) was applied around the cannula base and anchor screw, and the headcap was then reinforced with dental acrylic resin (Paladur®, Kulzer, Mitsui Chemicals Group). A layer of opaque varnish was finally applied to block light transmission through the headcap.

Viral vectors, optic fibers and stereotaxic coordinates are given in each experimental section, AP ML in mm from bregma and dorsoventral (DV) in mm from dura. All injections were made in the right hemisphere.

### Transcardial perfusion

Animals were deeply anesthetized with a mixture of ketamine (Ketamine 1000®, Virbac, 300 mg/kg) and xylazine (Paxman®, Virbac, 20 mg/kg), injected intraperitoneally (i.p.), and received lidocaine (2 mg/kg, s.c., at cage rib level) 2 min before the opening. They were then transcardially perfused with 0.1 M phosphate buffer (PB, pH 7.4) for 5 min, followed by 0.1 M PB with 4% paraformaldehyde (PFA) for 10 min. Brains were removed and post-fixed overnight at 4°C in 0.1 M PB with 4% PFA.

### Transcriptomic analysis

#### Surgery and tVTA laser capture microdissection

Rats were anesthetized intraperitoneally with ketamine, xylazine and acepromazine (80, 4 and 1 mg/kg) and placed in a stereotaxic frame (Stoelting, USA). Red fluorescent retrobeads (Red RetroBeads IX, FP-DX2100; Interchim, France) were pressure-injected bilaterally into the VTA, 150 nL *per* side, at AP -5.2, ML ±0.7 and DV 6.7 and 7.5^28^. One week later, rats received an overdose of pentobarbital (800 mg/kg, i.p.; Dolethal, Vetoquinol, France), and brains were removed, frozen in isopentane (-30 to -40°C) and stored at -80°C. Coronal sections of 100 µm containing the tVTA were cut on a cryostat (Microm HM550, Thermo Fisher Scientific, France), stained with cresyl violet (0.1% w/v in 70% ethanol, 2 min), dehydrated in graded ethanol (2 × 2 min at 75%, 95% and 100%), air-dried and vacuum-desiccated for 60 min. Microdissection was performed on an Arcturus Veritas system. The tVTA was identified at ×2 magnification from the retrograde bead labeling under fluorescence and dissected at ×10 with the capture-and-cut procedure (infrared laser 70 to 100 mW, 2500 to 5000 µs pulse, 1 to 3 hits, CapSure Macro LCM caps LCM0211 on PEN membrane frame slides LCM0521; ultraviolet laser 22.13 mW, 0.5 µm spot). Captured samples were collected in 1.5 mL tubes and stored at -80°C. The whole-brain samples came from three naive rats, which received the overdose alone and whose brains were homogenized with a Polytron.

#### mRNA-seq

Total RNA was extracted from the microdissected tVTA samples and from the whole-brain homogenates with the RNeasy Micro Kit (74004, Qiagen, Germany) according to the manufacturer’s instructions. Libraries were prepared from three tVTA samples and the three rat brains at the GenomEast platform (IGBMC, Illkirch, France). Full-length cDNA was generated from 5 ng of total RNA with the SMART-Seq v4 Ultra Low Input RNA Kit for Sequencing (634890, Clontech) using 9 cycles of amplification with Seq-Amp polymerase. Six hundred ng of pre-amplified cDNA were tagmented with the Nextera XT kit (FC-131-1096, Illumina), followed by 12 cycles of library amplification. Libraries were purified with Agencourt AMPure XP beads (A63882, Beckman Coulter) and their size and concentration assessed on an Agilent 2100 Bioanalyzer. Libraries were sequenced (Illumina HiSeq 4000) as 100 bp paired-end reads.

Reads were preprocessed using cutadapt 4.2^59^ in order to remove adapter, polyA and low-quality sequences (Phred quality score < 20). For further analysis, reads shorter than 40 bases were also discarded, as well as reads that were mapped to rRNA sequences of RefSeq and GenBank databases using bowtie 2 (2.3.5)^60^. Then, reads were mapped onto the GRCr8 assembly of *Rattus Norvegicus* genome using STAR 2.7.10b^61^. Gene expression quantification was performed from uniquely aligned reads using HTSeq 1.99.2^62^, with annotations from Ensembl v115 under union mode. Only non-ambiguously assigned reads to a gene were retained for further analyses. Read counts were normalized across samples using the median-of-ratios method^63^ to ensure comparability between samples. Differential expression comparisons were performed using the Wald test for count data, with *P*-value adjustment via the Benjamini-Hochberg method^64^. This approach was implemented in the DESeq2 Bioconductor package (1.50.2)^65^.

#### Downstream analysis

Analyses started from the gene table delivered by the platform, which already carried the log_2_ fold changes of the tVTA against whole brain, shrunken with the apeglm method^66^, and the adjusted *P* values. 24,393 of the 32,643 genes received an adjusted *P* after independent filtering. Enriched genes had an adjusted *P* < 0.05 and a shrunken log_2_ fold change > 1; the set used in **Fig. 1c** and in the Xenium analysis was restricted to the 863 protein-coding genes detected in all three tVTA libraries with a mean of at least 5 transcripts *per* million (TPM). The principal component analysis used the variance-stabilized counts^63^ of the 32,643 genes with at least one read, recomputed from the raw counts. Monoaminergic and oligodendrocyte markers were determined from the literature (**Supplementary Table 1**), and the myelin content of a library is the proportion of the eleven oligodendrocyte genes in its total TPM.

### RNAscope *in situ* hybridization (ISH)

For fluorescent and chromogenic RNAscope, animals deeply anesthetized with isoflurane (4%, 5 min, 0.5 L/min) were decapitated. Brains were rapidly frozen in isopentane (-30 to -40°C), stored at -80°C and cut on a cryostat (CM3050, Leica, Germany) into 14 µm coronal sections through the VTA and tVTA, thaw-mounted on SuperFrost slides (Epredia, France) and stored at -80°C. For combined RNAscope and immunohistochemistry, animals were perfused as described above, and brains were cryoprotected in graded sucrose (10/20/30%), frozen in isopentane and stored at -80°C. Sections were cut at 30 µm, kept in cryoprotectant solution at -20°C and mounted on SuperFrost slides before hybridization. All sections were post-fixed in 4% PFA, dehydrated in graded ethanol (50/70/100/100%, 5 min each), air-dried at room temperature (RT, 5 min), delimited with a hydrophobic pen (ImmEdge; Advanced Cell Diagnostics, ACD, France), pretreated with H₂O₂ (#322335, ACD; 10 min, RT) and then Protease IV (#322336, ACD; 30 min, RT), and rinsed in PB saline (PBS). Post-fixation lasted 1 h 30 min at 4°C for duplex chromogenic ISH and 15 min for multiplex fluorescent ISH. The RNAscope probes targeting *Sox14*, *Gad1*, *Oprm1*, *Th*, *Gad2*, *Sstr1*, *Pnoc* and *Foxp1* (ACD) and the antibodies are detailed in **Supplementary Tables 3 and 4**.

#### Chromogenic RNAscope and 3D reconstruction

Duplex chromogenic ISH was performed with the RNAscope 2.5 HD Duplex Detection Kit (#322500; ACD) in a hybridization oven (#321710, ACD). Undiluted *Sox14* probe was combined with *Gad1* diluted 1:50 in the C1 (*Sox14*) probe, and hybridized for 2 h at 40°C. After amplification (AMP1-AMP10), the C1 (*Sox14*) signal was developed as a blue precipitate (horseradish peroxidase, after AMP6) and the C2 (*Gad1*) signal as a red precipitate (alkaline phosphatase, after AMP10). Sections were counterstained with hematoxylin, dried at 60°C for 15 min, cleared in Roti-Histol and mounted (VectaMount). Chromogenic ISH was sampled every 4^th^ section in mouse and every 8^th^ section in rat (*n* = 1 animal *per* species). Whole-slide brightfield images were acquired on a NanoZoomer® S60 scanner (Hamamatsu; NDP Scan v3.4.4; ×40, 0.23 µm/pixel) and imported into Neurolucida (v2020, MBF Bioscience) for 3D reconstruction, in which the contours of the PAG, cerebral aqueduct (Aq), IPN, VTA and SNc were drawn on each serial section, *Sox14* and *Gad1* labeled neurons were manually plotted, serial sections were aligned and the tVTA volume was reconstructed. A cell was scored positive when at least three dots lay at or near its nucleus.

#### Fluorescent RNAscope

Multiplex fluorescent ISH was performed with the RNAscope Multiplex Fluorescent Reagent Kit v2 (#323110, ACD), with ACD wash buffer (#310091) between steps. Positive and negative (*dapB*) control probes were run on adjacent sections. Probes were hybridized 2 h at 40°C, amplified (AMP1-AMP3) and developed channel by channel (HRP, tyramide-coupled fluorophore 1:1,500, HRP-blocker between channels). Signal was revealed with Opal 520/570/690 dyes (#FP1487001KT for 520, #FP1488001KT for 570, #FP1497001KT for 690; Akoya Biosciences) for the *Sox14*/*Gad1*, *Sox14*/*Th*/*Oprm1* and *Gad1*/*Oprm1* experiments, and with tyramide signal amplification 520/570/650 (TSA) fluorophores (#323271 for 520, #323272 for 570, #323273 for 650; Bio-Techne) for the *Gad2*, *Sstr1*, *Pnoc* and *Foxp1*-transcript experiments; *Sox14* was consistently assigned to the 570 dyes. Nuclei were counterstained with DAPI 0.5 µg/mL (MBD0015, Roche®) for 30 s at RT and sections mounted in ProLong Gold (#P36930, Invitrogen).

#### Combined RNAscope (ISH) and immunohistochemistry (IHC)

Transcript detection of *Sox14* was combined with protein detection of NeuN (*i.e.* RBFOX3) or FOXP1. For *Sox14*/NeuN, IHC followed ISH. Sections were blocked (PBS + 0.2% Triton X-100 + 2% normal donkey serum, 1 h, RT), incubated overnight at 4°C with anti-NeuN (rabbit, 1:1,000) in PBS-Triton X-100 + 1% donkey serum, then 2 h at RT with donkey anti-rabbit Alexa Fluor 488 (1:400). For *Sox14*/FOXP1, IHC preceded ISH. Sections were blocked (PBS 0.1 M + 0.3% Triton X-100 + 5% normal donkey serum, 1 h), incubated overnight with anti-FOXP1 (rabbit, 1:500) in PBS + 0.1% Triton X-100 + 1% donkey serum, slide-mounted, re-fixed in 4% PFA (30 min), processed for fluorescent ISH, then incubated with donkey anti-rabbit Alexa Fluor 647 (1:400). Sections were then counterstained and mounted as above.

#### Image acquisition and quantification

Fluorescent sections were imaged on a Leica SP5 II confocal microscope (×20 objective, ×2 digital zoom; LAS AF software; 1,024 × 1,024 pixels; 387 × 387 µm field; 0.38 µm/pixel; 3× line averaging; 400 Hz) as z-stacks acquired every 1 µm, with two fields (one *per* hemisphere) *per* section; excitation lines were 405 nm (DAPI), 488 nm (520 dye), 561 nm (570 dye) and 633 nm (690 and 650 dye). AP levels were referenced to bregma according to mouse^29^ and rat^28^ atlases. Chromogenic labeling and 3D reconstruction spanned the full rostro-caudal extent of the tVTA, whereas fluorescent quantification was restricted to mid rostro-caudal levels (mouse -3.9 to -4.1 mm, 200 µm; rat -6.6 to -7.2 mm, 600 µm). Confocal stacks were flattened by maximum-intensity projection over 5 µm in Fiji. Marker channels were normalized by pixel classification in Ilastik 1.3.3 (trained on at least five representative images *per* marker); nuclei were segmented on the DAPI channel in QuPath 0.4.3 (pixel size 0.5 µm, background radius 8 µm, sigma 1.5 µm, area 10-400 µm², intensity threshold 25, cell expansion 8 µm, split by shape). A cell was scored positive for a transcript when its signal exceeded a batch-constant intensity threshold validated by visual inspection, with manual correction of false detections, and co-expression was defined as positivity for both markers within the same nucleus. The section was the sampling unit, pooling data from its two hemispheres. Percentage of single- and double-labeled cells (summing to 100%) gave one data point *per* section for the violin plots. Pie charts show co-expression pooled across sections, in both directions (double-labeled cells over all cells positive for the reference marker, for example *Sox14* among *Gad1*+ and the reverse). Pooled proportions and *per*-section means therefore differ slightly. Values are given as mean ± s.e.m. across sections in the figures and across animals in the Results, the value of an animal being computed on the pooled counts of its sections. Violin plots and pie charts were drawn in GraphPad Prism 8.4, and displayed images were processed in Fiji/ImageJ (v1.54g) and Adobe Photoshop (v26.10), with colors adjusted for color-blind readers.

### Xenium spatial transcriptomics

#### Tissue and Xenium run

Three mice were deeply anesthetized with isoflurane (4%, 5 min, 0.5 L/min) and decapitated, and brains were frozen in isopentane at -30°C and stored at -80°C. Six coronal sections (10 µm; Leica CM3050 cryostat) were placed on one Xenium slide (10x Genomics), five through the tVTA (two from each of mice 1 and 2, one from mouse 3) and one at the level of the VTA (mouse 3). The slide was processed at the PHEN-ICS platform (Illkirch, France) following the manufacturer’s protocols for fresh frozen tissue (CG000581 Rev D and CG000760 Rev C), with probe hybridization for 17 h at 50°C and cell segmentation staining. Probes targeted the Xenium Mouse 5K Pan Tissue & Pathways Panel (5,006 genes, *Sox14* included) and 100 custom genes (**Supplementary Table 5**). The slide was imaged on a Xenium Analyzer at the GenomEast platform (CG000584 Rev G), and transcripts were decoded and cells segmented with Xenium Onboard Analysis 3.3.0.1.

#### Preprocessing and cell classes

Cell filtering and clustering were run on the six sections; all subsequent analyses used the five sections through the tVTA. Cells with fewer than 10 transcripts, fewer than 5 genes or an area below 10 µm² were removed, leaving 435,402 cells (median of 856 transcripts and 551 genes *per* cell). Counts were scaled to the median total count *per* cell, log-transformed and standardized, and cells were clustered with the Leiden algorithm on 50 principal components (15 neighbors, resolution 1.0; scanpy^67,68^). Clusters were grouped into 14 cell classes, four of them neuronal (glutamatergic, GABAergic, serotonergic and dopaminergic), from canonical marker scores and their most differential genes. A cell expressed a gene when its count reached a detection threshold set for that gene, the smallest count whose probability under a Poisson background (negative control codewords and non-neuronal classes not expressing the gene) was below 0.05. Every analysis was repeated with a more stringent threshold (one then two counts for *Sox14*), and a result was kept only when it held under both. All cells were mapped onto the Allen whole mouse brain taxonomy (10x Whole Mouse Brain, CCN20230722^35^) with MapMyCells (hierarchical mapping, cell_type_mapper 1.7.4), which assigned each cell a class, subclass, supertype and cluster.

#### Delineation of the tVTA territory

The five sections were registered in a common frame built on their serotonergic neurons (midline through the dorsal and median raphe nuclei, origin at the median raphe), and the two hemispheres were folded onto each other. Route A scored each GABAergic neuron with the 863 enriched genes of the rat bulk screen (see above) that were in the panel and detected in at least 1% of the GABAergic neurons of each mouse (167 genes, without contamination, glial or monoaminergic genes, and without *Sox14*), as the mean of their z-scores weighted by the bulk fold change. Route B subclustered the GABAergic neurons above the 10^th^ percentile of transcripts (Leiden, resolution 1.0) and retained the subgroup with the highest detection of *Oprm1*, *Pnoc* and *Foxp1*. For each route, the local relief was the difference between a fine (75 µm) and a broad (400 µm) Gaussian smoothing of these values on a 25 µm grid (nodes with at least 15 neurons), and the territory was the region around its peak that stayed above 0.40 of the peak height over the background (minimum 0.02 mm²). The final territory was traced in the same way on the average of the two rescaled reliefs, and the DR, traced likewise from serotonergic neurons, was then removed from it. Overlaps were measured with the Dice coefficient. As controls, the procedure was applied to the GABAergic neurons expressing *Sox14*, to the four clusters of Allen subclass 268 with more than 25% of their cells in the region common to both routes (route C), and to 13 unrelated Allen clusters. The territory was also redrawn after leaving out each mouse or section and after varying each setting (**Supplementary Table 6**).

#### Analysis of the territory

A UMAP was computed on the cells of the final territory alone. Its GABAergic neurons were split into a dorsal compartment and ventral wings at 812 µm above the median raphe (Otsu’s method on their height^69^), and the wings into medial and lateral at their median distance from the midline (364 µm). The coverage of a gene was the proportion of the GABAergic neurons of the territory expressing it, and its selectivity was the ratio of this proportion to its mean in five 100 µm rings around the territory (geometric mean of the three mice), compared with 1,000 random placements of the territory. The Allen clusters of the tVTA were the clusters of subclass 268 with at least a quarter of their GABAergic neurons inside the territory; the zone of highest concentration of a cluster was where its local proportion among GABAergic neurons exceeded its median in the territory by 0.40 of the difference to its peak. As *Sox14* is one of the marker genes used by MapMyCells, the cells of the territory were also mapped without *Sox14*. A gene differed between two groups of neurons when the ratio of its proportions went the same way in every mouse, reached at least 2 as a geometric mean, and the gene was expressed by at least 20% of one group. A local marker of a cluster was considered is metting this criterion against the other GABAergic neurons of the territory, the three other clusters and each neighboring population, and was twice as frequent in the territory as in the 500 µm around it. The ten neighboring populations were drawn like the territory, from the density of their defining cells (**Supplementary Fig. 6d**).

#### Allen single-cell and MERFISH data

Local markers were also sought in the Allen whole-brain single-cell data (10x v3) and MERFISH atlas (brain C57BL6J-638850^35^), mapped to the same taxonomy and to the Allen CCF^30^. A gene was considered detected from 1 UMI in the single-cell data and 2 counts in the MERFISH atlas; a marker of a cluster was considered detected if present in at least 20% of its cells and at least twice as often as in every other tVTA cluster and neighboring cell group. On MERFISH sections, the tVTA territory was traced from the density of the cells of the six clusters (250 µm minus 400 µm Gaussian kernel, 25 µm grid, contour at 0.30 of the peak), without the serotonergic nuclei and the IPN. Each section was placed on its own plane in the CCF, and the volume was interpolated between sections.

#### Statistics and software

Three mice were used and comparisons were descriptive, without hypothesis testing. Unless stated otherwise, proportions pooled the three mice and ratios were geometric means of the *per*-mouse ratios; for selectivity and gene comparisons, proportions were the mean of the three mice.

### Patch-clamp recordings

#### Slice preparation and recordings

Mice received a stereotaxic injection of 180 nL of AAV5-Syn-FLEX-rc[ChrimsonR-tdTomato] (Addgene, #62723-AAV5; titer ≥ 5 x 10¹² vg/mL) in the right tVTA (AP -4.0, ML +0.8, DV -4.2, with a 5° lateral approach). Recordings were made in 250 µm thick slices, coronal, sagittal or horizontal for the tVTA (14 mice, 4 to 7 weeks after injection) and horizontal for the VTA (20 mice, 2 to 5 weeks after injection). Mice were killed by cervical dislocation and the brain was rapidly removed. Slices were cut on a vibratome (HM650V, Microm) in a solution containing (in mM) choline chloride 110, NaHCO₃ 26, KCl 2.5, NaH₂PO₄ 1.25, glucose 25, Na-ascorbate 11.6, Na-pyruvate 3.1, thiourea 2, N-acetyl-L-cysteine 12, kynurenic acid 2, MgSO₄·7H₂O 10 and CaCl₂ 0.5. Slices recovered for 30 min at 32°C and 15 min at room temperature in artificial cerebrospinal fluid (aCSF; in mM NaCl 126, NaHCO₃ 21.4, KCl 2.5, NaH₂PO₄ 1.25, glucose 11.1, MgCl₂·6H₂O 1.2, CaCl₂ 2.4) supplemented with Na-ascorbate 1, N-acetyl-L-cysteine 6, Na-pyruvate 1 and kynurenic acid 2, and were then kept at room temperature in aCSF until use. All solutions were continuously bubbled with carbogen (95% O₂, 5% CO₂). Slices were transferred to a recording chamber perfused at 2 mL/min with oxygenated aCSF at 32 to 34°C, on an upright microscope (Slicescope, Scientifica) under infrared oblique contrast illumination. Recordings were made with a Multiclamp 700B amplifier (Molecular Devices) and the AxographX software at 20 kHz, using 3 to 4 MΩ glass pipettes. For loose cell-attached (LCA) recordings, the pipette was filled with aCSF, positioned on the neuron, and a light suction was applied to reach a seal of 15 to 30 MΩ, the amplifier being used in voltage-clamp mode. Whole-cell pipettes were filled with (in mM) K-gluconate 135, MgCl₂ 2, CaCl₂ 0.025, EGTA 1, Na-ATP 4, Na-GTP 0.5, HEPES 10, Atto 488 0.02 (Atto-Tec) and neurobiotin 1 mg/mL (Vector Labs); pH 7.2, 269 mOsm/L. Membrane resistance and capacitance were measured. Light-evoked IPSCs were recorded at a holding potential of -56 mV. All potentials indicated in the article were corrected for the 16 mV liquid junction potential. Series resistances never exceeded 20% of membrane resistance and were not compensated.

#### Optogenetic stimulation and drug application

Light was delivered through the ×40 objective by a 585 nm LED (CoolLED) driven by the acquisition software at 50 to 90% of its maximal power (0.6 to 1.0 mW at the sample). In the tVTA, *Sox14*+ neurons were identified by tdTomato fluorescence and by their action potential response to a 500 ms flash. When the stimulation was a train of 10 to 300 brief (1 ms) flashes delivered at 30 to 40 Hz (25 to 30 ms between flashes), fidelity was the proportion of flashes that evoked a spike, in %. In the VTA, Sox14+ tVTA fibers were stimulated with a single 1 ms flash, pairs of flashes 50 ms apart or trains of ten flashes at 20 Hz. Gabazine (5 µM, Abcam, #ab120042) and DAMGO (1 µM, Abcam, #ab120674) were bath applied for at least 5 min.

#### Morphological recovery and tyrosine hydroxylase (TH) immunodetection

The pipette was slowly withdrawn at the end of the recording and the slice was fixed in 4% PFA for at least 24 h at 4°C, then rinsed three times in PBS. Neurobiotin was revealed by an overnight incubation at 4°C, in the dark and with agitation, in PBS containing 0.3% Triton X-100, % goat serum and streptavidin conjugated to Alexa Fluor 488 (1:1,000; Supplementary Table 4). In slices processed for TH, the same blocking solution was used for an overnight incubation at 4°C with a chicken anti-TH antibody (1:1,000), followed after three rinses by 2 h at room temperature with an anti-chicken secondary antibody (STAR635P, 1:500) together with the streptavidin. After three rinses, slices were mounted in Fluoromount-G (Invitrogen) and imaged on a Zeiss Axio Imager M2 epifluorescence microscope equipped with an X-Cite 120LED light source and a Hamamatsu camera (6.5 µm pixels, 2 × 2 binning), driven by ZEN software (v3.5, Zeiss Microscopy). The recording site of each neuron was assigned post hoc by visual inspection of the fixed slice.

#### Analysis and statistics

Action potentials recorded in LCA were detected in AxographX with an amplitude threshold set for each recording. Their duration was measured on an average trace, from the onset of the negative peak, taken at 10% of that peak, to the maximum of the positive peak^36^. For spontaneous discharge, the interval between consecutive action potentials (interspike interval, ISI) was measured, and its coefficient of variation computed as CV = s.d./mean. In LCA recordings of VTA neurons, the ISI preceding the flash (ISI1) and the next ISI, encompassing the flash (ISI2), were each averaged over at least twenty consecutive episodes recorded every 10 s. VTA neurons recorded in LCA were considered as putative dopamine neurons when they fired regularly (CV < 0.4) with wide action potentials (duration > 1.2 ms)^36,37^. In whole-cell recording, the delay of the first action potential evoked by a depolarizing current step was measured from the onset of the current injection to the base of that action potential; a delay >100 ms is typical in dopamine neurons^36^. The latency of a light-evoked IPSC was measured from the onset of the flash to the point where the current reached 10% of its peak, and its jitter was the s.d. of that latency across sweeps; its amplitude is the peak of an average response computed from several sweeps, and its rise time the interval from 10 to 90% of that peak. The IPSC decay was fitted by a double exponential with *t* = 0 at the peak, and the values given in the text are weighted decay time constants, τw = (τ1A1 + τ2A2)/(A1 + A2). The paired pulse ratio is the amplitude of the second IPSC over that of the first. For the current-voltage relation, IPSCs were recorded at holding potentials between -96 and -36 mV, each amplitude was normalized to that of the same neuron at -56 mV, and the reversal potential was read from a linear fit. Six neurons were excluded from the analysis of the IPSC kinetics (**Supplementary Table 10**). Comparisons were paired within neurons, before against after DAMGO and ISI1 against ISI2. Exact *P* values are given test by test in **Supplementary Table 11a**.

### In vivo electrophysiology and fiber photometry

Recordings were performed 4 to 6 weeks after injection, during the active phase of the animals. Five mice (three males and two females) were used for *in vivo* extracellular single-unit recording, four of which (two males and two females) also underwent fiber photometry recording. Mice received two stereotaxic injections in the right hemisphere, 180 nL of AAV5-Syn-FLEX-rc[ChrimsonR-tdTomato] (Addgene, #62723-AAV5) in the tVTA (AP -4.0, ML +0.8, DV -4.2, 5° lateral approach), and 480 nL of AAV5-hSyn-dLight1.2 (Addgene, #111068-AAV5) in the NAc (AP +1.5, ML +1.2, DV -3.8, vertical).

#### Recording surgery

Anesthesia was induced with urethane (Sigma, 250 mg/mL in distilled water; 1 g/kg, i.p.), supplemented with isoflurane during induction (3%, 2 L/min air) and surgery (1-2%, 0.5 L/min air). The mouse was then placed in a stereotaxic frame and prepared as described above for the viral injection, with the craniotomies re-drilled over the recording sites. Two screws were placed in contact with the dura for the electroencephalographic (EEG) recording, one over the prefrontal cortex (AP +2.0, ML -2.0) and one over the cerebellum (AP -7.0, ML -2.0), with a silver electrode under the skin of the neck as ground. Isoflurane was switched off at least 1 h before any recording. Depth of anesthesia, characterized by slow-wave activity on the EEG and by the absence of a hindpaw withdrawal reflex to pinch, was monitored throughout the session and maintained with additional urethane as needed (20-50 µL per injection, i.p., at least 30 min apart).

#### Optogenetic stimulation

A 200 µm optical fiber (FP200URT, 0.50 NA, Thorlabs) was positioned just above the tVTA (AP -3.9, ML +0.9, DV -3.9, 5° lateral approach), 100 µm lateral to the injection site to leave room for the recording pipette. Light came from a red LED (625 nm; LEDFRJ_625, Doric Lenses), and output at the fiber tip was 4.0 to 4.6 mW, measured before insertion. Pulse trains were generated by the TTL sequencer of Spike2 v8.00a (2013) through a CED Power1401 interface (Cambridge Electronic Design). All stimulation pulses were 5 ms long. In each animal the fiber photometry recording was performed first, and the single-unit recordings followed within the same session, up to four hours later.

#### Fiber photometry

A 400 µm fiber (FP400URT, 0.50 NA, Thorlabs) was lowered just above the NAc (AP +1.5, ML +1.2, DV -3.6) to collect dLight1.2 fluorescence. Each stimulation frequency was tested with ten trains of ten pulses at 10, 20, 30, 40 or 50 Hz, with 30 s between trains, the trains lasting 900, 450, 297, 225 and 180 ms respectively. The 20 Hz series was not run in animal BM36. Fluorescence was recorded with a Doric Lenses system. Two excitation wavelengths were used, 465 nm for the dLight1.2 signal (LEDC1-B_FC) and 405 nm as an isosbestic reference (LEDC1-405_FC), both driven by a two-channel programmable LED console (LEDD_4) and combined into the collection fiber through a minicube (FMC2). The fiber was photobleached before use and light intensity at its tip was held between 10 and 30 µW throughout every session. Emitted fluorescence was collected through the same fiber onto a photoreceiver (DFD_FOA_FC), demodulated online by the acquisition software (Doric Neuroscience Studio v6.1.1.1) and sampled at 12048 Hz.

The demodulated 465 nm signal was smoothed with a Savitzky-Golay filter, window 5 ms and order 3, and corrected for photobleaching by fitting a monoexponential F₀(t) = a·e^-bt^ + c to that same channel over the whole recording. The normalized signal is ΔF/F(t) = [F(t) - F₀(t)] / F₀(t). The 405 nm channel was acquired as a control and does not enter this calculation. A trial was the window from -2 to +2 s around the first pulse of a train, resampled to 1 ms. Within a recording, every trial was z-scored on its own baseline (2 s preceding train onset), and the ten z-scored trials were averaged, giving one trace *per* animal and frequency. Across animals, these traces were averaged and the band is the s.e.m. across animals wherever a group trace is drawn. The animal response was tested against its own baseline by comparing the mean z-score over 0 to +1 s with the matched window -1 to 0 s, paired trial by trial.

All metrics were measured on the mean z-scored trace of one animal at one frequency, and are listed *per* animal and frequency in the Source Data. Onset is the first time at which the trace falls below 1.2 s.d. of its baseline, end is the first time at which it returns above that threshold, and duration is their difference. Excess over the light is duration minus the measured train duration. Trough is the lowest value of the trace over 0 to +2 s. Two further measures were read after a centered 21 ms moving mean.

Three controls were run on the 30 Hz recording of each of the four animals and are reported in **Supplementary Table 11b**. For the isosbestic control, the 405 nm channel was processed exactly as the 465 nm channel and the two were compared over 0 to +1 s. For the return to baseline, the same recordings were re-cut from -2 to +10 s and each successive 2 s window was compared with the 2 s preceding the train. For the independence of successive trains, the mean ΔF/F of the 2 s preceding each train was regressed on the rank of the train. Levels are expressed in s.d. of the baseline recorded before the first train of the session.

#### Single-unit recording

At the end of the photometry acquisition the NAc fiber was withdrawn, the stimulation fiber being left in place, and a glass micropipette (1-3 µm tip diameter, 10-20 MΩ; Harvard Apparatus, #30-0044 GC120F-10) filled with 0.9% saline was lowered at 5 µm/s to just above the target with a motorized micromanipulator (LinLab, Scientifica), then advanced at 1 µm/s while light pulses were delivered regularly, a neuron being detected either by its spontaneous discharge or by its response to the light. Three to seven tracks were made *per* animal through the tVTA (DV -3.5 to -4.5) and, in one animal, also through the VTA (DV -3.5 to -5.5). Three tVTA stimulation protocols were analyzed: trains of 10 pulses at 30 Hz, of 10 pulses at 50 Hz, and of 30 pulses at 30 Hz lasting 957 ms, with 30 s between trains in each protocol. The three dopamine neurons were all stimulated at 30 Hz but not with the same protocol, 10 trains of 10 pulses in one, 20 and 50 trains of 30 pulses in the other two. Signals were acquired, amplified and filtered (0.3-5 kHz bandpass) with conventional electronics (Neurodata IR183A preamplifier; Humbug 50 Hz noise eliminator, World Precision Instruments; Brownlee 440 differential amplifier), and collected online with a laboratory interface and software (CED Power1401, Spike2 v8.00a; Cambridge Electronic Design) at 6666.67 Hz. The EEG was filtered from 0.30 to 1.5 kHz by the same amplifier. Spikes were isolated from the noise in Spike2, a detection threshold being placed visually just above the noise level of each recording and every threshold crossing stored as a waveform window. These events were then separated from artefacts, and occasionally from a second unit, by multidimensional principal component analysis. Sorted events of a single stored sample, lacking the positive rebound of a true action potential, were removed on the correlation of their waveform to the light-evoked mean waveform of the same recording. Where the signal-to-noise ratio was inconsistent and visual thresholding failed, detection was performed on the raw unit channel at three times the robust noise, and events were kept only when their waveform matched the light-evoked template of the same recording. This approach was applied to all files of the 957 ms trains and of the 30 against 50 Hz comparison.

#### Light response and photoidentification

The latency is the smallest interval between a spike and a pulse within 0 to 25 ms. A spike belongs to the response peak of a neuron when its latency is within 1.3 ms of the median latency of that neuron. Fidelity is the proportion of pulses followed by such a spike, and jitter is the standard deviation of the peak latencies. Spikes *per* pulse is the number of spikes in the light window divided by its ten pulses (**Supplementary Table 13**). Chance level was computed for each neuron from its own spontaneous rate as 1 - e^(-rate^ ^×^ ^window)^, the spontaneous rate being measured between blocks, with 0.5 s excluded before and 2 s after each block. For neurons with no spontaneous spike the rate is an upper 95% Poisson limit (**Supplementary Table 12**). In each recording, only the first uninterrupted series of 30 Hz trains was analyzed. The same three windows were used for the tVTA and the dopamine neurons: the 500 ms preceding train onset, the first 333 ms of the train, and 500 ms starting 25 ms after the last pulse, the gap leaving room for the response to that pulse. The light window is anchored on train onset and the after window on the last pulse, because the trains differ in length. *Sox14*+ neurons in the tVTA were photoidentified when they responded to the light pulses above the chance level, no waveform criterion being used. For VTA recordings, the criteria used to identify putative dopamine neurons were those from previous studies^38–40^, known to be indicative^70^, namely: *(1)* an action potential with a biphasic or triphasic waveform of at least 2.5 ms in duration; *(2)* at least 1.1 ms from spike onset to the negative trough; and *(3)* a slow spontaneous firing rate <10 spikes/s. Measurements were made in Spike2 on the mean waveform of each unit (**Supplementary Table 12**).

#### Histology

At the end of the recording session the animal was decapitated, the brain collected and fixed for 48 h at 4°C in 0.1 M PB with 4% PFA, then stored at 4°C in PBS with thimerosal until sectioning. Serial coronal sections were cut at 40 µm on a vibratome (VT1000S, Leica) and collected in PBS. One section every 120 µm was processed, expression of dLight1.2 in the NAc being revealed by IHC against GFP on free-floating sections (rat anti-GFP, 1/1000; anti-rat Alexa Fluor 488, 1/400; **Supplementary Table 4**), while ChrimsonR-tdTomato was imaged from its native fluorescence. Sections were mounted on SuperFrost slides, air-dried, counterstained with DAPI (0.5 µg/mL, 30 s) and coverslipped in Fluoromount-G. Sections were imaged on a NanoZoomer® S60 scanner (×40 objective, 0.23 µm/pixel). Every animal was checked for opsin expression in the tVTA, sensor expression in the NAc and the position of the two optical fibers. Eight mice were injected and three were excluded, for absent or misplaced viral expression, for a misplaced optical fiber, or for both.

#### Analysis

Tests were run within a neuron or within an animal, without correction for multiple comparisons, every member of a reported family being given, significant or not. The trend of fidelity along the 957 ms trains was tested on a planned linear contrast of the six blocks of five pulses, weights -5 -3 -1 +1 +3 +5, one contrast *per* train. Photoidentification was tested on the responses falling within the peak window of each neuron. Every test, with its unit, its exact *P* and its effect size, is listed in **Supplementary Table 11b**. On graphs, \**P* < 0.05, \*\**P* < 0.01, \*\*\**P* < 0.001, n.s. not significant, and n.t. not testable, meaning that the smallest *P* the number of repetitions could have returned is itself above 0.05.

### Whole-brain connectome of *Sox14*+ tVTA neurons

#### Viral vectors and tracing strategies

For anterograde tracing, mice received an injection of AAV5-hSyn-DIO-EGFP (Addgene, #50457; 10¹³ vg/mL; 120 nL) in the tVTA (AP -4.0, ML +0.8, DV -4.2, 5° lateral approach). The pipette was withdrawn step by step to limit transduction of *Sox14*+ neurons along the track, in particular in the superior colliculus. Mice were perfused three weeks later, and only mice with transduction restricted to the tVTA were retained for analysis (*n* = 4 of 5 injected). For monosynaptic retrograde rabies tracing, five mice (three males and two females) received a first injection of a Cre-dependent helper AAV5-EF1a-DIO-TVA-V5-t2A-RabiesG expressing, from a single cassette, the avian receptor TVA fused to a V5 epitope tag together with the rabies glycoprotein (TVA-V5-RG; 120 nL), in the tVTA through the same angled approach. Three weeks later the same animals received an injection of an EnvA-coated, glycoprotein-deleted rabies virus expressing nuclear GFP (RV-H2B-GFP; 250 nL) at the same antero-posterior coordinate but vertically, without angle (AP -4.0, ML +0.4, DV -4.0), so that the first pipette track was not reused. Both vectors were produced by the laboratory of K. Meletis (Karolinska Institutet, Stockholm, Sweden), and the constructs and tracing procedure were as described previously^42–44^. The rabies surgery was performed 15 days before perfusion. Two of the five injected mice (one male and one female) showed GFP labeling at the injection site and were retained for analysis; the three others (two males and one female) showed no or only sparse labeling and were excluded.

#### Perfusion and histology

Mice were perfused and the brains post-fixed as described above, then stored at 4°C in PBS with 0.02% NaN₃. Serial coronal sections were cut at 40 µm on a vibratome (VT1000S, Leica) and collected in PBS. Each brain was sectioned in full and one section in three was mounted, giving a sampling interval of 120 µm and 63 to 74 mounted sections per brain. Sections from rabies-traced brains were first processed as free-floating sections for IHC (see below). Sections were mounted on SuperFrost Plus slides, air-dried, counterstained with DAPI (0.5 µg/mL, 30 s) and coverslipped in Fluoromount-G (Electron Microscopy Sciences, #EM-17984-25). Every mounted section was imaged, registered and quantified. The series covered the brain from +2.9 to -6.6 mm relative to bregma, corresponding to 2.4 to 11.9 mm along the antero-posterior axis of the Allen Mouse Brain Common Coordinate Framework (CCFv3), the reference space in which all sections were subsequently registered; the rostral and caudal limits varied between animals (rostral, +2.0 to +2.9 mm; caudal, -5.7 to -6.6 mm), so that the most caudal medullary levels were not sampled in every brain.

#### Immunohistochemistry

IHC performed on free-floating sections for rabies-traced brains; sections from anterograde tracing were mounted directly, the EGFP signal being imaged from its native fluorescence. Incubations were carried out at RT, under gentle agitation and protected from light. Sections were rinsed three times for 5 min in PBS, blocked for 1 h in PBS containing 0.2% Triton X-100 (PBS-T) and 2% normal serum from the species of the secondary antibody, and incubated overnight with the primary antibody in PBS-T containing 1% of the same serum. After three 5-min rinses in PBS, sections were incubated for 2 h with the corresponding fluorophore-conjugated secondary antibody in PBS-T, rinsed three times for 5 min in PBS and stored in PBS with 0.02% NaN₃ until mounting. Every section was immunostained for the V5 tag, which identifies helper-transduced cells, using a chicken anti-V5 antibody (1:1,000) revealed with a goat anti-chicken IgY Alexa Fluor Plus 647 antibody (1:400). TH was additionally immunolabeled on a subset of sections to delineate the dopaminergic midbrain (eight sections, -2.91 to -3.60 mm from bregma) and the locus coeruleus (four sections, -5.51 to -5.63 mm from bregma), using a mouse anti-TH antibody (1:1,000) revealed with a goat anti-mouse IgG Alexa Fluor 594 antibody (1:400). Antibodies information in **Supplementary Table 4.** Rabies-driven nuclear GFP was imaged from its native fluorescence.

#### Image acquisition

Whole-section images used for brain-wide quantification were acquired on a Zeiss Axio Imager M2 epifluorescence microscope equipped with an EC Plan-Neofluar 20×/0.50 objective, driven by ZEN software (v3.5, Zeiss Microscopy). Each coronal section was imaged as a tile scan covering the whole section, acquired in 16-bit and exported as 8-bit composite images with a final pixel size of 0.65 µm. Rabies-traced sections were acquired in three channels (DAPI, EGFP and V5), with a fourth channel on the subset of sections immunolabeled for TH; anterograde sections were acquired in two channels (DAPI and EGFP). Injection sites were imaged separately on a Leica Stellaris 8 confocal microscope with a 40× objective, as 16-bit tile scans covering the whole injection territory, acquired as a single optical plane of 5 µm with a pixel size of 0.284 µm.

#### Atlas registration, region annotation and signal detection

Each coronal section was registered to the Allen Mouse Brain CCFv3 (10 µm) using ABBA (Aligning Big Brains and Atlases; QuPath extension v0.3.3)^71^. Sections were first assigned to the corresponding atlas plane, and the alignment was then refined manually on anatomical landmarks by an affine transformation followed by non-linear spline warping. The warped atlas annotations were imported into QuPath (v0.6.0), with the left and right hemispheres kept as separate annotations, and the annotated area of every region on every section was returned. Region annotations covering damaged, folded or missing tissue, or overlapping the tVTA itself, were marked for exclusion on the sections concerned. Labeled nuclei in rabies-traced sections were detected on the EGFP channel with the BraiAn QuPath extension (v1.1.0)^71^, configured through a project-level YAML file (watershed cell detection; 0.65 µm pixel size, detection threshold 5, cell area 20 to 150 µm², cell expansion 3 µm). Fibers in anterograde sections were segmented on the GFP channel with pixel classifiers trained in QuPath (OpenCV random trees, 50 trees, depth 25, minimum 10 samples *per* node; 0.65 µm *per* pixel, 512 by 512 pixel tiles; six families of texture features at four spatial scales), each pixel being assigned to fiber or to background. For each animal, a classifier was trained on one section chosen as representative of the labeling density and applied to the whole series; sections on which the segmentation was visually unsatisfactory were used to train further classifiers, applied in turn, until the segmentation was satisfactory throughout. Two mice required a single classifier and the two others five and six. For both datasets, region-level measurements, that is labeled-cell counts or classified fiber area together with the annotated area of each region, were exported from QuPath as a single table *per* animal.

#### Pipeline

In both datasets, whole ontological branches were removed before aggregation (fiber tracts, ventricular systems, retina, cerebellum, cerebellar nuclei, grooves), and the regions marked for exclusion in QuPath were discarded. Sections mounted in reversed orientation were corrected by swapping the left and right hemisphere labels, the correction being verified by the *per*-section left to right balance of labeling, and the ipsilateral side was then assigned *per* animal.

Signal was measured on both hemispheres and analyzed separately: the maps presented are based on ipsilateral signal. Analysis was restricted to structure level 8 of the Allen ontology, a single level, so that no region was counted twice, merging four groups of structures based on neurochemical or functional grounds: the VTA complex (VTA-cx: VTA, interfascicular nucleus raphe, paranigral nucleus, rostral linear nucleus raphe and medial terminal nucleus of the accessory optic tract), which corresponds to the A10 dopaminergic group, the medial terminal nucleus being included because of atlas imprecision at this level; the locus coeruleus (LC: locus coeruleus and subcoeruleus nucleus), which corresponds to the A6 noradrenergic group; the parasubthalamic nucleus (PSTN: parasubthalamic and preparasubthalamic nuclei); and the medullary raphe (RaM: nucleus raphe magnus, nucleus raphe obscurus and nucleus raphe pallidus), which correspond to the B1 to B3 serotonergic groups. Region names and acronyms follow the Allen CCFv3 nomenclature throughout, so that LH denotes the lateral habenula and LHA the lateral hypothalamic area. Thirty structures were then excluded from both datasets. Eleven cranial nerve nuclei, motor or cholinergic structures with no documented monosynaptic connection with the tVTA, and ten cranial nerve fiber bundles, which are not neuronal populations, were excluded, together with four generic aggregating categories of the Allen ontology serving as registration landmarks rather than as structures. As the tVTA is not delineated in the Allen atlas, three regions that may partially contain it were also excluded: the retrorubral area, the red nucleus and the central linear nucleus raphe. The inferior colliculus was excluded, no monosynaptic connection with the tVTA being documented, and the anterior tegmental nucleus as it was detected in a single animal.

For the tVTA inputs, EGFP counts and annotated areas were summed across all sections for each region or group of merged regions, and density was computed as the ratio of these two sums, expressed in cells/mm². A region was retained if it was detected in both mice with at least 3 EGFP cells, and if the density averaged over the two mice reached at least 5 cells/mm², a minimum set a priori and more than twice the 95^th^ percentile of the background density measured in 34 reference regions that receive no tVTA input. This minimum was lowered to 3 cells/mm² for the cortical subplate, the ventral striatum, the medial amygdalar nucleus and the parvicellular reticular nucleus, and waived for the isocortex, the olfactory areas and the lateral septal nucleus, where labeling is genuinely sparse and large reference volumes dilute density. This procedure yielded 84 afferent regions.

For the tVTA outputs, annotations smaller than 2,000 µm² were removed before aggregation. Fiber areas and annotated areas were then summed across all sections for each region, and fiber density was expressed as the percentage of the region area occupied by fibers. A detection threshold was derived from the data for each animal: background was estimated as the median fiber density across the regions of the isocortex, cortical plate and hippocampal region, pooled over both hemispheres, these being areas where tVTA projections are negligible, and the threshold was set at three times this value. For the two mice processed with several classifiers, background was estimated separately for each classifier and the highest of these medians was retained. Thresholds ranged from 0.024 to 0.033% of fiber area across the four mice. Densities below the animal threshold were set to zero and excluded from the across-animal statistics, the mean, s.e.m. and coefficient of variation of a region being computed only over the mice in which it exceeded the threshold, and a region was retained if it exceeded the threshold in at least three of the four mice, on either side. Within the isocortex, cortical plate, cortical subplate, hippocampal region, ventral striatum and periventricular region, where tVTA projections are sparse, a region was also retained if it exceeded the threshold in two mice with an inter-animal coefficient of variation below 80%, so that sparse but reproducible signal was not discarded while isolated signal was. This procedure yielded 110 efferent regions.

#### Connectome architecture

The afferent and efferent region lists were merged in a table of 124 regions. A region present in both lists was called reciprocal, the others afferent- or efferent-only. Within each list, regions were ranked by ipsilateral signal, rank 1 being the strongest, and each region assigned to one of nine anatomical super-groups (cortex, striatum, pallidum, amygdala, thalamus, hypothalamus, midbrain, pons and medulla) based on the Allen ontology. Reciprocal regions ranking in the top 30% of both lists were called hubs, that is, with an afferent rank of 26 or better and an efferent rank of 33 or better. The concentration of each distribution was summarized by the proportion of the summed ipsilateral signal carried by its five densest regions, and by the Gini coefficient of the regional values. The preference of the unidirectional regions for one direction was tested on the 54 afferent-only and efferent-only regions with an exact two-tailed binomial test against 0.5.

For each animal, the ipsilateral signal was also overall summed *per* section and plotted against the antero-posterior coordinate of that section, taken from the atlas registration. Each profile was scaled to unit area by trapezoidal integration, then linearly interpolated onto a common axis running from 2 to 13 mm (CCFv3 coordinate) in steps of 0.05 mm, smoothed with a Gaussian kernel of 0.15 mm and averaged across animals. The position of each peak was read as the maximum of the averaged profile, excluding the tVTA itself, and approximate bregma coordinates were estimated as 5.30 mm minus the CCFv3 coordinate.

For the schematic map, a region was drawn when it carried at least 1% of the total ipsilateral signal (labeled cells for inputs and labeled fiber area for outputs), computed *per* mouse and averaged across mice; 24 input and 14 output regions met this criterion and were displayed in three classes of proportion (at least 4%, 2 to 4%, 1 to 2%). Each region is an ellipse whose center and second moments were computed from all voxels of the region in one hemisphere, both hemispheres for unpaired midline structures, in the Allen CCF 2020 annotation volume (25 µm), projected on the horizontal plane and drawn at half the atlas extent. The background is a horizontal section of the atlas 4.5 mm below its dorsal edge, inputs in one hemisphere and outputs mirrored in the other, and the cross is marking the center of the tVTA territory as mapped in the Allen atlas (**Supplementary Fig. 9**).

#### Injection site

The tVTA is not delineated in the Allen atlas. In both tracing experiments, its boundaries were therefore drawn manually on each injection-site image, from anatomical landmarks and molecular delineation previously established in this study. Counts were obtained on the confocal sections of a single 5 µm optical plane, whereas areas were measured on the widefield images of the whole series. In the rabies experiment, labeled cells were counted manually within these outlines and classified as GFP-only, V5-only, or double-labeled (*i.e.* starter cells), the somatic V5 signal being too diffuse for reliable automated segmentation. The tVTA, V5-positive and starter areas were measured on the same sections. Each density was the summed cell count divided by the summed tVTA area, starter density from double-labeled cells for each side, and helper-negative GFP+ density from GFP-only cells ipsilaterally. In the anterograde experiment, transduced somata were counted within and outside the tVTA outline, the injection area was measured, and somatic density computed as above. For both series, the injection index was computed on these densities as LI = (ipsilateral - contralateral) / (ipsilateral + contralateral), and the antero-posterior center of mass (CoM) was obtained by weighting each section by its starter (retrograde) or injection (anterograde) labeled area. Specificity was the proportion of EGFP-labeled somata within the tVTA, computed for each side on confocal sections.

#### Lateralization

The lateralization index (LI) was computed on ipsilateral and contralateral signal obtained for each animal separately, and then averaged. Regions were classified as strongly ipsilateral (LI > 0.5), ipsi-dominant (0.2 to 0.5), bilateral (absolute LI at most 0.2), contra-dominant (-0.2 to -0.5) or strongly contralateral (< -0.5). The index was computed on the measured values of both sides, for every animal with signal on at least one side, a side below threshold entering with its measured value so that no index was forced to +1 or -1; a region carried an index when at least two animals contributed, and all 84 afferent and all 110 efferent regions carried one. No index was computed for the tVTA itself. Each distribution of regional LI was tested against zero with a two-tailed one-sample Wilcoxon signed-rank test, its median is given with a 95% bootstrap confidence interval from 20,000 resamples. Inputs and outputs were never compared to each other.

#### Analysis

Group values are given as mean ± s.e.m. across animals and alpha was 0.05 throughout. Inter-animal agreement was assessed on log-transformed ipsilateral densities, with a Pearson correlation between the two rabies mice over the 74 afferent regions to which a non-zero density minimum applied, and between each pair of anterograde mice over the regions above the detection threshold of both. Variability between animals was summarized by the coefficient of variation, *i.e.* s.d. divided by the mean, on the ipsilateral signal of the mice above threshold. Effect sizes are the Clopper-Pearson interval for a proportion, the Pearson *r* with a Fisher-transform interval for a correlation, and the matched-pairs rank-biserial correlation for the Wilcoxon test. Every test, its exact *P* and its effect size are listed in **Supplementary Table 11c**.

#### Behavioral experiments

Behavioral procedures were performed during the dark phase, under red light. During tests, video tracking (Computar, MEGAPIXEL, #28895) and red LED activation (635 nm; LEDFRJ-R_FC, Doric Lenses; maximum output power: 5.5 mW at fiber tip) were controlled using ANY-maze software (Aniphy®). Animal locomotion was recorded and analyzed using the ANY-maze analysis module (v7.2, Stoelting Co.) Locomotor activity was defined as the time during which the animal was detected as mobile, and freezing as the time during which it was detected as freezing, over minimum episode duration of 2 s. Some detection artifacts occurring during place preference recordings (*e.g.* erroneous detection of the animal due for example to the stimulation cable) were identified and manually removed. The edited video was then reanalyzed using AnyMaze with the same tracking parameters, and the missing data replaced by the animal’s mean activity immediately before and after the edited segment.

#### Experiment 1: tVTA optogenetic stimulation

In C57BL/6J and Sox14-Cre mice, 180 nL of AAV5-Syn-ChrimsonR-tdT (Addgene, #59171) or AAV5-Syn-1(S) tdTomato-WPRE (Addgene, #51506) were delivered into the right LH (AP -1.7, ML 0.4, DV -2.8, vertical). In Sox14-Cre mice, the same method was used to deliver 180 nL of AAV5-Syn-FLEX-rc[ChrimsonR-tdTomato] (Addgene, #62723) or AAV5-hSyn-DIO-mCherry (Addgene, #50459) into the right tVTA (AP -4.0, ML 0.8, DV -4.2, 5° lateral approach), leading to bilateral transfection. In all groups, the optic fiber (MFC_200/250-0.66_4.5mm_ZF1.25_FLT, Doric) was implanted 200 µm above the tVTA (AP -4.0, ML 0.8, DV -4.0, 5° lateral approach).

After 4 weeks of recovery, a real-time place aversion (RTPA) was conducted. The arena consisted of two identical Plexiglas chambers (23 x 23 x 17 cm) separated by an opening (8.5 cm) and covered with clean wood shavings (∼0.5 cm). On the first day (pretest), mice were plugged and free to explore the apparatus for 20 min. Animals spending over 70% of the total time in one chamber were excluded from the study. The second day, optogenetic stimulation was turned on when the mouse entered its head and forepaws in the stimulation-paired chamber and turned off when it quitted the compartment. The C57BL/6J mice underwent stimulation of LH terminals in the tVTA under a protocol similar to Stamatakis and Stuber^9^ (60 Hz continuous, 5 ms light pulse duration, 635 nm illumination). The Sox14-Cre mice underwent optogenetic stimulation based on our ex vivo electrophysiological tests (30 Hz, 1 sec on / 1 sec off, 5 ms light pulse duration).

One week later, a first open-field procedure was conducted in a square open-field arena (40 × 40 × 30 cm). It consisted in a single 6-min test of free exploration. Following connection of the patch cord, mice were placed in the center of the arena. The stimulation protocol was identical across all groups (30 Hz, 5 ms pulse width, continuous). After an initial 2-min baseline period without stimulation, optogenetic stimulation was delivered for 2 min, followed by a 2-min post-stimulation period without stimulation. This design allowed assessment of locomotor activity before, during, and after stimulation (0-2 min off, 2-4 min on, 4-6 min off). Five weeks later, a second procedure was conducted consisting in four 10 min-sessions performed in the open field, with different stimulation protocols during the last 5 min. During the first day no stimulation was applied, the 2^nd^ day consisted in stimulation at 30 Hz, 5 ms, 5 s on / 15 s off, the 3rd day in continuous 30 Hz stimulation and the last day in continuous 60 Hz stimulation

One week later, a repeated RTPA was conducted. The apparatus was of the same size as described above, but included distinct visual cues differentiating the two compartments (solid black walls vs. black-and-white striped walls). Experiments were conducted over 4 consecutive days (10 min *per* day) under ∼45 lux illumination to facilitate visual discrimination. Day 1 served as a pretest following the same procedure as the above RTPA baseline session, and days 2 to 4 consisted in sessions during which optogenetic stimulation (30 Hz, 5 ms pulse width, continuous) was delivered upon each entry into the stimulation-paired compartment. One mouse from the tVTA group lost the optic fiber during the pretest and was thus excluded from this test.

#### Experiment 2: optogenetic stimulation of *Sox14*+ tVTA neurons or of their VTA/SNc terminal

Surgery was performed as described above, with tVTA transfection and optic fiber placed either above tVTA or above the VTA/SNc region (AP -3.2, ML 1.0, DV -4.0, 5° lateral approach; or AP -3.1, ML 1.0, DV -3.7, vertical). Four weeks later, mice underwent the RTPA task using differentiated compartments. The protocol was conducted over 6 days, including pretest, 4 stimulation sessions and a conditioned test in absence of stimulation. One week later, mice underwent an open-field procedure (0-2 min off, 2-4 min on, 4-6 min off).

#### Experiment 3: optogenetic stimulation of VTA or SNc terminals

Surgery was performed as described above, using a smaller volume (66 nL) of viral vectors and thinner optic fibers (100 μm core, 0.22 NA; RWD Life Science, #R-FOC-BL100C-22NA) to discriminate the VTA (AP -3.2, ML 1.0, DV -4.0, 5° lateral approach) from the SNc (AP -3.3, ML 1.25, DV -3.9, vertical). Stimulation and behavioral testing were performed as described for Experiment 2, with a one-week delay between RTPA and open-field tests. **Control of injection sites.** Animals were perfused as described above and the brains stored at 4°C in 0.1M PBS (pH 7.4) until cutting. Coronal sections (40 µm) were obtained using a vibratome (VT 1000S, Leica) and serially collected in PBS. Brain slices were mounted on slides (1 section every 120 µm) (Epredia™, SuperFrost Plus™). After drying, DAPI (0.5 µg/mL) was added for 30 seconds, removed, and coverslip set on the slide with Fluoromount-G mounting medium. Images were captured with a NanoZoomer® S60 Digital slide scanner (x40 objective, 0.23 μm/pixel, NDP Scan v3.4.4) in the mCherry/tdTomato and DAPI channels. Mice in which the injection site or the optical fiber was misplaced were excluded from the analyses.

#### Analysis

Statistical analyses were performed in GraphPad Prism 8.4.2 (GraphPad Software, USA), using one- or two-way repeated-measures ANOVA followed by Sidak’s or Dunnett’s post-hoc tests. Every test, with its groups and *n*, its exact *P*, its effect size and its post-hoc comparisons, is listed in **Supplementary Table 11d**.

### Python analysis

Two conda environments were used. The bulk RNA-seq analyses were run in Python 3.12.14 with NumPy 2.5.2, pandas 3.0.5, Matplotlib 3.11.1, openpyxl 3.1.5, pydeseq2 0.5.4^72^ and adjustText 1.4.0, pandas 3 being required by pydeseq2. All other analyses were run in Python 3.11.14 with NumPy 2.4.2, pandas 2.3.3, SciPy 1.17.1, Matplotlib 3.10.8 and openpyxl 3.1.5, together with scanpy 1.11.5, anndata 0.12.18, igraph 1.0.0, leidenalg 0.12.0, cell_type_mapper 1.7.4, h5py 3.16.0 and pyarrow 24.0.0 for the Xenium analyses, and neo 0.14.5 and quantities 0.16.4 for the *in vivo* recordings. Versions are given in the requirements file of each repository.

### Artificial intelligence (AI)

The large language model Claude (Anthropic) assisted J.K. in writing the Python code for the analyses and graphs of the bulk RNA-seq, Xenium, *in vivo* recording (fiber photometry and electrophysiology) and connectome data. All codes were run and checked by J.K., who takes full responsibility for the analyses and their results.

## Data availability

The bulk RNA-seq and Xenium spatial transcriptomics datasets generated in this study will be deposited in the Gene Expression Omnibus upon publication. The raw data underlying the other figures will be deposited upon publication.

## Code availability

The analysis code will be made available on GitHub and archived on Zenodo with a DOI upon publication.

## ACKNOWLEDGEMENTS

This work was supported by the Centre National de la Recherche Scientifique (contract UPR3212), the University of Strasbourg, the French National Research Agency (ANR) through the ANR-24-CE37-3736 (E.D, D.D.S and J.K), ANR-22-CE37-0009 (M.B.), ANR-23-CE16-0016 (E.D.), and the Programme d’Investissement d’Avenir EURIDOL graduate school of pain ANR-17-EURE-0022 (B.M.M., P.A.D., L.M., and M.V.), an IdEx Young Investigator award of University of Strasbourg (J.K.), a NARSAD Distinguished Investigator Grant from the Brain & Behavior Research Foundation (ID24220; M.B.), the Association France Parkinson 2024-CS-0019 (B.M.M.), the Fondation pour la Recherche Médicale (FDT20170437322; F.F.), and EU ErasmusMundus Neurotime program (A.J., M.B). The authors would also like to acknowledge the Chronobiotron UAR3415 (CNRS UAR3415, University of Strasbourg, CELPHEDIA) for animal care, the In Vitro UAR 3156 imaging platform for microscopy, GenomEast platform, a member of the France Génomique consortium (ANR-10-INBS-0009) supported by the Grand Est Region (Spatial-seq project) and the Ligue Nationale contre le Cancer for Xenium and sequencing experiments, the PHEN-ICS platform (Université de Strasbourg, CNRS UAR2062 PHEN-ICS, INSERM US66, CELPHEDIA) and more specifically Patrice Goetz-Reiner for Xenium slide preparation, Hugues Jacobs and Olivia Wendling for Xenium experiment and Tania Sorg as a project manager, the CompOpt platform (CNRS UPR3212) for behavioral experiments, Alessio Delogu (King’s College London, London, UK) for the Sox14-Cre transgenic line, Konstantinos Meletis and Thomas Contesse (Karolinska Institutet, Stockholm, Sweden) for the rabies viruses, Dorothée Daniel for technical support. Behavioral and microscopy platforms were supported by the Région Grand-Est (Fonds Régional de la Coopération pour la Recherche, CLueDol project).

## AUTHOR CONTRIBUTIONS

Bulk genomic experiments: F.F., A.J., S.L.G., P.E.L., C.S., M.B., and J.K.; RNAscope experiments: B.M.M., E.C., G.F., P.H., and J.K.; spatial transcriptomics: E.C., E.D., C.E., and J.K.; mouse colony: S.D.; ex vivo electrophysiology: D.D.S.J., B.M.M., T.P.; in vivo electrophysiology and fiber photometry: B.M.M., P.A.D., and J.K.; connectome: B.M.M., E.C., L.M., and J.K.; behavioral experiments: B.M.M., M.V., M.B., and J.K.; experimental design: M.B. and J.K.; data analyses: B.M.M., S.L.G., D.D.S.J., M.B., and J.K.; fundings: M.B., and J.K.; manuscript preparation: B.M.M., E.C., M.B., and J.K.

