## Supplementary Figures 1-18 and Supplementary Tables 1-17 for "Multimodal definition and characterization of the tail of the ventral tegmental area through *Sox14*"

### Supplementary material

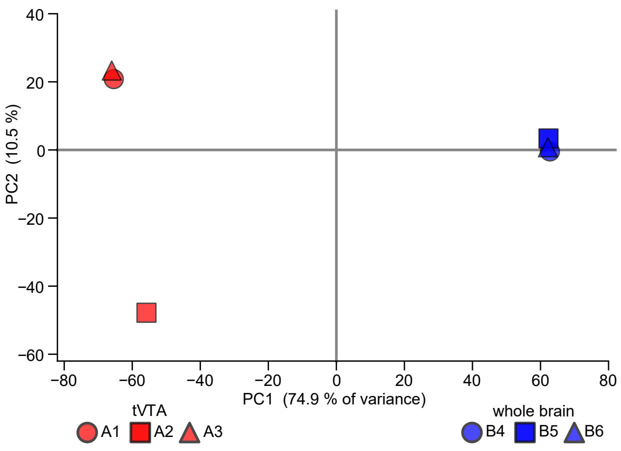


**Supplementary Figure 1 | The tVTA and whole-brain libraries separate on the first principal component.** A principal component analysis was computed on the variance-stabilized counts of the 32,643 genes with at least one read. Red: the three tVTA libraries (A1 to A3); blue: the three whole-brain libraries (B4 to B6); one symbol *per* library, as in **Fig. 1c**. The first component of the analysis separated the tVTA libraries from the whole-brain ones. Group sizes: 3 rats for the tVTA, 3 rats for the whole brain; one library *per* rat.

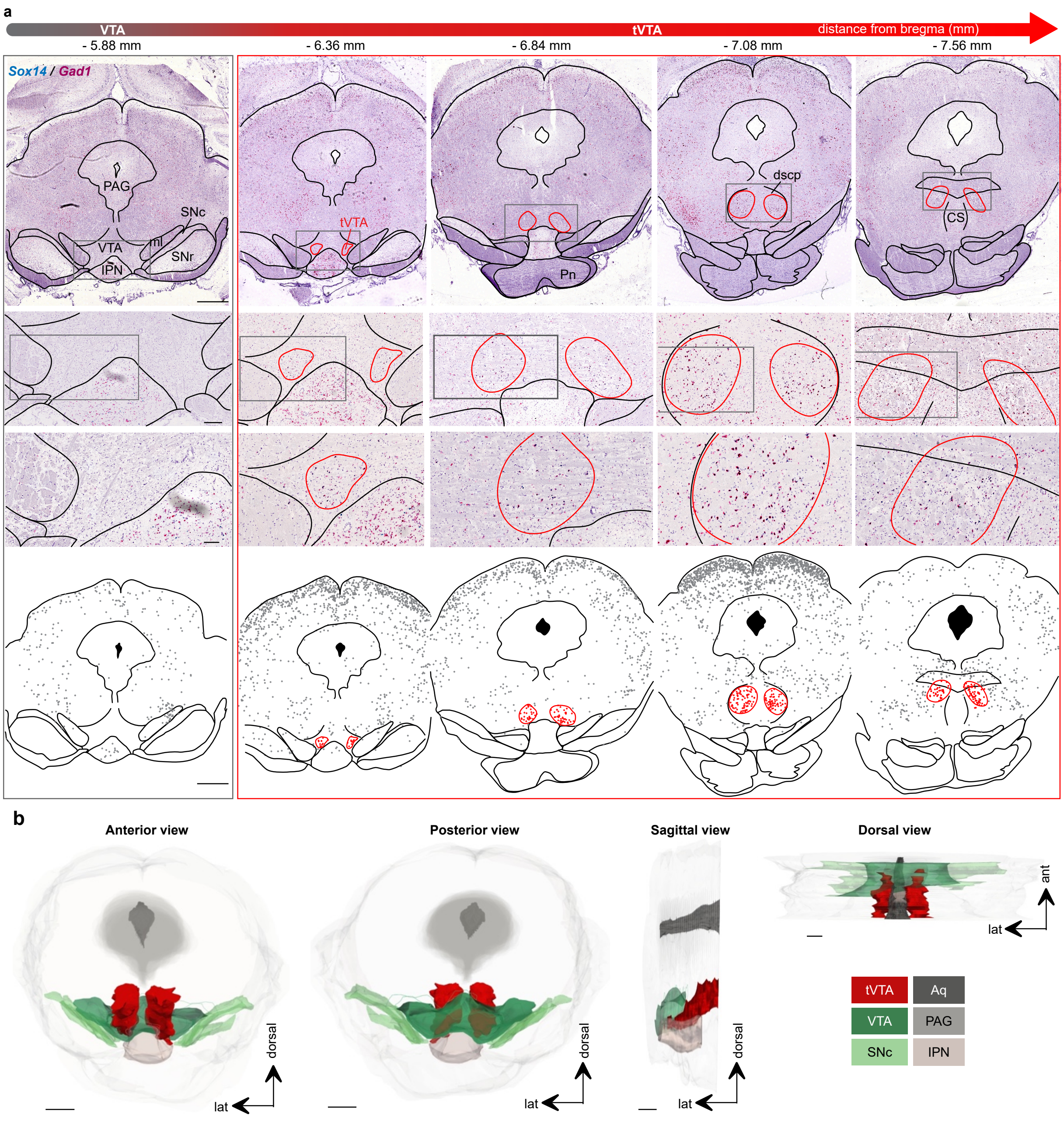


**Supplementary Figure 2 | *Sox14/Gad1 in situ* hybridization delineates the rat tVTA as in the mouse.** **a** Serial coronal sections of a rat brain along the mesopontine rostro-caudal axis, processed for duplex chromogenic RNAscope *in situ* hybridization against *Sox14* (blue) and *Gad1* (red) mRNA, with hematoxylin counterstain. The leftmost column is a reference level through the VTA, rostral to the tVTA (-5.88 mm from bregma); the four columns framed in red span the tVTA, from -6.36 to -7.56 mm from bregma. From top to bottom: whole coronal sections; the boxed VTA/tVTA region at two increasing magnifications; and the corresponding delineation of the tVTA (red outline) relative to the VTA and adjacent structures. The *Sox14/Gad1* double-labeled cell cluster defined the rat tVTA from its rostral pole, embedded within the caudal VTA, to its caudal end in the pons. As in the mouse, the structure shifted progressively dorsally along its caudal extension, following the decussation of the superior cerebellar peduncle (dscp), and its cross-sectional area increased caudally, the two horns lying farther apart than in the mouse. **b** Three-dimensional reconstruction of a rat tVTA (red) generated from serial *Sox14/Gad1* sections (one 14 µm section every eight; *n* = 1 rat), shown in anterior, posterior, sagittal and dorsal views and positioned relative to the VTA (green), substantia nigra pars compacta (SNc, light green), periaqueductal gray (PAG), cerebral aqueduct (Aq) and interpeduncular nucleus (IPN). Scale bars: 1 mm (**a**, top and bottom rows), 200 µm and 100 µm (**a**, upper and lower magnification rows), 1 mm (**b**, anterior and posterior views) and 250 µm (**b**, sagittal and dorsal views). CS, median raphe; ml, medial lemniscus; Pn, pontine nuclei; SNr, substantia nigra pars reticulata.

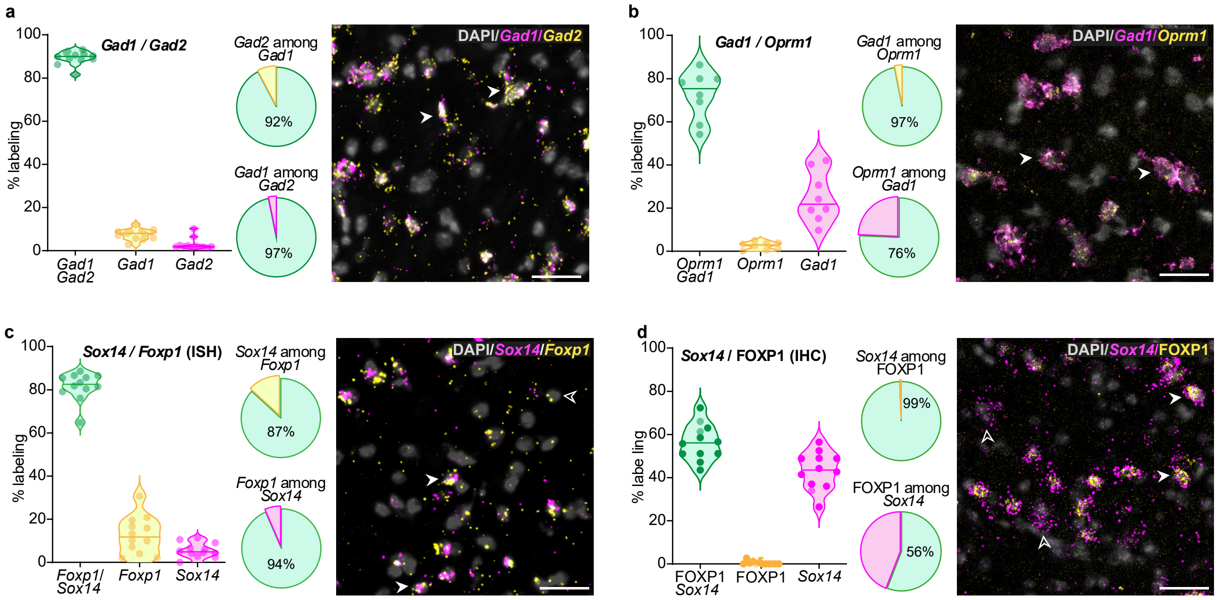


**Supplementary Figure 3 | *Oprm1* is almost confined to GABAergic neurons, and *Foxp1* transcript exceeds FOXP1 protein in *Sox14*+ neurons.** Markers were analyzed at mid rostro-caudal levels of the mouse tVTA. For each panel: left, proportion of double-labeled and single-labeled cells *per* section (one dot *per* section; lines, median and quartiles); middle, pooled co-expression across sections shown as pie charts**;** right, representative confocal image (the two probed markers in magenta and yellow, as labeled on each panel), in which filled arrows indicate double-labeled cells and open arrows single-labeled cells. Co-expression is given below as mean ± s.e.m. *per* section; the values across animals are displayed in the Results. **a** *Gad1*/*Gad2* (encoding glutamate decarboxylases): *Gad1* was detected in 96.8±0.8% of *Gad2*+ neurons, and *Gad2* in 92.2±0.7% of *Gad1*+ neurons (*n* = 12 sections, 3 mice, 1,690 cells). **b** *Gad1*/*Oprm1*: *Gad1* was detected in 96.7±0.8% of *Oprm1*+ neurons, and *Oprm1* encoding the mu-opioid receptor in 74.3±4.2% of *Gad1*+ neurons (*n* = 8 sections, 3 mice, 1,184 cells). **c** *Sox14*/*Foxp1* mRNA (same data as in **Fig. 3h),** shown alongside **d** to compare *Foxp1* transcript and FOXP1 protein detection within *Sox14*+ neurons: *Foxp1* was detected in 93.7±1.0% of *Sox14*+ neurons, and *Sox14* in 86.7±2.5% of *Foxp1*+ neurons (*n* = 12 sections, 3 mice, 1,459 cells). **d** *Sox14*/FOXP1: FOXP1 protein detected by immunohistochemistry was combined with *Sox14 in situ* hybridization. FOXP1 was detected in 56.7±2.5% of *Sox14*+ neurons, and *Sox14* in 99.3±0.4% of FOXP1+ neurons (*n* = 12 sections, 3 mice, 2,680 cells). Scale bar: 30 µm.

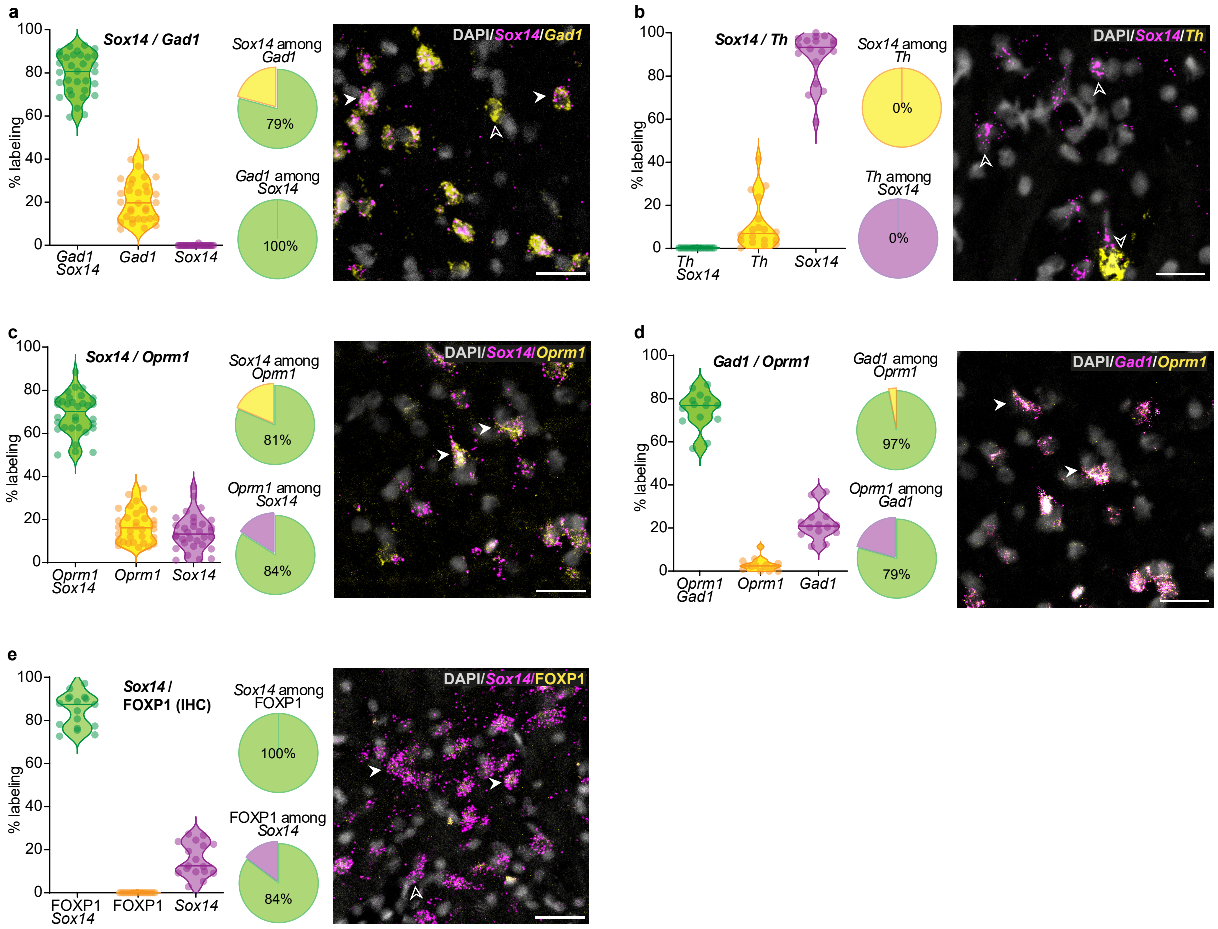


**Supplementary Figure 4 | The GABAergic and opioid profile of *Sox14*+ tVTA neurons is conserved in the rat.** Markers were analyzed at mid rostro-caudal levels of the rat tVTA. For each panel: left, proportion of double-labeled and single-labeled cells per section (one dot per section; lines, median and quartiles); middle, pooled co-expression across sections shown as pie charts; right, representative confocal image (the two probed markers in magenta and yellow, as labeled on each panel), in which filled arrows indicate double-labeled cells and open arrows single-labeled cells. Co-expression is given below as mean ± s.e.m. *per* section; the values across animals are displayed in the Results. **a** *Sox14*/*Gad1*: *Gad1* was detected in every *Sox14*+ neuron (100%), and *Sox14* in 79.2±1.7% of *Gad1*+ neurons (*n* = 32 sections, 6 rats, 2,559 cells). **b** *Sox14*/*Th*: no double-labeled cell was detected in any section (*n* = 18 sections, 3 rats, 1,105 cells). **c** *Sox14*/*Oprm1*: *Oprm1* was detected in 83.2±1.7% of *Sox14*+ neurons, and *Sox14* in 80.2±1.5% of *Oprm1*+ neurons (*n* = 32 sections, 6 rats, 2,287 cells). d *Gad1*/*Oprm1*: *Gad1* was detected in 96.1±1.1% of *Oprm1*+ neurons, and *Oprm1* in 77.0±2.1% of *Gad1*+ neurons (*n* = 14 sections, 3 rats, 1,114 cells). **e** *Sox14*/FOXP1: FOXP1 protein detected by immunohistochemistry was combined with *Sox14 in situ* hybridization. FOXP1 was detected in 84.5±1.9% of *Sox14*+ neurons, and *Sox14* in every FOXP1+ neuron (100%) (*n* = 17 sections, 3 rats, 2,213 cells). Scale bar: 30 µm.

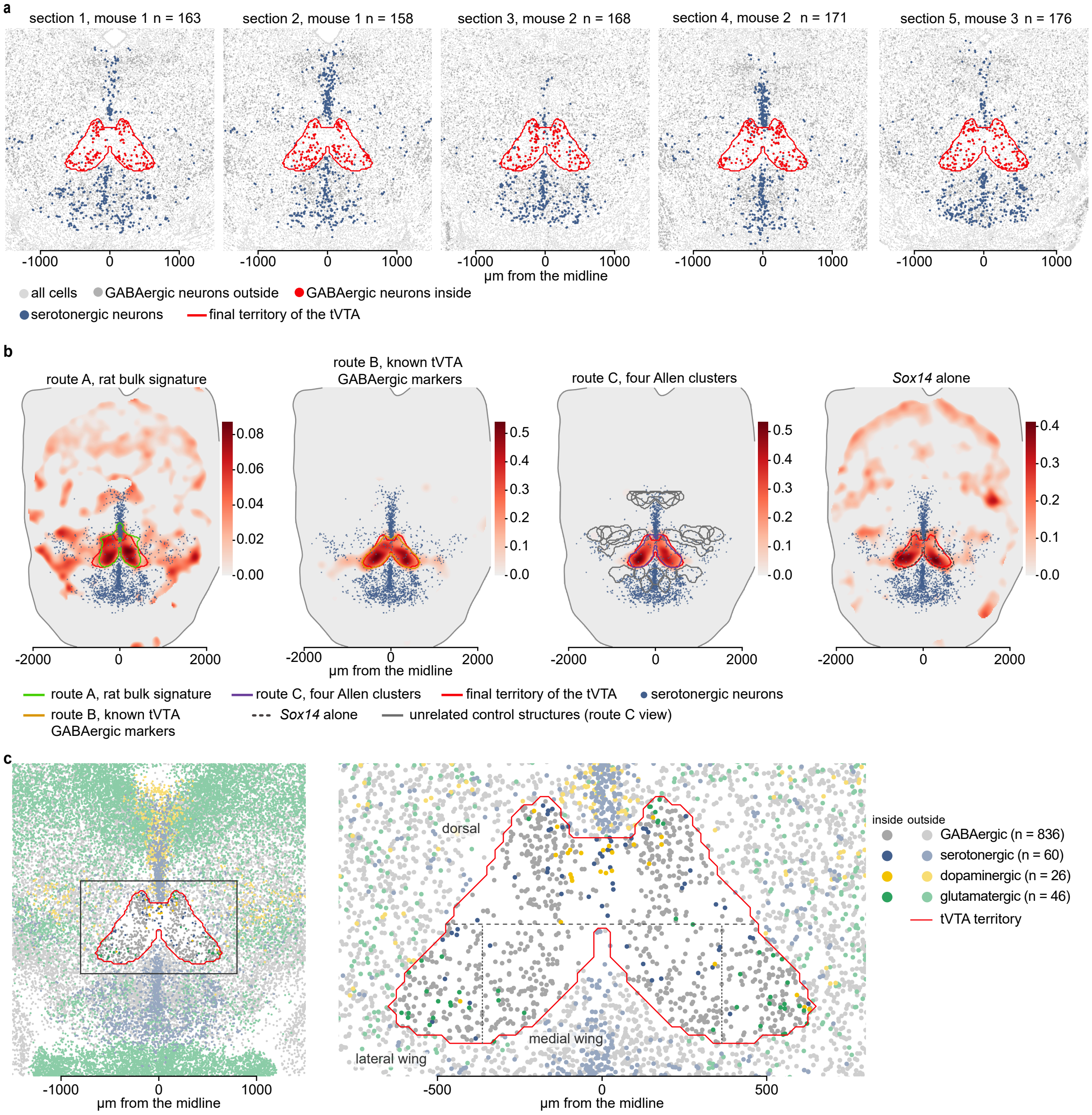


**Supplementary Figure 5 | Delineation and general composition of the tVTA.** **a** Same final tVTA territory (red contour) shown on each of the individual Xenium sections (1 and 2, mouse 1; 3 and 4, mouse 2; 5, mouse 3). *n*, GABAergic neurons inside tVTA territory *per* section (836 in total). On every section and mouse, the contour enclosed a similar number of tVTA GABAergic neurons (158 to 176). **b** One map *per* route, on representations pooling the five tVTA sections. Each heat map shows the local relief of the route score or of the proportion of expressing cells among GABAergic neurons (red scale), the route contour, and the final tVTA territory contour (red). Route A, rat bulk signature; route B, known tVTA GABAergic markers; route C, neuron area of the four main Allen clusters identified in the tVTA (thin gray lines, GABAergic neurons of thirteen unrelated Allen clusters); *Sox14* alone, proportion of *Sox14*-expressing GABAergic neurons. Each heat map peaked at the same location. **c** Map of neurons by class in and around the tVTA, on a representation pooling the five tVTA sections (right, view of the framed area; *n*, neurons inside the tVTA). Dashed line, limit between the dorsal compartment and the ventral wings; dotted lines, limit between the medial and lateral wings**.** Within the tVTA, sparse monoaminergic and glutamatergic neurons lay mostly in the dorsal compartment (next to the dorsal raphe) and lateral wings respectively.

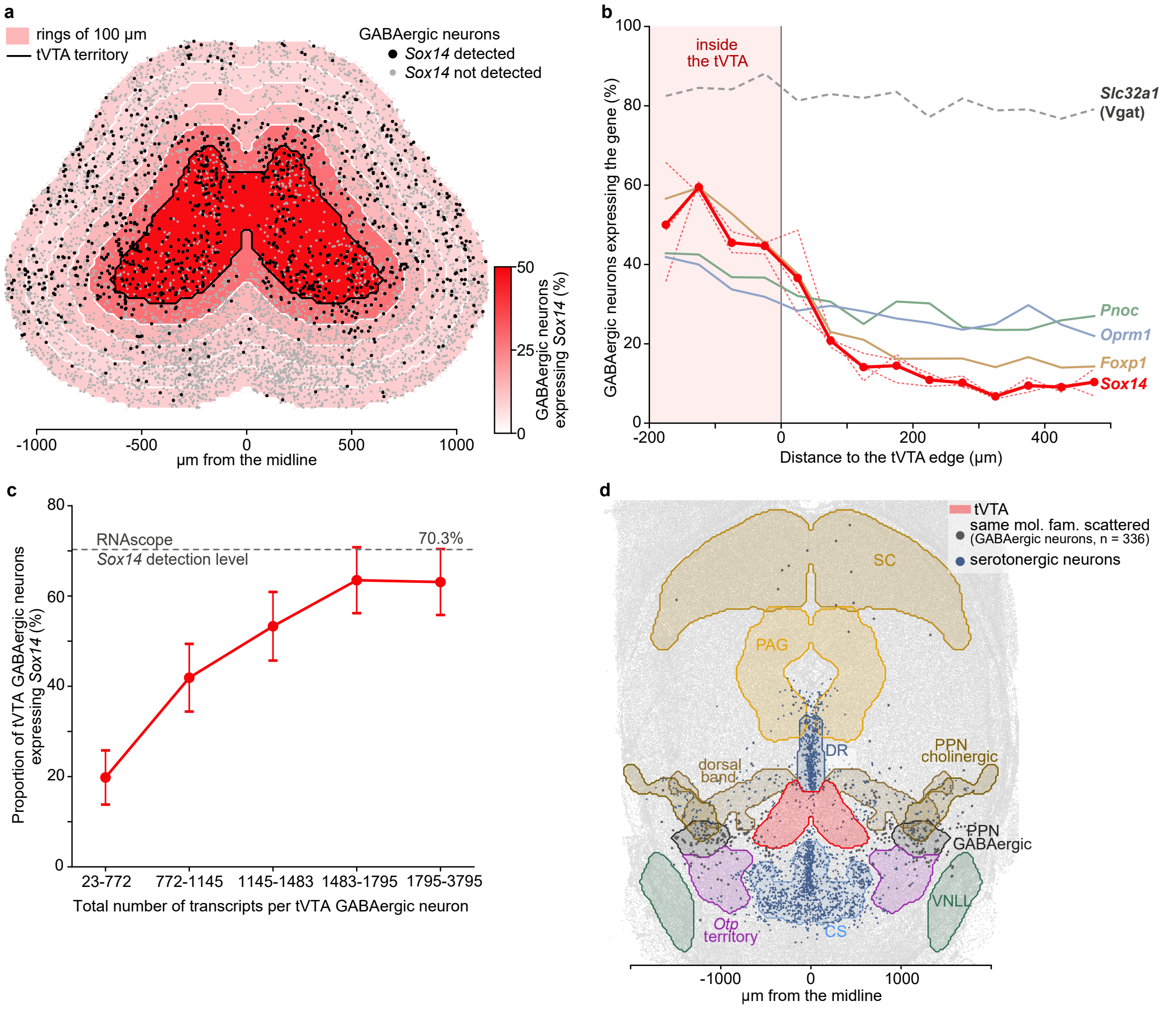


**Supplementary Figure 6 | *Sox14* expression concentrates in the tVTA and drops at its edge.** **a** Proportion of GABAergic neurons expressing *Sox14* (color, mean of three mice) in the tVTA final territory and in five 100 µm rings around its edge, five Xenium sections pooled. Dots, GABAergic neurons. *Sox14*-expressing neurons clustered in the two wings of the tVTA and thinned out from the first ring. **b** Gene expression profile across the tVTA edge. Proportion of GABAergic neurons expressing each gene by 50 µm distance to the edge of the final territory. Thick lines, mean of three mice; thin dashed red lines, each mouse for *Sox14*; dashed gray, *Slc32a1*, reference. *Sox14* and *Foxp1* dropped by half across the edge, whereas *Pnoc* and *Oprm1* decreased by less than a fifth. Beyond 300 µm, *Sox14* was the least frequent of these genes. **c** *Sox14* detection against total transcripts *per* neuron in Xenium experiment. The 836 tVTA GABAergic neurons (five sections, three mice pooled) were sorted into five groups of equal size by their total number of transcripts (x-axis)**.** The graph displays the proportion and 95% binomial interval. *Sox14* detection rose from 20% in the neurons with the fewest transcripts to 63% in the two upper fifths, close to the RNAscope value in *Gad1*+ neurons (dashed line). **d** Anatomical location of tVTA neighboring structures on a representation merging the five tVTA Xenium sections. Each of nine structures was drawn as the zone of highest concentration of its identity cells: serotonergic neurons (dorsal and median raphe, DR and CS), Allen cluster 4701 (GABAergic pedunculopontine nucleus, PPN), Allen cluster 3435 (dorsal band), cholinergic neurons (cholinergic PPN), *Slc6a5*-expressing neurons (ventral nucleus of the lateral lemniscus, VNLL), Allen subclass labels (superior colliculus, SC; periaqueductal gray, PAG**)**, *Otp*-expressing neurons (*Otp* territory). The tenth population, scattered GABAergic neurons of the Allen subclass 268 outside the tVTA and its clusters, is shown as dots (same mol. fam., scattered).

| | tVTA | ring 0-100 µm | ring 100-200 µm | ring 200-300 µm | ring 300-400 µm | ring 400-500 µm | dorsal raphe, 12 µm | dorsal band, 12 µm | median raphe, 94 µm | Otp territory, 129 µm | PPN GABAergic, 262 µm | PPN cholinergic, 394 µm | periaqueductal gray, 468 µm | VNLL, 896 µm | superior colliculus, 1901 µm | same mol. fam., scattered |
|---|---|---|---|---|---|---|---|---|---|---|---|---|---|---|---|---|
| candidate genes | | | | | | | | | | | | | | | | |
| *Foxp1* | 52 | 30 | 19 | 16 | 15 | 14 | 20 | 17 | 13 | 16 | 18 | 14 | 14 | 32 | 17 | 37 |
| *Sox14* | 48 | 28 | 14 | 11 | 8 | 10 | 13 | 20 | 3 | 3 | 12 | 10 | 16 | 5 | 40 | 50 |
| *Sema3e* | 24 | 15 | 7 | 4 | 4 | 4 | 4 | 5 | 4 | 4 | 7 | 5 | 5 | 5 | 3 | 37 |
| *Tmem132c* | 23 | 11 | 5 | 4 | 4 | 6 | 3 | 6 | 3 | 3 | 12 | 12 | 7 | 4 | 8 | 27 |
| RNAscope markers | | | | | | | | | | | | | | | | |
| *Pnoc* | 38 | 31 | 28 | 27 | 23 | 27 | 33 | 33 | 17 | 32 | 29 | 19 | 21 | 12 | 12 | 17 |
| *Oprm1* | 35 | 29 | 27 | 24 | 27 | 23 | 16 | 30 | 19 | 28 | 29 | 25 | 10 | 8 | 10 | 32 |
| *Sstr1* | 27 | 15 | 10 | 6 | 9 | 8 | 7 | 12 | 6 | 9 | 15 | 16 | 11 | 2 | 7 | 22 |
| *Gad1* | 55 | 50 | 47 | 45 | 43 | 44 | 43 | 44 | 35 | 48 | 47 | 39 | 38 | 16 | 52 | 59 |
| *Gad2* | 72 | 69 | 67 | 67 | 68 | 67 | 64 | 76 | 57 | 66 | 75 | 64 | 64 | 31 | 65 | 80 |
| *Th* | 2 | 3 | 2 | 2 | 1 | 2 | 4 | 3 | 3 | 0.2 | 1 | 2 | 0.5 | 0.6 | 0.1 | 0.6 |
| bulk top 20 | | | | | | | | | | | | | | | | |
| *Pax7* | 2 | 9 | 9 | 3 | 3 | 2 | 0.6 | 4 | 8 | 0.8 | 17 | 19 | 8 | 0 | 24 | 1 |
| *Cryba2* | 0.5 | 0.5 | 0.3 | 0.6 | 0.3 | 0.3 | 0.4 | 0.4 | 0.5 | <0.1 | 0.1 | <0.1 | 0.2 | 2 | <0.1 | 0.3 |
| *Otp* | 3 | 9 | 14 | 18 | 16 | 13 | 0.5 | 8 | 20 | 25 | 20 | 16 | 0.1 | <0.1 | <0.1 | 4 |
| *Gata3* | 30 | 22 | 18 | 15 | 12 | 16 | 20 | 23 | 8 | 5 | 7 | 10 | 28 | 23 | 26 | 28 |
| *En1* | 43 | 27 | 22 | 18 | 18 | 18 | 34 | 34 | 4 | 7 | 30 | 32 | 8 | 11 | 0.3 | 50 |
| *Irx6* | 3 | 1 | 0.9 | 0.7 | 0.3 | 0.7 | 0.9 | 1 | 0.2 | 0.7 | 2 | 0.8 | 0.3 | 0.3 | 0.5 | 2 |
| *Glp1r* | 0.4 | 0.4 | 0.8 | 0.7 | 0.9 | 0.7 | 0.2 | 0.3 | 1 | 1 | 0.6 | 0.2 | 0.6 | 3 | 0.6 | 0.4 |
| *Irx3* | 33 | 20 | 19 | 19 | 15 | 17 | 11 | 16 | 15 | 21 | 23 | 11 | 3 | 2 | 4 | 20 |
| *Irx5* | 16 | 11 | 10 | 10 | 9 | 8 | 4 | 8 | 11 | 10 | 10 | 5 | 2 | 1 | 3 | 10 |
| monoaminergic (bulk) | | | | | | | | | | | | | | | | |
| *Slc6a3* | 2 | 3 | 2 | 1 | 1 | 2 | 4 | 5 | 0.2 | 0.4 | 0.9 | 2 | 0.5 | 0.3 | 0.2 | 0.5 |
| *Tph2* | 2 | 4 | 5 | 5 | 4 | 4 | 13 | 3 | 7 | 2 | 1 | 0.6 | 2 | 0.7 | 0.5 | 2 |
| *Slc18a2* | 3 | 5 | 5 | 6 | 5 | 6 | 11 | 5 | 10 | 4 | 3 | 2 | 3 | 2 | 1 | 4 |
| control | | | | | | | | | | | | | | | | |
| *Slc32a1* | 86 | 82 | 83 | 80 | 79 | 78 | 70 | 81 | 68 | 82 | 83 | 70 | 70 | 52 | 74 | 91 |

relative to the tVTA (log scale): 8, 4, 2, 1, 1/2, 1/4, 1/8

**Supplementary Figure 7 | The candidate genes are expressed more often in the tVTA than in neighboring areas.** Number, percentage of the GABAergic neurons of the tVTA (first column), of its five 100 µm rings and of neighboring structures, that express the gene; color, this percentage relative to that of the tVTA (log scale). Neighboring structures as in **Supplementary Fig. 6d**, ordered by distance to tVTA edge. Rows, candidate genes selected from the Xenium analysis, RNAscope markers (**Fig. 3**), selected other genes of the bulk top 20 present in the Xenium panel and monoaminergic markers (**Fig. 1c**), and *Slc32a1* as a control. The candidate genes were less frequent in every ring and region than in the tVTA, except *Sox14*, *Sema3e* and *Tmem132c* in the scattered neurons of the same molecular family. *Pax7* and *Otp*, among the most enriched genes of the bulk, were expressed by 2 and 3% of tVTA GABAergic neurons and more often around the tVTA. Five tVTA sections from three mice, mean of the three mice. mol. fam., molecular family; PPN, pedunculopontine nucleus; VNLL, ventral nucleus of the lateral lemniscus.

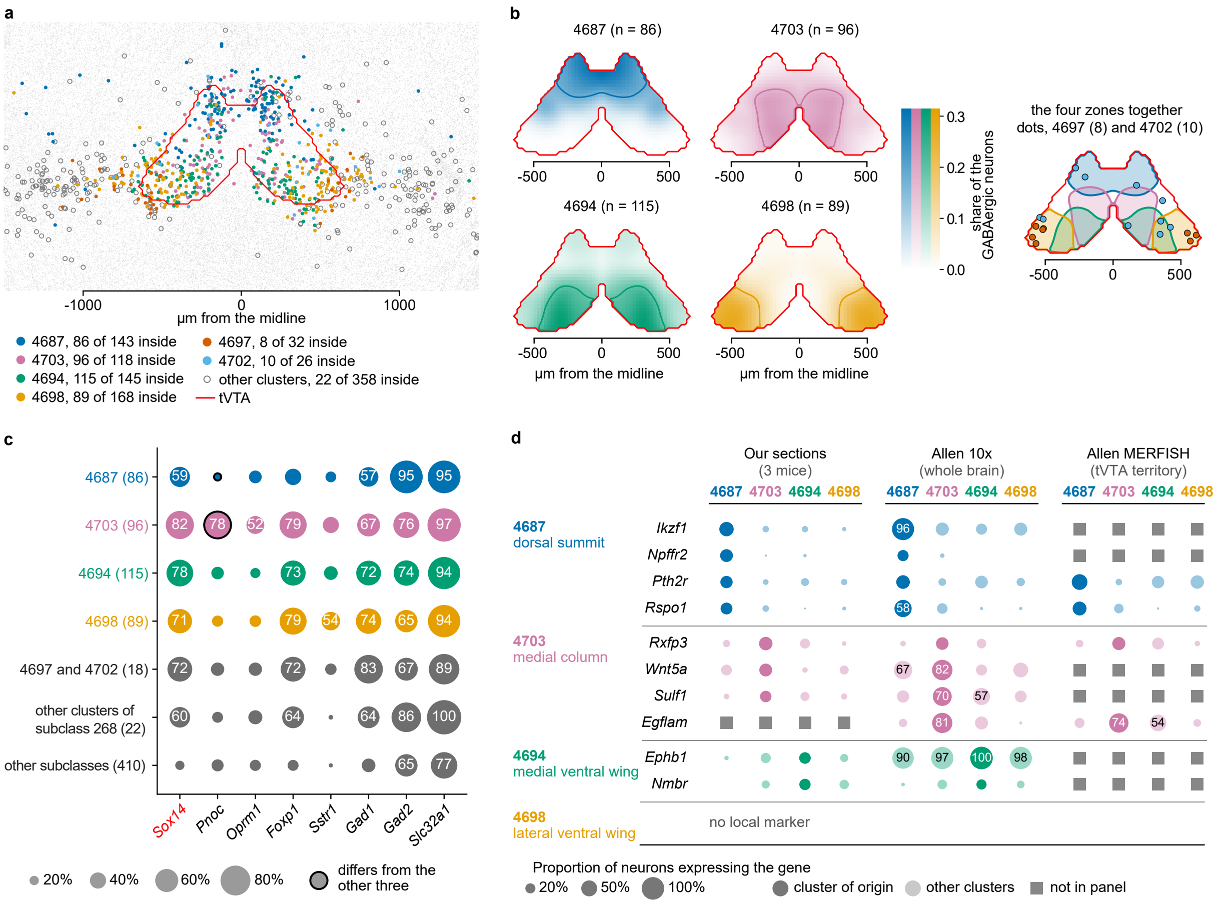


**Supplementary Figure 8 | Molecular subpopulations of the tVTA neurons are defined by Allen taxonomy. a** The eighteen GABAergic clusters of Allen subclass 268 on the merged representation of the five tVTA sections. Colors, the six Allen clusters of the tVTA, which have at least a quarter of their neurons inside the territory; open circles, the twelve other clusters of the subclass; numbers, neurons inside the tVTA among those of the cluster on the sections. The six clusters lay in and around the tVTA, the other clusters outside it. **b** Zone of highest concentration of the four largest Allen clusters in the tVTA. Representation of the five tVTA sections merged; color, local proportion of the GABAergic neurons that belong to the cluster; *n*, neurons of the cluster in the tVTA; contour, zone of highest concentration of the cluster; right map, the four zones together, with the neurons of the two small clusters, 4697 (8) and 4702 (10), as dots, too few to map; red line, edge of the tVTA. The four clusters were intermingled but topographically ordered, 4687 at the dorsal summit, 4703 in a medial column, 4694 and 4698 in the medial and lateral ventral wings. **c** Proportion of the GABAergic neurons of each cluster that express *Sox14*, the RNAscope markers of **Fig. 3** and *Slc32a1* (Vgat) as a control, three mice pooled**;** *n* in brackets; gray rows, the other GABAergic neurons of the tVTA; dot diameter indicates the proportion, value written above 50%; black edge, proportion differing from the other three large clusters. *Sox14* was expressed by 59 to 82% of the neurons of the four large clusters without distinguishing them; only *Pnoc* differed between clusters. **d** Local markers of **Fig. 4f** in the four large Allen clusters. Circle area, proportion of the cluster neurons expressing the gene in Xenium sections (three mice), in the Allen 10x cells of the cluster (whole brain) and in the Allen MERFISH atlas (neurons of the cluster inside the tVTA territory traced on the atlas, one brain); full color, cluster in which the marker was identified; values written above 50%. In Xenium sections, each marker was most frequent in its own cluster; in the Allen 10x cells, several were also frequent in other clusters (*Ephb1*, 90 to 100% in the four clusters).

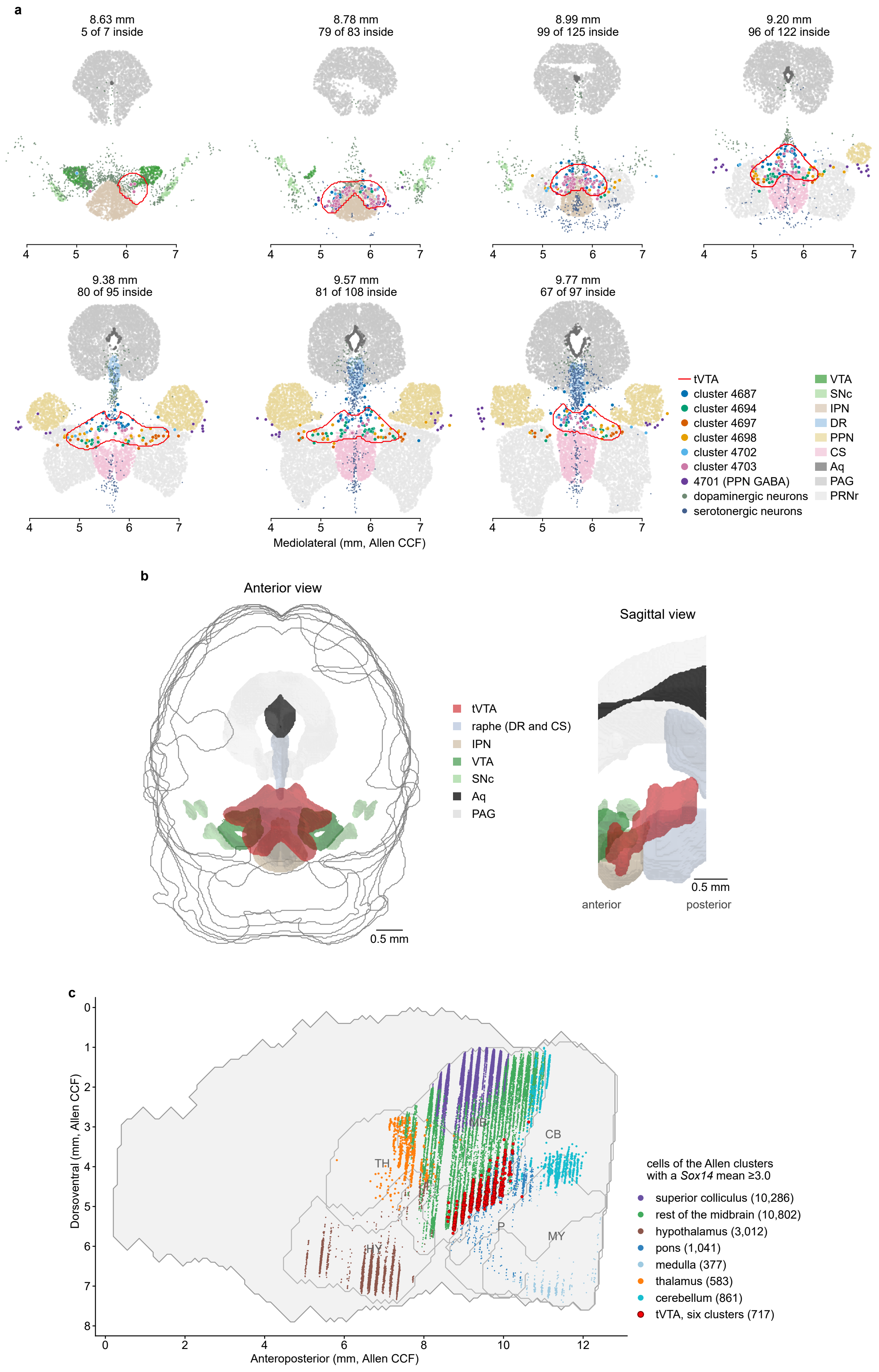
a
8.63 mm
5 of 7 inside
8.78 mm
79 of 83 inside
8.99 mm
99 of 125 inside
9.20 mm
96 of 122 inside
9.38 mm
80 of 95 inside
9.57 mm
81 of 108 inside
9.77 mm
67 of 97 inside
Mediolateral (mm, Allen CCF)
tVTA
cluster 4687
cluster 4694
cluster 4697
cluster 4698
cluster 4702
cluster 4703
4701 (PPN GABA)
dopaminergic neurons
serotonergic neurons
VTA
SNc
IPN
DR
PPN
CS
Aq
PAG
PRNr
b
Anterior view
Sagittal view
tVTA
raphe (DR and CS)
IPN
VTA
SNc
Aq
PAG
0.5 mm
anterior
posterior
0.5 mm
c
Dorsoventral (mm, Allen CCF)
Anteroposterior (mm, Allen CCF)
TH
MB
CB
HY
P
MY
cells of the Allen clusters
with a Sox14 mean ≥3.0
superior colliculus (10,286)
rest of the midbrain (10,802)
hypothalamus (3,012)
pons (1,041)
medulla (377)
thalamus (583)
cerebellum (861)
tVTA, six clusters (717)

**Supplementary Figure 9 | The six tVTA clusters form a compact volume in the Allen MERFISH atlas.** **a** Representation of the seven coronal sections, from anterior to posterior, of the Allen MERFISH atlas[35] on which the tVTA territory was traced (red line) where cells of its six Allen clusters (colored dots) were most concentrated. Above each section, position in mm on the anteroposterior axis of the Allen Common Coordinate Framework (CCF)[30]and number of cells of the six clusters inside the territory among those on the section. Purple, Allen cluster 4701 (GABAergic PPN) of the same subclass 268 but outside the tVTA; gray-green, dopaminergic neurons; blue, serotonergic neurons; light colored background, cells of each CCF structure. Despite the limited anatomical precision of the Allen MERFISH atlas, the tVTA territory observed through this strategy began in the caudal VTA, above the IPN, and ended between the raphe and the PPN, with a shape similar to our RNAscope and Xenium characterizations. **b** Three-dimensional representation, anterior and sagittal views, of the extent of the tVTA in the Allen brain atlas, built from the territory in **a**, traced on each section from the six Allen clusters identified from our Xenium sections. Each section was placed on its own plane in the CCF and the volume interpolated between sections, with the CCF structures of the same anteroposterior window (8.4 to 10.0 mm). The tVTA according to this representation measured 0.78 $mm^3$ over 1.14 mm, starting in the caudal VTA, above the IPN, in front, to end between the raphe nuclei behind. Scale bars, 0.5 mm. **c** Sagittal projection of *Sox14*-rich clusters (Allen clusters with a mean *Sox14* expression of at least 3.0 on the Allen log scale) on the Allen MERFISH atlas; *n* of cells in brackets. The six tVTA clusters held 717 of the 30,255 cells; the 2,576 cells outside the listed divisions were not drawn. Outside the tVTA, most *Sox14*-rich clusters lay more than 1 mm away. Aq, cerebral aqueduct; CB, cerebellum; CCF, Allen Common Coordinate Framework; CS, superior central nucleus (median raphe); DR, dorsal raphe; HY, hypothalamus**;** IPN, interpeduncular nucleus; MB, midbrain; MY, medulla; P, pons; PAG, periaqueductal gray; PPN, pedunculopontine nucleus; PRNr, pontine reticular nucleus, rostral part; SNc, substantia nigra pars compacta; TH, thalamus; VTA, ventral tegmental area.

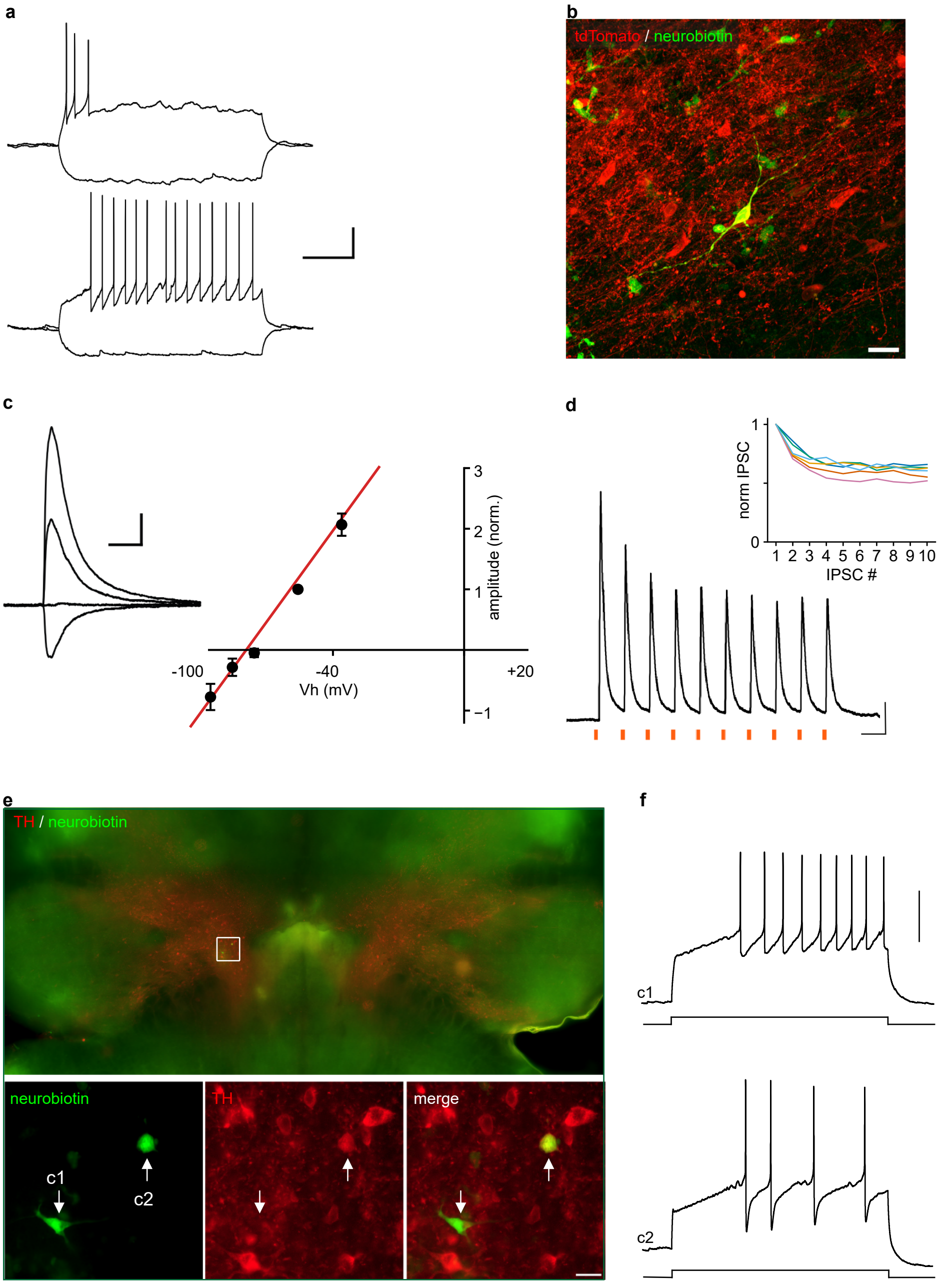


**Supplementary Figure 10 | Supporting data for the slice recordings of *Sox14*+ tVTA neurons and VTA neurons.** **a** Whole-cell current-clamp recordings from two representative *Sox14*+ tVTA neurons, responses to current steps of -20 and +20 pA (top) and -20 and +40 pA (bottom). Top: a short burst at step onset, as in five of the fourteen neurons tested; bottom: irregular spiking throughout the step, as in eight of the fourteen neurons tested, one neuron firing a single spike. The sixteen neurons recorded in whole-cell had a membrane resistance of 636±72 MΩ and a capacitance of 13.5±1.1 pF. **b** Example of a neurobiotin-filled *Sox14*+ tVTA neuron (green) among tdTomato-expressing neurons (red); a fusiform soma with two or three fine processes was observed, as in most of the nine neurons recovered. **c** Current-voltage relation of the light-evoked inhibitory postsynaptic current (IPSC), each amplitude normalized to that of the same neuron at -56 mV; means ± s.e.m from seven neurons; red line: the linear fit. The current reversed at -78 mV, close to the chloride equilibrium potential (ECl = -89 mV). Inset: IPSCs of one neuron at -96, -76, -56 and -36 mV. **d** Train of ten IPSCs evoked at 20 Hz in a representative VTA neuron. Inset: amplitude of each IPSC normalized to the first, in the six neurons tested, one color *per* neuron. **e** Localization in a horizontal slice (within the white square) of two responsive VTA neurons. Below, zoom on the boxed area. The two neurobiotin-filled neurons were labeled with Alexa Fluor 488-coupled streptavidin (green) and tyrosine hydroxylase (TH) immunolabeling (red). Cell c1 is TH-negative, and cell c2 is TH-positive. **f** Whole-cell current-clamp responses of the same two neurons as in **e** evoked by injection of a 1 s positive current step. Each cell was held around -60 mV. Note the slow initial rising phase delaying the first action potential, as typical of VTA dopaminergic neurons. Values, mean ± s.e.m. Scale bars: 20 mV, 0.5 s (**a**); 20 µm (**b**); 100 pA, 10 ms (**c**); 100 pA, 50 ms (**d**); 20 µm (**e**, bottom); 30 mV (**f**).

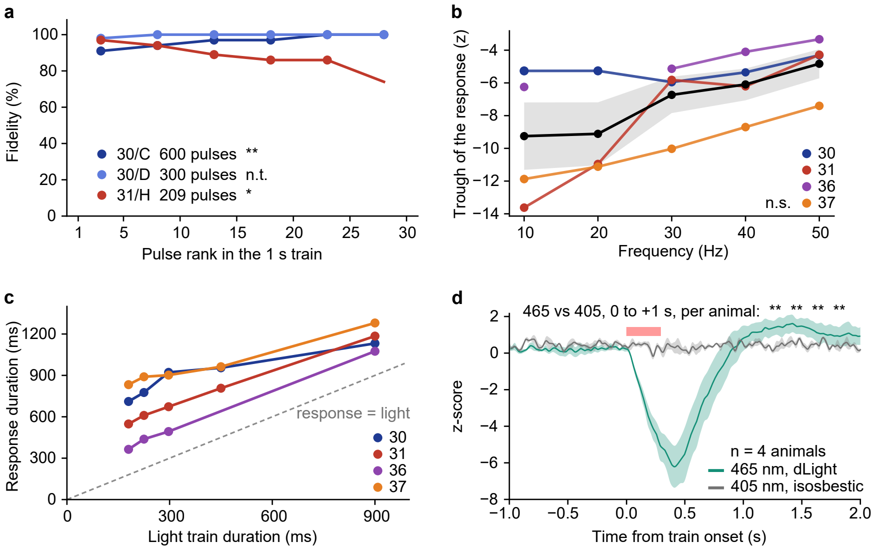


**Supplementary Figure 11 | Supporting data for the photoidentified tVTA neurons and the accumbal dopamine response.** **a** *Sox14*+ tVTA neurons responded to trains of light pulses (957 ms, 30 Hz) with high fidelity (87.6 to 99.7%, three neurons, 1109 pulses). A test of the trend along each train (Wilcoxon signed-rank on a planned linear contrast of the six blocks of five pulses, one contrast *per* train) showed that fidelity rose in one neuron, fell in another without going below 74%, and was not testable in the third, whose trains were answered almost perfectly. **b** Trough of the response against stimulation frequency; one line *per* animal, black the mean of the four. The trough did not vary detectably with frequency ($F_{(2,6)}$ = 4.44, corrected $P$ = 0.12). **c** Duration of the dopamine response, from its onset to its end, against light train duration; one point *per* animal and frequency, one line *per* animal; dashed line, equality. The response outlasted the light in all 19 recordings, and its duration varied with frequency (repeated measures ANOVA over 10, 30 and 50 Hz, $F_{(2,6)}$ = 39.7, Greenhouse-Geisser corrected $P$ = 0.0044), whereas its onset did not ($F_{(2,6)}$ = 0.35, corrected $P$ = 0.62). **d** The isosbestic control (gray line, 405 nm channel) showed that the decrease in extracellular dopamine (green line, 465 nm dLight channel) was not an artifact. Displayed data correspond to the 30 Hz stimulation train (pink bar; Wilcoxon within each animal, paired over its ten trains, $P$ = 0.0020 in three animals and $P$ = 0.0098 in the fourth). In (**a**) each dot is a five-pulse block in one neuron; in **(b, d)** lines are mean and s.e.m. * $P$ < 0.05, ** $P$ < 0.01, *** $P$ < 0.001; n.s., not significant; n.t., not testable. Detailed statistics are provided in **Supplementary Table 11b**.

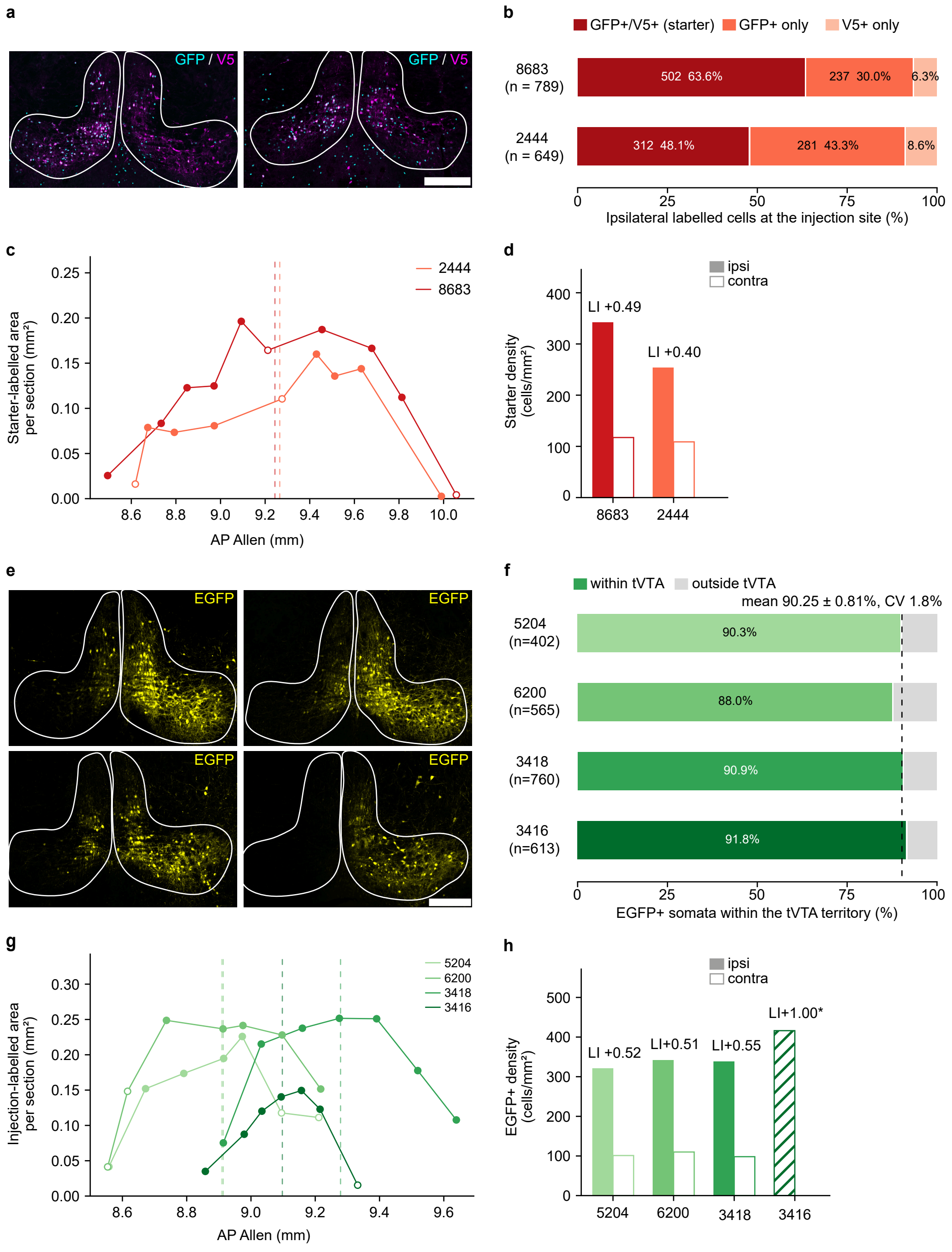


**Supplementary Figure 12 | Validation of the tVTA injection sites for retrograde and anterograde tracing.** **a-d** Monosynaptic rabies tracing, n = 2 mice. **a** Confocal image of the injection site in each mouse: merge of GFP (rabies, cyan) and V5 (helper, magenta). **b** Cellular composition of the ipsilateral injection site: GFP+/V5+ starter cells (*Sox14*+), GFP-only cells (helper-negative local presynaptic partners) and V5-only cells. **c** Starter-labeled area *per* section along the antero-posterior axis, one line *per* mouse. Filled symbols, confocal; open symbols, widefield. Dashed vertical lines, area-weighted center of mass of each mouse. **d** Starter-cell density *per* hemisphere, with the injection laterality index, LI = (ipsilateral - contralateral) / (ipsilateral + contralateral), above each pair. **e-h** Anterograde tracing, n = 4 mice. **e** Confocal images of the injection site in each mouse: EGFP+ somata (yellow, single channel). **f** Proportion of EGFP+ somata inside *versus* outside the tVTA territory (ipsilateral, confocal sections); dashed line, group mean. **g** Injection-labeled area *per* section, conventions as in **c**. **h** EGFP+ somatic density *per* hemisphere, injection LI above each pair, as in **d**. Asterisk, mouse #3416, with no detectable contralateral signal: its index saturates at +1.00. All panels show individual animals. Antero-posterior coordinates, Allen CCFv3. Scale bar: 250 µm in all photomicrographs.

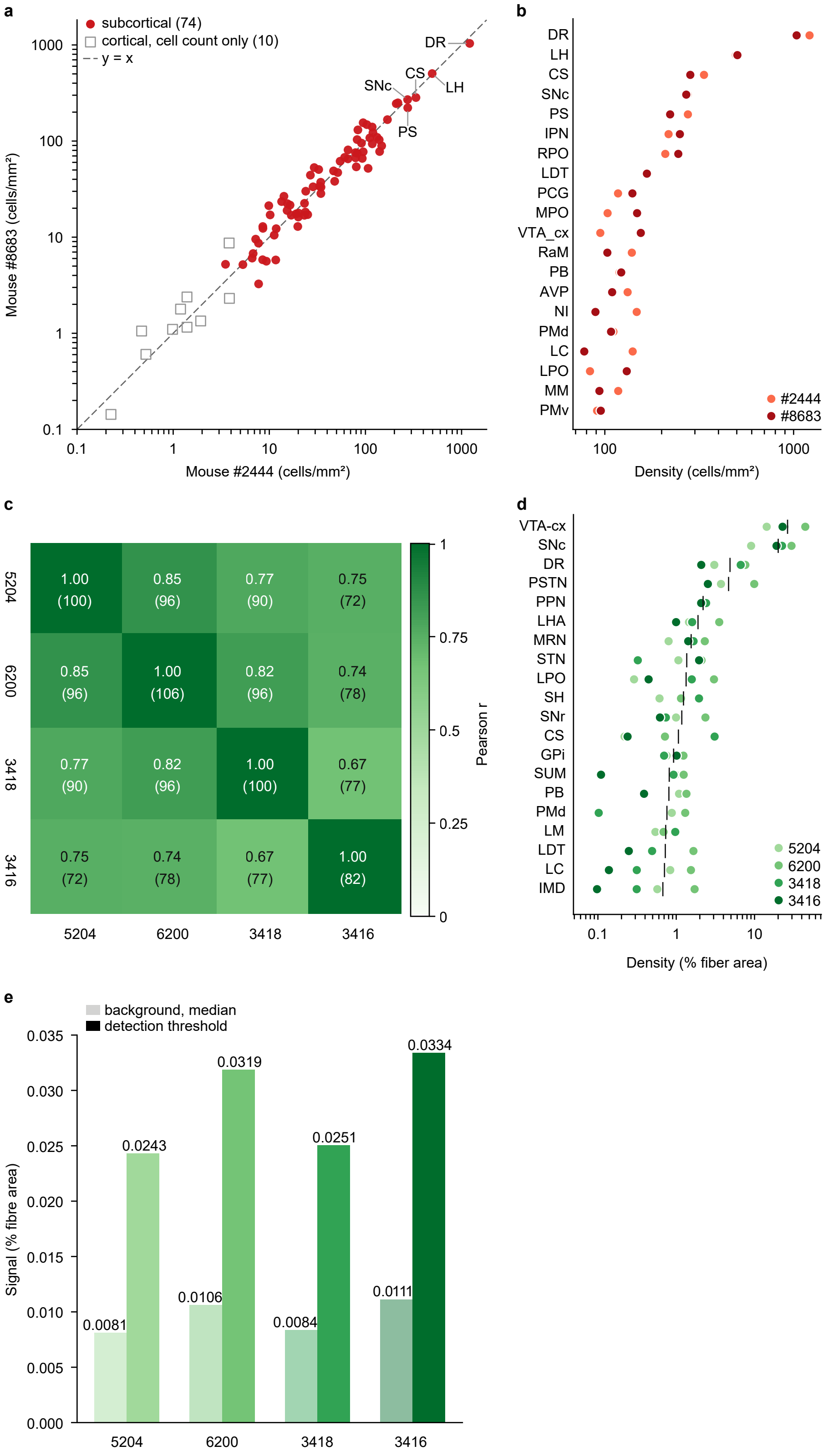


**Supplementary Figure 13 | Inter-animal reproducibility of the *Sox14*+ tVTA connectome.** **a, b** Monosynaptic rabies tracing, n = 2 mice. **a** Labeled-cell density of the afferent regions in mouse #2444 against mouse #8683, log-log. Filled red circles, the 74 subcortical regions on which the correlation is computed, r = 0.96; open gray squares, the 10 isocortical, olfactory and lateral septal regions kept in the connectome for cell count alone but not included in the correlation; dashed line, y = x. Labeled, the five densest regions. **b** The twenty most densely innervated afferent regions, ranked by mean density. Filled circles, the two mice in the red shades of **Fig. 8a**. **c-e** Anterograde tracing, n = 4 mice. **c** Pairwise Pearson correlation of the log of ipsilateral fiber density between the four mice, restricted to the regions above threshold in both animals of a pair. Each cell gives the correlation coefficient (from 0.67 to 0.85) and, in brackets, the number of considered regions (from 72 to 96). Number of regions above threshold in each animal is given in dark green diagonal cells. **d** The twenty densest efferent regions, ranked by mean ipsilateral density; filled circles, the four mice in the green shades of **Fig. 8a**; black tick, group mean. A missing dot means that the region was below detection threshold for that animal, 76 of the 80 possible dots being drawn. **e** For efferent analyses, the detection threshold for fiber density (filled bars) was set for each animal at three times the background (pale bar), defined as the median fiber density across the isocortex, cortical plate and hippocampal region, where tVTA projections are negligible. Nomenclature, Allen Mouse Brain Common Coordinate Framework (CCFv3); full region names, **Supplementary Table 15**. Coefficients of variation and exact *P* values, **Supplementary Table 14 and 11c**.

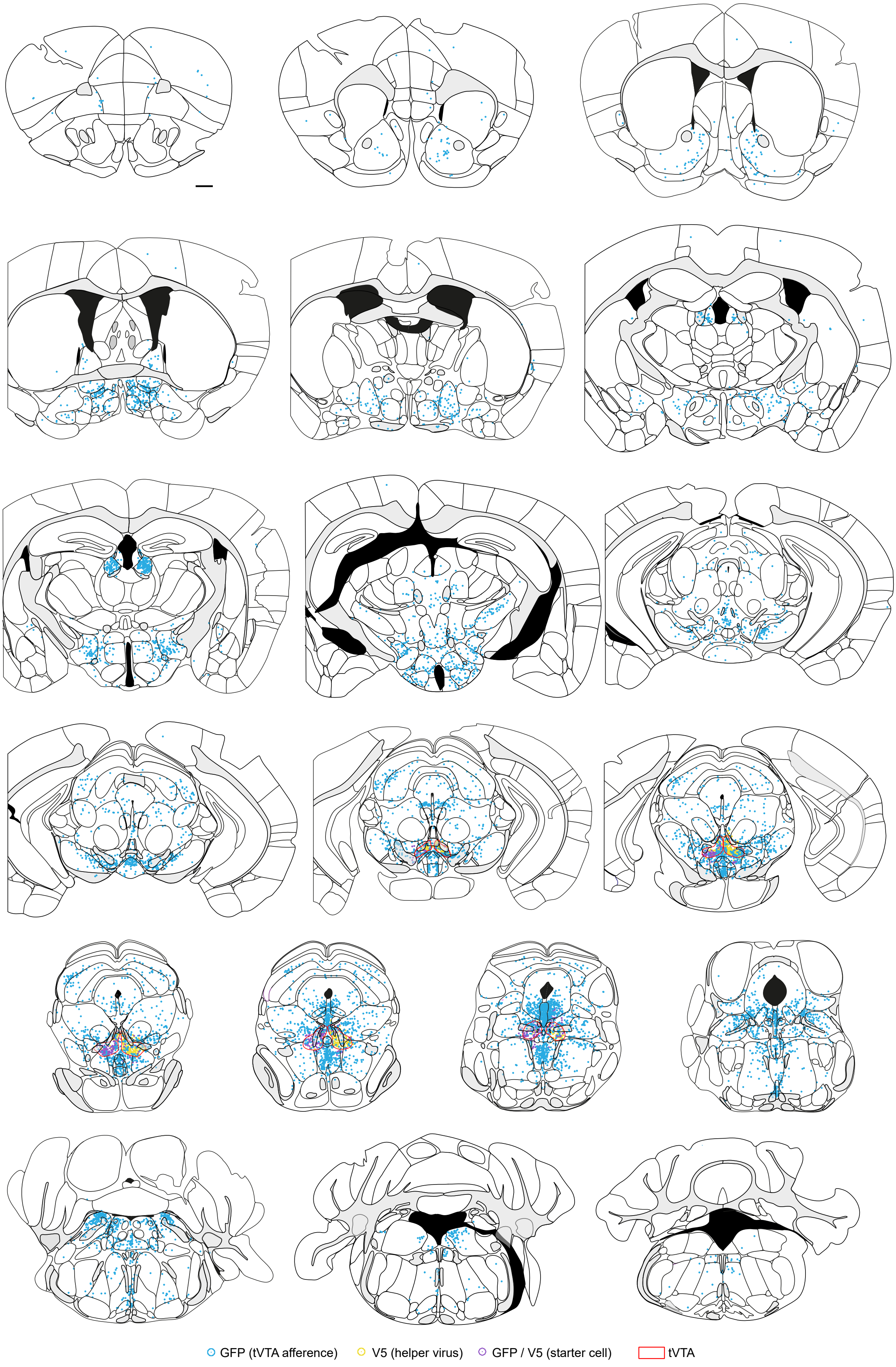
GFP (tVTA afference)
V5 (helper virus)
GFP / V5 (starter cell)
tVTA

**Supplementary Figure 14 | Whole-brain cartography of *Sox14*+ tVTA afferents.** Two-dimensional reconstruction of retrograde labeling from one representative animal (mouse #8683), on 19 serial coronal atlas templates from rostral (top left) to caudal (bottom right). Cyan, GFP+ input neurons; yellow, V5+ helper-expressing cells; purple, GFP+/V5+ starter cells at the injection site; red outline, the tVTA. Sections were chosen to illustrate the rostro-caudal distribution of labeling. This is the section-by-section support for the summarized afferent connectome displayed in **Fig. 8a** and for the representative levels in **Fig. 7c**. Scale bar: 500 µm.

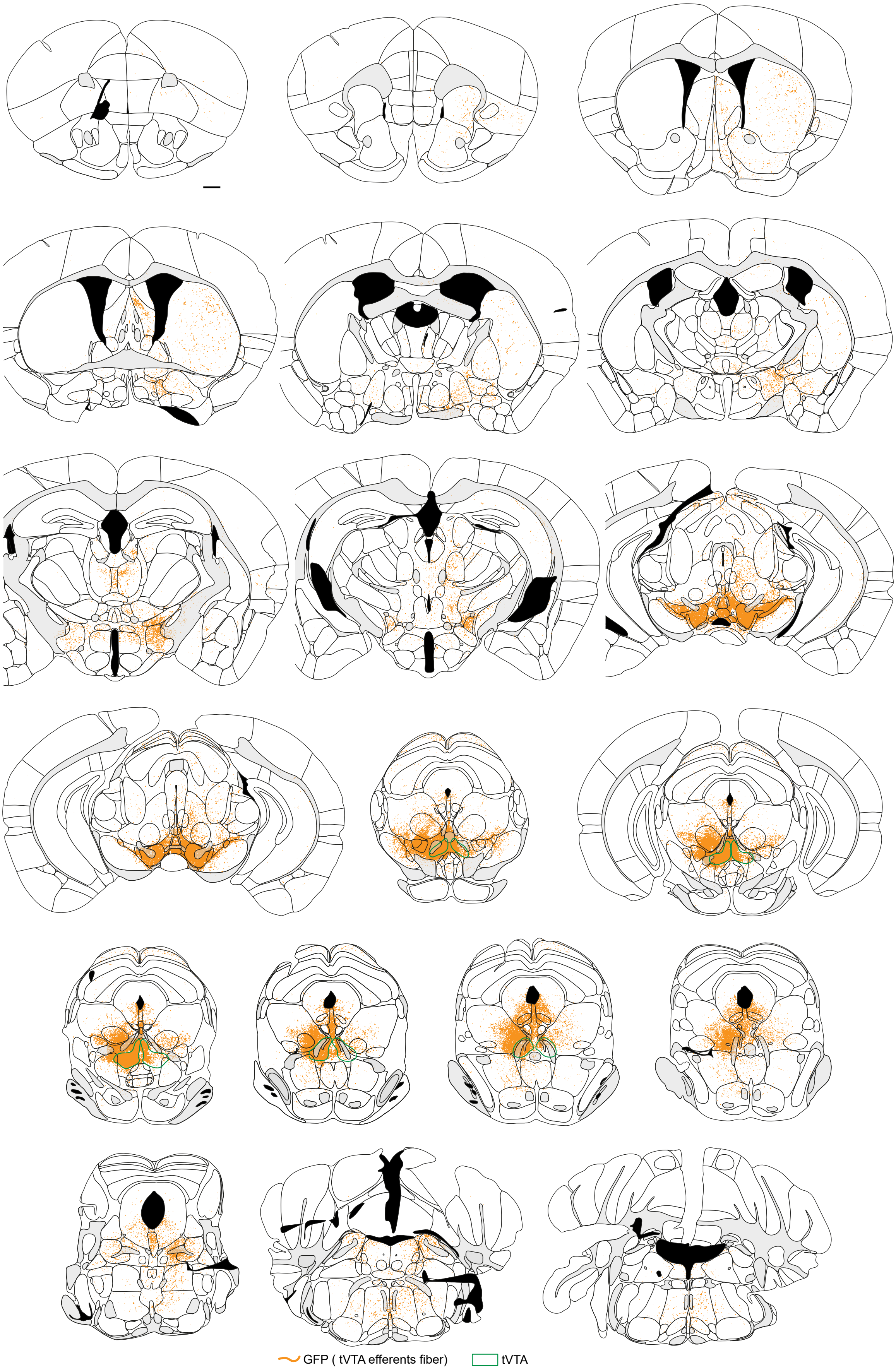

GFP ( tVTA efferents fiber)
tVTA

**Supplementary Figure 15 | Whole-brain cartography of Sox14+ tVTA efferents.** Two-dimensional reconstruction of anterograde labeling from one representative animal (mouse #6200), on 19 serial coronal atlas templates and with the same conventions as **Supplementary Fig. 14**. Orange, EGFP+ fibers; green outline, the tVTA. Sections were chosen to illustrate the rostro-caudal distribution of labeling. This is the section-by-section support for the summarized efferent connectome displayed in **Fig. 7a** and for the representative levels in **Fig. 6d**. Scale bar: 500 µm.

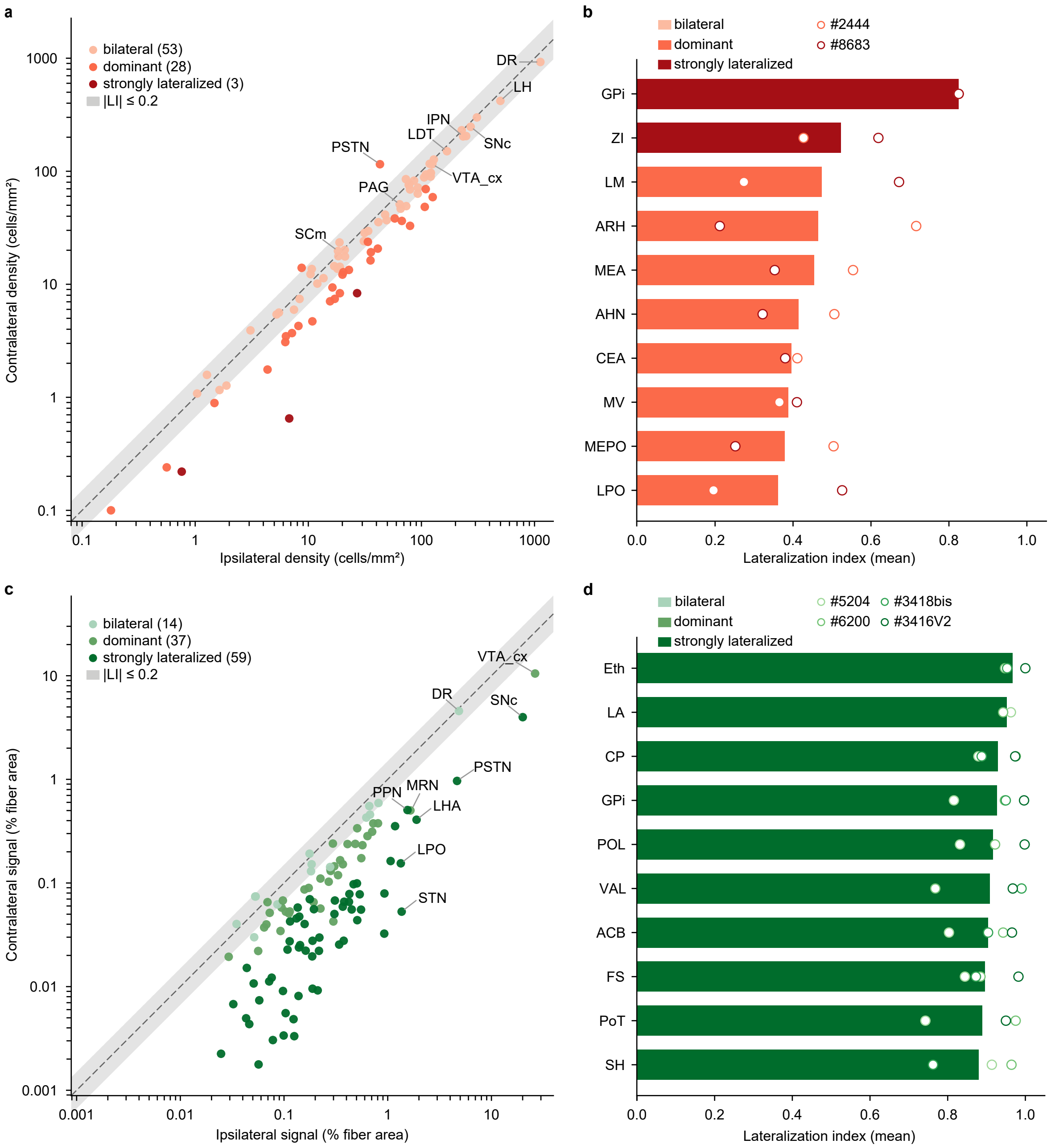


**Supplementary Figure 16 | Hemispheric lateralization of the *Sox14*+ tVTA connectome.** **a** Afferent connectome, n = 2 mice: mean ipsilateral against mean contralateral labeled-cell density, one dot *per* region, log-log. Dot shade indicates bilateral (pale, 53), dominant (medium, 28), and strongly lateralized (dark, 3) regions. Gray area indicates the bilateral range, defined as lateralization index |LI| ≤ 0.2; dashed line, y = x. Labeled, nine regions of interest. Most regions lie in the gray area with predominantly bilateral signal. **b** The ten most ipsilateral afferent regions, ordered by LI; VISC, retained on cell count only, was excluded from this representation. Bars, mean value; open circles, individual mice. **c** Efferent connectome, n = 4 mice: mean ipsilateral against mean contralateral fiber area, averaged over the four animals. Dot shade indicates bilateral (pale, 14), dominant (medium, 37), and strongly lateralized (dark, 59) regions, according to the mean of the *per*-animal indices; gray area and dashed line as in (**a**). Labeled, the nine densest ipsilateral regions. The cloud lies systematically below the diagonal, illustrating the notable ipsilateral bias. **d** As in (**b**), for the efferent connectome; open circles, individual mice. Lateralization index, LI as in **Fig. 8e**; +1, exclusively ipsilateral; 0, balanced. Bilateral, |LI| ≤ 0.2; dominant, |LI| ≤ 0.5; strongly lateralized, |LI| > 0.5. Nomenclature, Allen CCFv3; full region names, **Supplementary Table 15**. Region counts per anatomical super-group in **Supplementary Table 17**.

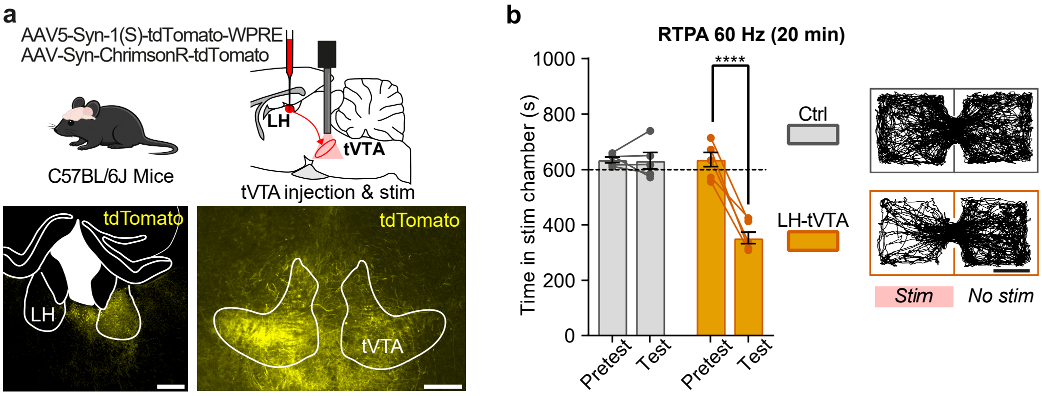


**Supplementary Figure 17 | Optogenetic activation of LH terminals in the tVTA induces behavioral avoidance.** **a** Top, schematic of virus delivery for optogenetic stimulation of terminals of the LH-tVTA pathway in C57BL/6J mice. Bottom, representative images of the LH injection site and of terminal fibers in the tVTA. **b** Real-time place avoidance (RTPA). Left, optogenetic stimulation of LH-tVTA terminals (1 session, 20min, 60Hz, 5ms pulses, uninterrupted) reduced the time spent in the stimulation chamber (Time x Group, $F_{(1,9)} = 44.63$, $P < 0.0001$; pretest *versus* test post-hoc: Ctrl $P = 0.99$, LH-tVTA $P < 0.0001$). Right, representative tracks on the test day. Stimulation parameters as previously reported for this pathway[9]. Group sizes: Ctrl 5, LH-tVTA 6. Graphs display mean ± s.e.m.; dots: individual animals. Detailed analyses of variance and post-hoc comparisons are provided in **Supplementary Table 11d**. Asterisks: pretest *versus* test within a group. ****$P < 0.0001$. Scale bars in a: 300 µm (left) and 200 µm (right).

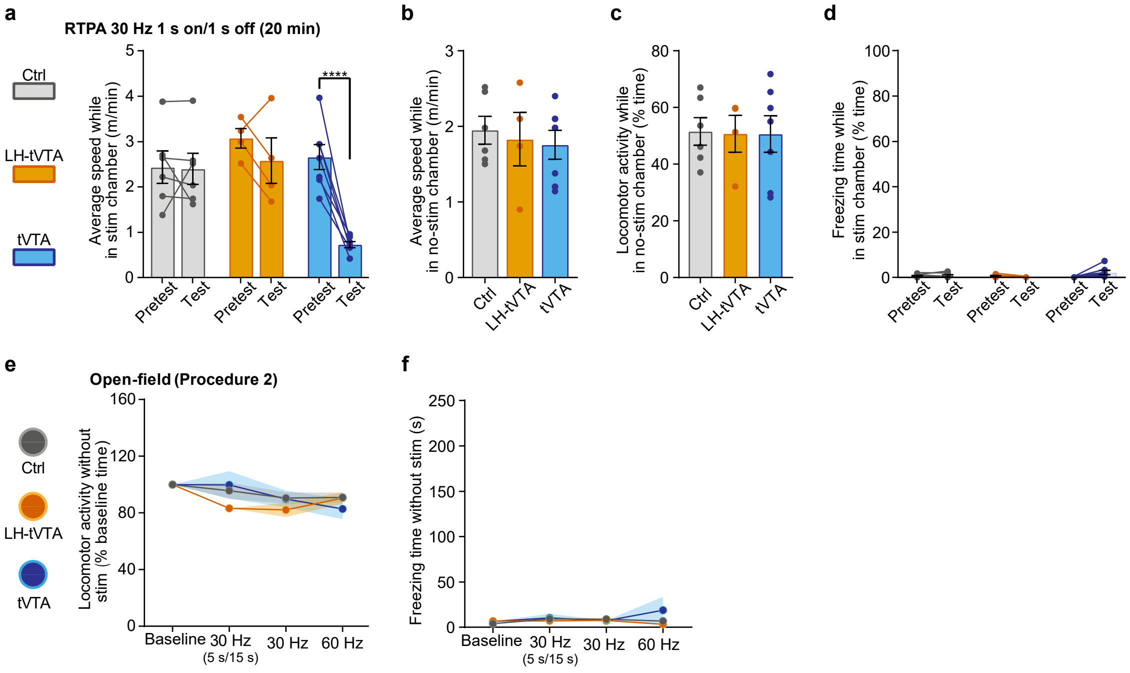


**Supplementary Figure 18 | Locomotor consequences of tVTA activation during the real-time place avoidance and open-field procedures.** **a-d** Real-time place avoidance (RTPA) (1 session, 20 min, 30 Hz, 5 ms pulses, 1 s on/1 s off). **a** Average speed in the stimulation chamber decreased with stimulation of *Sox14*+ tVTA neurons (Time x Group, $F_{(2,14)} = 11.77$, $P = 0.0010$; pretest *versus* test post-hoc: Ctrl $P = 0.99$, LH-tVTA $P = 0.48$, tVTA $P < 0.0001$). **b, c** The average speed and the proportion of time in locomotor activity in the non-stimulated chamber were unaffected on the test day (Speed: $F_{(2,14)} = 0.20$, $P = 0.81$; Locomotor activity: $F_{(2,14)} = 0.0078$, $P = 0.99$)**.** **d** Stimulation of *Sox14*+ tVTA neurons did not induce freezing behavior under this protocol (Time x Group, $F_{(2,14)} = 3.57$, $P = 0.06$). **e, f** Open-field, procedure 2 (4 daily sessions). In the 5 min preceding stimulation, locomotor activity **e** and freezing **f** did not differ between groups or protocols (Protocol x Group, **e** $F_{(6,75)} = 1.59$, $P = 0.16$; **f** $F_{(6,75)} = 1.13$, $P = 0.35$). Group sizes (Ctrl, LH-tVTA, tVTA): 6, 4, 7 Sox14-Cre mice in (**a-d**); 11, 10, 7 in (**e**, **f**) with C57BL/6J and *Sox14*-Cre mice pooled in Ctrl and LH-tVTA groups. Graphs display mean ± s.e.m.; dots in (**a-d**): individual animals. Detailed analyses of variance and post-hoc comparisons are provided in **Supplementary Table 11d**. Asterisks: pretest *versus* test within a group. ****$P < 0.0001$.

**Supplementary Table 1 | Markers of monoaminergic contamination and of oligodendrocytes, with their references.**

| Family | Gene | Ensembl gene ID | Product and localization | Primary reference | Additional reference | Mean TPM, tVTA |
|---|---|---|---|---|---|---|
| dopaminergic | *Slc6a3* | ENSRNOG00000017302 | dopamine transporter, VTA and SNc | Giros et al. 1996[1] | Tiklová et al. 2019[2] | 32.7 |
| dopaminergic | *Th* | ENSRNOG00000020410 | tyrosine hydroxylase, VTA, SNc, A8 to A14 | Hökfelt et al. 1973[3] | Björklund & Dunnett 2007[4] | 121.7 |
| dopaminergic | *Pitx3* | ENSRNOG00000019194 | pituitary homeobox 3, VTA and SNc | Smidt et al. 2004[5] | Veenvliet et al. 2013[6] | 3.6 |
| dopaminergic | *Aldh1a1* | ENSRNOG00000017619 | aldehyde dehydrogenase 1A1, ventral SNc and ventral VTA subpopulation | McCaffery & Dräger 1994[7] | Poulin et al. 2014[8] | 95.6 |
| serotonergic | *Tph2* | ENSRNOG00000003880 | tryptophan hydroxylase 2, DR, MnR, B1 to B9 | Walther et al. 2003[9] | Okaty et al. 2019[10] | 476.4 |
| serotonergic | *Slc6a4* | ENSRNOG00000003476 | serotonin transporter, DR, MnR, B1 to B9 | Blakely et al. 1991[11] | Okaty et al. 2015[12] | 131.1 |
| serotonergic | *Fev* | ENSRNOG00000017856 | Pet1 transcription factor, DR, MnR, B1 to B9 | Hendricks et al. 2003[13] | Okaty et al. 2015[12] | 19.2 |
| shared monoaminergic | *Ddc* | ENSRNOG00000004327 | DOPA decarboxylase, all monoaminergic neurons | Berry et al. 1996[14] | Stansley & Yamamoto 2013[15] | 50.2 |
| shared monoaminergic | *Slc18a2* | ENSRNOG00000008890 | VMAT2, all monoaminergic neurons | Erickson et al. 1992[16] | Peter et al. 1995[17] | 95.3 |
| shared monoaminergic | *Gch1* | ENSRNOG00000011039 | GTP cyclohydrolase 1, monoamine synthesis | Ichinose et al. 1994[18] | Mencacci et al. 2014[19] | 49.7 |
| shared monoaminergic | *Slc17a8* | ENSRNOG00000007581 | VGLUT3, subpopulation of raphe serotonergic neurons, also striatal cholinergic interneurons, hippocampus and cortex | Gras et al. 2002[20] | - | 7.8 |
| oligodendrocyte | *Mbp* | ENSRNOG00000016516 | myelin basic protein | Roach et al. 1983[21] | Marques et al. 2016[22] | 38,001.4 |
| oligodendrocyte | *Plp1* | ENSRNOG00000002419 | proteolipid protein 1 | Milner et al. 1985[23] | Marques et al. 2016[22] | 10,973.0 |
| oligodendrocyte | *Mobp* | ENSRNOG00000018700 | myelin-associated oligodendrocyte basic protein | Yamamoto et al. 1994[24] | Marques et al. 2016[22] | 10,024.9 |
| oligodendrocyte | *Cnp* | ENSRNOG00000017496 | 2',3'-cyclic-nucleotide 3'-phosphodiesterase | Sprinkle 1989[25] | Marques et al. 2016[22] | 2,433.4 |
| oligodendrocyte | *Mog* | ENSRNOG00000078049 | myelin oligodendrocyte glycoprotein | Gardinier et al. 1992[26] | Marques et al. 2016[22] | 354.8 |
| oligodendrocyte | *Mag* | ENSRNOG00000021023 | myelin-associated glycoprotein | Quarles et al. 1973[27] | Marques et al. 2016[22] | 577.1 |
| oligodendrocyte | *Sox10* | ENSRNOG00000011305 | SRY-box transcription factor 10 | Stolt et al. 2002[28] | Marques et al. 2016[22] | 95.6 |
| oligodendrocyte | *Cmtm5* | ENSRNOG00000016828 | CKLF-like MARVEL transmembrane domain-containing 5 | Buscham et al. 2022[29] | Marques et al. 2016[22] | 594.0 |
| oligodendrocyte | *Mal* | ENSRNOG00000015445 | myelin and lymphocyte protein | Schaeren-Wiemers et al. 1995[30] | Marques et al. 2016[22] | 826.2 |
| oligodendrocyte | *Cldn11* | ENSRNOG00000010263 | claudin-11 (oligodendrocyte-specific protein), myelin protein | Bronstein et al. 1996[31] | - | 278.3 |
| oligodendrocyte | *Gal3st1* | ENSRNOG00000042041 | cerebroside sulfotransferase, sulfatide synthesis | Honke et al. 2002[32] | - | 121.0 |

Markers chosen a priori from the literature. Mean TPM, mean of the three tVTA libraries. A8 to A14, dopaminergic cell groups; B1 to B9, serotonergic cell groups; DR, dorsal raphe nucleus; MnR, median raphe nucleus; SNc, substantia nigra pars compacta; TPM, transcripts per million; VTA, ventral tegmental area; -, none.

**Supplementary Table 2 | Sensitivity of the twenty most enriched genes to the ranking criterion.**

| Gene | Ensembl gene ID | Rank, fold change | Rank, adjusted P | Shrunken log2 fold change | Adjusted P | Mean TPM, tVTA | Mean TPM, whole brain | Contamination family | In both |
|---|---|---|---|---|---|---|---|---|---|
| *Slc6a4* | ENSRNOG00000003476 | 1 | 20 | 5.56 | 4.9e-133 | 131.1 | 3.7 | serotonergic | yes |
| *Tph2* | ENSRNOG00000003880 | 2 | 135 | 5.33 | 1.1e-51 | 476.4 | 15.5 | serotonergic | no |
| *Sox14* | ENSRNOG00000022084 | 3 | 5 | 5.24 | 1.7e-220 | 34.9 | 1.3 | - | yes |
| *Fev* | ENSRNOG00000017856 | 4 | 21 | 5.19 | 1.4e-129 | 19.2 | 0.7 | serotonergic | no |
| *Pax7* | ENSRNOG00000018739 | 5 | 687 | 4.79 | 2.2e-13 | 5.3 | 0.2 | - | no |
| *Cryba2* | ENSRNOG00000017996 | 6 | 470 | 4.59 | 1.0e-22 | 9.4 | 0.5 | - | no |
| *Otp* | ENSRNOG00000010560 | 7 | 143 | 4.53 | 2.1e-50 | 12.0 | 0.7 | - | no |
| *Gata3* | ENSRNOG00000019336 | 8 | 17 | 4.33 | 1.7e-138 | 17.8 | 1.2 | - | yes |
| *En1* | ENSRNOG00000056580 | 9 | 13 | 4.27 | 1.1e-151 | 18.7 | 1.3 | - | yes |
| *Pnoc* | ENSRNOG00000014231 | 10 | 1 | 4.19 | <1e-300 | 281.0 | 21.0 | - | yes |
| *Gchfr* | ENSRNOG00000012290 | 11 | 2 | 4.15 | 3.8e-259 | 213.9 | 16.4 | - | yes |
| *Irx6* | ENSRNOG00000011037 | 12 | 256 | 3.87 | 5.6e-36 | 6.8 | 0.6 | - | no |
| *Glp1r* | ENSRNOG00000001152 | 13 | 405 | 3.74 | 5.2e-26 | 6.1 | 0.6 | - | no |
| *Ddc* | ENSRNOG00000004327 | 14 | 44 | 3.56 | 9.2e-94 | 50.2 | 5.7 | shared monoaminergic | no |
| *Slc18a2* | ENSRNOG00000008890 | 15 | 24 | 3.53 | 3.1e-124 | 95.3 | 11.2 | shared monoaminergic | no |
| *Cbln4* | ENSRNOG00000004372 | 16 | 3 | 3.44 | 8.7e-240 | 60.5 | 7.6 | - | yes |
| *Slc6a3* | ENSRNOG00000017302 | 17 | 309 | 3.36 | 1.6e-32 | 32.7 | 4.2 | dopaminergic | no |
| *Irx3* | ENSRNOG00000011533 | 18 | 49 | 3.33 | 1.0e-90 | 20.1 | 2.7 | - | no |
| *Slc17a8* | ENSRNOG00000007581 | 19 | 271 | 3.30 | 1.2e-34 | 7.8 | 1.1 | shared monoaminergic | no |
| *Irx5* | ENSRNOG00000011180 | 20 | 101 | 3.30 | 3.6e-61 | 9.4 | 1.3 | - | no |
| *Gal3st1* | ENSRNOG00000042041 | 36 | 18 | 2.53 | 1.0e-135 | 121.0 | 28.5 | oligodendrocyte | no |
| *Vsig10l* | ENSRNOG00000023411 | 37 | 11 | 2.48 | 3.3e-156 | 57.5 | 14.0 | - | no |
| *Ppp1r14a* | ENSRNOG00000020676 | 58 | 8 | 2.25 | 1.2e-167 | 144.6 | 41.4 | - | no |
| *Cldn11* | ENSRNOG00000010263 | 90 | 6 | 2.00 | 2.1e-208 | 278.3 | 94.9 | oligodendrocyte | no |
| *Nap1l5* | ENSRNOG00000083613 | 97 | 7 | 1.96 | 3.2e-179 | 295.8 | 103.1 | - | no |
| *Scg2* | ENSRNOG00000015055 | 101 | 4 | 1.94 | 2.4e-224 | 598.3 | 212.2 | - | no |
| *Paqr6* | ENSRNOG00000026059 | 103 | 12 | 1.94 | 1.1e-151 | 203.8 | 72.5 | - | no |
| *Prmt2* | ENSRNOG00000001297 | 109 | 9 | 1.89 | 2.4e-167 | 197.8 | 72.5 | - | no |
| *Rnd2* | ENSRNOG00000020698 | 110 | 19 | 1.89 | 2.0e-134 | 547.5 | 201.1 | - | no |
| *Plekhb1* | ENSRNOG00000018627 | 154 | 15 | 1.70 | 5.9e-142 | 1,415.9 | 593.4 | - | no |
| *Cmtm5* | ENSRNOG00000016828 | 162 | 14 | 1.67 | 2.8e-144 | 594.0 | 254.7 | oligodendrocyte | no |
| *Gaa* | ENSRNOG00000047656 | 166 | 10 | 1.65 | 1.7e-157 | 326.2 | 141.4 | - | no |
| *Sec11c* | ENSRNOG00000017036 | 188 | 16 | 1.59 | 2.9e-140 | 1,314.4 | 594.9 | - | no |

Genes among the first twenty by shrunken log2 fold change or by adjusted P, ranked within the 863 eligible genes of Fig. 1c. In both, gene among the first twenty by fold change and by adjusted P. TPM, transcripts per million; -, none.

**Supplementary Table 3 | RNAscope probes.**

| Target | Species | Channel | Catalog no. (ACD) |
|---|---|---|---|
| *Sox14* | mouse | C1 | 516411 |
| *Sox14* | rat | C1 | 549571 |
| *Gad1* | mouse | C2 | 400951-C2 |
| *Gad1* | rat | C2 | 316401-C2 |
| *Gad2* | mouse | C3 | 439371-C3 |
| *Oprm1* | mouse | C3 | 315841-C3 |
| *Oprm1* | rat | C3 | 410691-C3 |
| *Th* | mouse | C2 | 317621-C2 |
| *Th* | rat | C2 | 314651-C2 |
| *Sstr1* | mouse | C2 | 437711-C2 |
| *Pnoc* | mouse | C3 | 437881-C3 |
| *Foxp1* | mouse | C2 | 586761-C2 |
| Positive control (*Polr2a*, *Ppib*, *Ubc*) | mouse | C1, C2, C3 | 320881 |
| Positive control (*Polr2a*, *Ppib*, *Ubc*) | rat | C1, C2, C3 | 320891 |
| Negative control (*DapB*) | any | C1, C2, C3 | 320871 |

ACD, Advanced Cell Diagnostics; C1 to C3, probe channels.

**Supplementary Table 4 | Antibodies.**

| Antibody | Type | Host | Dilution | Supplier | Catalog no. | RRID |
|---|---|---|---|---|---|---|
| Anti-NeuN (Fox3/Rbfox3) | primary | rabbit | 1:1,000 | Abcam | ab104225 | AB_10711153 |
| Anti-FOXP1 | primary | rabbit | 1:500 | Sigma-Aldrich / Millipore | AB2277 | AB_10631732 |
| Anti-TH | primary | chicken | 1:1,000 | Abcam | ab76442 | AB_1524535 |
| Anti-TH | primary | mouse | 1:1,000 | Santa Cruz Biotechnology | sc-25269 | AB_628422 |
| Anti-GFP | primary | rat | 1:1,000 | Nacalai Tesque | 04404-84 (clone GF090R) | AB_10013361 |
| Anti-V5 tag | primary | chicken | 1:1,000 | Novus Biologicals | NB600-379 | AB_10003214 |
| Donkey anti-rabbit, Alexa Fluor 488 | secondary | donkey | 1:400 | Invitrogen | A-21206 | AB_2535792 |
| Donkey anti-rabbit, Alexa Fluor 647 | secondary | donkey | 1:400 | Invitrogen | A-31573 | AB_2536183 |
| Goat anti-chicken, STAR635P | secondary | goat | 1:500 | Abberior | 2-0102-007-4 | - |
| Goat anti-chicken IgY, Alexa Fluor Plus 647 | secondary | goat | 1:400 | Invitrogen | A32933 | AB_2762845 |
| Goat anti-mouse IgG, Alexa Fluor 594 | secondary | goat | 1:400 | Invitrogen | A-11005 | AB_2534073 |
| Donkey anti-rat, Alexa Fluor 488 | secondary | donkey | 1:400 | Invitrogen | A-21208 | AB_2535794 |
| Streptavidin, Alexa Fluor 488 | conjugate | - | 1:1,000 | Invitrogen | S11223 | AB_2336881 |

RRID, Research Resource Identifier. TH, tyrosine hydroxylase. Dashes, not applicable, the streptavidin conjugate having no host species and the Abberior antibody no registered RRID.

**Supplementary Table 5 | The 100 custom genes added to the Xenium panel.**

| Gene | Ensembl gene ID | Gene | Ensembl gene ID |
|---|---|---|---|
| Adam11 | ENSMUSG00000020926 | Kcnip3 | ENSMUSG00000079056 |
| Adamtsl3 | ENSMUSG00000070469 | Kcnk13 | ENSMUSG00000045404 |
| Adcy5 | ENSMUSG00000022840 | Klhl1 | ENSMUSG00000022076 |
| Adcy7 | ENSMUSG00000031659 | Lgi2 | ENSMUSG00000039252 |
| Adgra1 | ENSMUSG00000025475 | Lrp1b | ENSMUSG00000049252 |
| Adgrb3 | ENSMUSG00000033569 | Mcc | ENSMUSG00000071856 |
| Art4 | ENSMUSG00000030217 | Nlgn1 | ENSMUSG00000063887 |
| Atp6ap1l | ENSMUSG00000078958 | Nmbr | ENSMUSG00000019865 |
| Avpr1a | ENSMUSG00000020123 | Npffr1 | ENSMUSG00000020090 |
| Bcan | ENSMUSG00000004892 | Nrg3 | ENSMUSG00000041014 |
| Cacna1e | ENSMUSG00000004110 | Nrgn | ENSMUSG00000053310 |
| Cacna1f | ENSMUSG00000031142 | Nrxn1 | ENSMUSG00000024109 |
| Cck | ENSMUSG00000032532 | Ntm | ENSMUSG00000059974 |
| Cckar | ENSMUSG00000029193 | Nxph1 | ENSMUSG00000046178 |
| Cdh15 | ENSMUSG00000031962 | Otud7a | ENSMUSG00000033510 |
| Cnr1 | ENSMUSG00000044288 | P2rx2 | ENSMUSG00000029503 |
| Cpa1 | ENSMUSG00000054446 | Paqr6 | ENSMUSG00000041423 |
| Cplx1 | ENSMUSG00000033615 | Pclo | ENSMUSG00000061601 |
| Crtac1 | ENSMUSG00000042401 | Penk | ENSMUSG00000045573 |
| Dpp6 | ENSMUSG00000061576 | Pld6 | ENSMUSG00000043648 |
| Ebf1 | ENSMUSG00000057098 | Ptp4a3 | ENSMUSG00000059895 |
| Elfn1 | ENSMUSG00000048988 | Ptprn | ENSMUSG00000026204 |
| Fam155a | ENSMUSG00000079157 | Ptprt | ENSMUSG00000053141 |
| Fat3 | ENSMUSG00000074505 | Rtn4rl1 | ENSMUSG00000045287 |
| Fev | ENSMUSG00000055197 | Rtn4rl2 | ENSMUSG00000050896 |
| Fgf13 | ENSMUSG00000031137 | Scg2 | ENSMUSG00000050711 |
| Flywch2 | ENSMUSG00000023911 | Sema4f | ENSMUSG00000000627 |
| Fnbp1l | ENSMUSG00000039735 | Sgk2 | ENSMUSG00000017868 |
| Gabrb1 | ENSMUSG00000029212 | Shank1 | ENSMUSG00000038738 |
| Gabre | ENSMUSG00000031340 | Shank2 | ENSMUSG00000037541 |
| Galntl6 | ENSMUSG00000096914 | Shisa9 | ENSMUSG00000022494 |
| Glrb | ENSMUSG00000028020 | Slc34a2 | ENSMUSG00000029188 |
| Gng8 | ENSMUSG00000063594 | Snhg11 | ENSMUSG00000044349 |
| Gpc5 | ENSMUSG00000022112 | Spp1 | ENSMUSG00000029304 |
| Gpr165 | ENSMUSG00000031210 | Srd5a1 | ENSMUSG00000021594 |
| Gprc5c | ENSMUSG00000051043 | Sstr5 | ENSMUSG00000050824 |
| Gprin3 | ENSMUSG00000045441 | Stk32b | ENSMUSG00000029123 |
| Gpx3 | ENSMUSG00000018339 | Stxbp2 | ENSMUSG00000004626 |
| Grin3a | ENSMUSG00000039579 | Syndig1 | ENSMUSG00000074736 |
| Grm5 | ENSMUSG00000049583 | Synpr | ENSMUSG00000056296 |
| Hepacam2 | ENSMUSG00000044156 | Syt1 | ENSMUSG00000035864 |
| Igfbp5 | ENSMUSG00000026185 | Syt17 | ENSMUSG00000058420 |
| Il16 | ENSMUSG00000001741 | Syt7 | ENSMUSG00000024743 |
| Irx1 | ENSMUSG00000060969 | Tmem132c | ENSMUSG00000034324 |
| Itpka | ENSMUSG00000027296 | Tmod1 | ENSMUSG00000028328 |
| Kcna1 | ENSMUSG00000047976 | Trim30a | ENSMUSG00000030921 |
| Kcnab1 | ENSMUSG00000027827 | Trpm3 | ENSMUSG00000052387 |
| Kcnc3 | ENSMUSG00000062785 | Vsig10l | ENSMUSG00000070604 |
| Kcnd1 | ENSMUSG00000009731 | Zar1 | ENSMUSG00000063935 |
| Kcnd3 | ENSMUSG00000040896 | Zfhx4 | ENSMUSG00000025255 |

Add-on panel mBrain_100g (design M3Z4TG) of the Xenium Mouse 5K Pan Tissue & Pathways Panel (5,006 genes); genes in alphabetical order, read down the first pair of columns, then the second.

**Supplementary Table 6 | Robustness of the delineation of the tVTA territory.**

| Family of variants | Route A (rat bulk signature) | Route B (GABAergic subgroup) | Route C (Allen clusters) | Final territory |
|---|---|---|---|---|
| Without one section | 5, 0.907 to 0.944 | 5, 0.929 to 0.965 | 5, 0.931 to 0.963 | one section per mouse: 4, 0.918 to 0.947 |
| Without one mouse | not run | not run | not run | 3, 0.896 to 0.949 |
| One mouse alone | 3, 0.321 to 0.793 | 3, 0.578 to 0.868 | 3, 0.569 to 0.854 | not run |
| One section alone | not run | 5, 0.285 to 0.688 | not run | not run |
| One hemisphere alone | not run | not run | not run | 2, 0.912 to 0.924 |
| Contour level | 3, 0.872 to 0.950 | 5, 0.823 to 0.943 | 4, 0.878 to 0.947 | 4, 0.891 to 0.954 |
| Consensus rule | - | - | - | 3, 0.852 to 0.931 |
| Smoothing (kernel width) | 4, 0.865 to 0.938 | 4, 0.934 to 0.974 | 4, 0.941 to 0.968 | not run |
| Minimum support of a node, minimum area | 5, 0.988 to 1.000 | 4, 1.000 | 4, 1.000 | support 5 to 60: 7, 0.921 to 0.999 |
| Window around the raphe | 3, 0.939 to 0.989 | 3, 0.985 to 0.997 | 3, 0.986 to 1.000 | - |
| Separation of the lobes | - | - | - | 2, 1.000 |
| Gene list and detection thresholds | 5, 0.841 to 0.988 | 1, 1.000 | - | - |
| Score weighting | 3, 0.782 to 0.980 | 1, 0.956 | - | - |
| Subgroup resolution and composition | - | resolution: 5, 0.000 to 0.988; ranks merged: 3, 0.000 to 0.941 | cluster added: 3, 0.971 to 0.985; cluster dropped: 2, 0.795 to 0.855 | - |
| Scored population | all neurons, not only GABAergic: 1, 0.000 | - | 4, 0.875 to 0.962 | - |
| Earlier settings, combined | 1, 0.909 | - | - | - |
| Negative controls, largest Dice (13 unrelated populations and cluster 4701) | 0.101 (cluster 3435) | not run | 0.016 (cluster 3435) | 0.043 (cluster 3435) |

Number of reruns per family of variants and range of their Dice coefficient with the reference territory of the same route, or with the final territory. Route C was read after the limits were drawn.

### Supplementary Table 7 | Cell composition of the tVTA territory.

**a | All cells by class**

| Cell class | Mouse 1 | Mouse 2 | Mouse 3 | All | Proportion in the tVTA (%) | Proportion on the whole sections (%) |
|---|---|---|---|---|---|---|
| Oligodendrocytes | 776 | 783 | 403 | 1,962 | 43.5 | 19.0 |
| GABAergic neurons | 321 | 339 | 176 | 836 | 18.5 | 14.3 |
| Astrocytes | 198 | 196 | 102 | 496 | 11.0 | 14.6 |
| Endothelial cells | 154 | 176 | 68 | 398 | 8.8 | 7.9 |
| Oligodendrocyte precursors | 82 | 93 | 52 | 227 | 5.0 | 2.7 |
| Microglia | 55 | 71 | 36 | 162 | 3.6 | 2.5 |
| Vascular smooth muscle cells | 66 | 36 | 19 | 121 | 2.7 | 2.5 |
| Mural cells | 42 | 54 | 23 | 119 | 2.6 | 2.0 |
| Serotonergic neurons | 11 | 38 | 11 | 60 | 1.3 | 0.5 |
| Dopaminergic neurons | 3 | 20 | 3 | 26 | 0.6 | 0.4 |
| Glutamatergic neurons | 16 | 21 | 9 | 46 | 1.0 | 27.8 |
| Fibroblasts (VLMC) | 22 | 9 | 7 | 38 | 0.8 | 4.6 |
| Perivascular macrophages | 13 | 5 | 5 | 23 | 0.5 | 1.0 |
| Ependymal cells | 0 | 0 | 0 | 0 | 0.0 | 0.2 |
| All cells | 1,759 | 1,841 | 914 | 4,514 | 100.0 | 100.0 |

**b | Neurons by compartment**

| Neuron class | n | Dorsal part | Medial ventral wing | Lateral ventral wing |
|---|---|---|---|---|
| GABAergic neurons | 836 | 333 (40%) | 252 (30%) | 251 (30%) |
| Serotonergic neurons | 60 | 39 (65%) | 9 (15%) | 12 (20%) |
| Dopaminergic neurons | 26 | 21 (81%) | 1 (4%) | 4 (15%) |
| Glutamatergic neurons | 46 | 9 (20%) | 6 (13%) | 31 (67%) |

**c | Sox14 by neuron class**

| Neuron class | Sox14+ in the tVTA | Sox14+ in the rest of the five sections |
|---|---|---|
| GABAergic neurons | 404 / 836 (48.3%) | 11,236 / 48,473 (23.2%) |
| Serotonergic neurons | 1 / 60 (1.7%) | 75 / 2,286 (3.3%) |
| Dopaminergic neurons | 0 / 26 (0.0%) | 14 / 899 (1.6%) |
| Glutamatergic neurons | 3 / 46 (6.5%) | 1,233 / 81,870 (1.5%) |

a, All cells of the tVTA territory by class and mouse; proportion of each class in the tVTA and on the whole Xenium sections (435,402 cells). b, Neurons of the tVTA by compartment, three mice pooled. c, Neurons expressing Sox14, by class, in the tVTA and in the rest of the five tVTA sections, three mice pooled.

### Supplementary Table 8 | Sox14 and serotonin receptors in the tVTA, its rings and its neighbors.

**a | The tVTA and its rings**

| Population | GABAergic neurons (n) | Sox14+, mean of the mice | Mouse 1 | Mouse 2 | Mouse 3 |
|---|---|---|---|---|---|
| tVTA | 836 | 48.0 | 48.0 | 49.6 | 46.6 |
| Ring 0 to 100 µm | 844 | 28.1 | 24.2 | 25.9 | 34.1 |
| Ring 100 to 200 µm | 998 | 14.3 | 16.8 | 12.1 | 14.1 |
| Ring 200 to 300 µm | 1,127 | 10.6 | 10.7 | 9.5 | 11.5 |
| Ring 300 to 400 µm | 1,507 | 8.1 | 8.3 | 7.0 | 9.0 |
| Ring 400 to 500 µm | 1,492 | 9.7 | 10.0 | 8.5 | 10.5 |
| tVTA-to-rings ratio | - | 3.41 (geometric mean) | 3.42 | 3.93 | 2.94 |
| 1,000 random placements of the territory, ratio | - | 95th percentile 1.31; 100% below the real ratio | - | - | - |

**b | The ten neighbors**

| Population | Distance to the tVTA edge (µm) | Neurons (n) | Sox14+ (%) | tVTA-to-neighbor ratio (geometric mean) |
|---|---|---|---|---|
| Dorsal raphe nucleus | 12 | 346 | 13.5 | 3.49 |
| Dorsal band | 12 | 1,903 | 20.0 | 2.36 |
| Median raphe nucleus | 94 | 1,809 | 2.6 | 13.82 |
| Otp territory | 129 | 1,190 | 3.4 | 12.03 |
| Pedunculopontine nucleus, GABAergic | 262 | 1,227 | 11.8 | 3.87 |
| Pedunculopontine nucleus, cholinergic | 394 | 1,427 | 10.5 | 4.35 |
| Periaqueductal gray | 468 | 3,396 | 15.9 | 2.95 |
| Ventral nucleus of the lateral lemniscus | 896 | 1,319 | 5.5 | 10.26 |
| Superior colliculus | 1,901 | 17,183 | 40.4 | 1.18 |
| Same molecular family, scattered | by label | 336 | 50.1 | 0.96 |

**c | Xenium against RNAscope**

| Measure | Xenium, mean of the mice (mice 1; 2; 3) | RNAscope, mean ± s.e.m. |
|---|---|---|
| Sox14+ among the Gad1+ GABAergic neurons of the tVTA (%) | 59.1 (62.9; 58.8; 55.6) | 70.3 ± 2.6 |
| Sox14 transcripts per Sox14+ GABAergic neuron of the tVTA | 1.99 (1.85; 2.14; 1.98) | - |

**d | Serotonin receptors of the panel**

| Gene | Detection threshold (transcripts) | GABAergic neurons of the tVTA (%), mean (mice 1; 2; 3) | Sox14+ neurons of the tVTA (%), mean (mice 1; 2; 3) | Rings 0 to 500 µm (%) | tVTA-to-rings ratio | Rank |
|---|---|---|---|---|---|---|
| Htr2c | 2 | 39.8 (38.0; 39.8; 41.5) | 47.5 (48.7; 51.2; 42.7) | 20.6 | 1.93 | 13 |
| Htr7 | 1 | 28.7 (26.5; 31.9; 27.8) | 32.7 (33.1; 34.5; 30.5) | 23.3 | 1.23 | 89 |
| Htr1a | 1 | 12.9 (14.3; 14.2; 10.2) | 16.1 (18.2; 16.7; 13.4) | 15.3 | 0.84 | - |
| Htr2a | 1 | 11.4 (11.8; 12.7; 9.7) | 13.0 (13.6; 13.1; 12.2) | 14.8 | 0.76 | - |
| Htr1f | 1 | 8.6 (7.5; 8.6; 9.7) | 10.5 (10.4; 11.3; 9.8) | 4.3 | 2.00 | - |
| Htr4 | 1 | 8.4 (8.1; 9.7; 7.4) | 8.9 (10.4; 10.1; 6.1) | 10.8 | 0.78 | - |
| Htr1b | 1 | 1.6 (1.2; 0.6; 2.8) | 1.6 (0.6; 0.6; 3.7) | 1.8 | 0.72 | - |
| Htr1d | 1 | 0.3 (0.6; 0.3; 0.0) | 0.0 (0.0; 0.0; 0.0) | 0.3 | - | - |
| Htr3a | 1 | 0.3 (0.6; 0.3; 0.0) | 0.2 (0.0; 0.6; 0.0) | 0.7 | - | - |
| Htr3b | 1 | 0.1 (0.0; 0.3; 0.0) | 0.0 (0.0; 0.0; 0.0) | 0.1 | - | - |
| Htr2b | 1 | 0.0 (0.0; 0.0; 0.0) | 0.0 (0.0; 0.0; 0.0) | 0.1 | - | - |

a, GABAergic neurons of the tVTA and of the five 100 µm rings around it; *Sox14*+ proportions in %, mean of the three mice and per mouse; ratios without unit. b, The ten neighbors, ordered by distance to the tVTA edge; tVTA-to-neighbor ratio, geometric mean of the per-mouse ratios, one percentage point added to both proportions. c, RNAscope value of the Results (mean ± s.e.m. across animals). d, The eleven serotonin receptor genes of the panel, in the GABAergic neurons of the tVTA and in their *Sox14*+ subset; ratio as in b; rank among the 1,259 genes detected in at least a fifth of those neurons; Htr5b and Htr6 are not in the panel.

### Supplementary Table 9 | The six Allen clusters of the tVTA and their local markers.

**a | The six clusters**

| Allen cluster | GABAergic neurons in the tVTA, of the five sections | Position in the tVTA | Sox14+ (%) | Pnoc+ (%) | MERFISH atlas, cells in the tVTA territory |
|---|---|---|---|---|---|
| 4687 | 86 of 143 (60%) | dorsal summit | 59 (50 of 85) | 21 (18 of 86) | 132 of 171 (77%) |
| 4703 | 96 of 118 (81%) | medial column | 82 (69 of 84) | 78 (75 of 96) | 178 of 190 (94%) |
| 4694 | 115 of 145 (79%) | medial ventral wing | 78 (88 of 113) | 37 (42 of 115) | 98 of 113 (87%) |
| 4698 | 89 of 168 (53%) | lateral ventral wing | 71 (56 of 79) | 33 (29 of 89) | 51 of 82 (62%) |
| 4697 | 8 of 32 (25%) | not described | 72 (13 of 18), 4697 and 4702 together | 39 (7 of 18), 4697 and 4702 together | 26 of 49 (53%) |
| 4702 | 10 of 26 (38%) | not described | 72 (13 of 18), 4697 and 4702 together | 39 (7 of 18), 4697 and 4702 together | 22 of 32 (69%) |
| The six together | 404 of 632 (64%); 404 of the 836 tVTA GABAergic neurons | - | - | - | 507 of 637 (80%); 7 sections, 1.14 mm, 0.78 mm3 |
| The 12 other clusters of subclass 268 | 22 of 358 (6%) | scattered | 60 (9 of 15) | 32 (7 of 22) | 44 of 352 (12.5%) |

**b | Local markers**

| Gene | Allen cluster | Both detection thresholds | Xenium, proportion in the cluster (%) | Xenium, ratio against the other three clusters | Allen 10x cells | Allen MERFISH atlas |
|---|---|---|---|---|---|---|
| Ikzf1 | 4687 | yes | 40.7 | 5.55 | detected, not selective (96%) | not in panel |
| Npffr2 | 4687 | yes | 32.4 | 16.76 | detected, not selective (24%) | not in panel |
| Pth2r | 4687 | yes | 29.7 | 2.92 | detected, not selective (37%) | selective (48%) |
| Rspo1 | 4687 | no | 27.6 | 5.94 | detected, not selective (58%) | selective (38%) |
| Rxfp3 | 4703 | no | 35.6 | 2.93 | detected, not selective (31%) | detected, not selective (34%) |
| Wnt5a | 4703 | yes | 32.5 | 2.23 | detected, not selective (82%) | not in panel |
| Sulf1 | 4703 | no | 25.8 | 2.30 | detected, not selective (70%) | not in panel |
| Ephb1 | 4694 | yes | 26.3 | 2.05 | detected, not selective (100%) | not in panel |
| Nmbr | 4694 | no | 25.0 | 2.33 | detected, not selective (20%) | not in panel |
| none | 4698 | - | - | - | - | - |
| Egflam | 4703 | no | not in the Xenium panel | - | selective (81%) | selective (74%) |

a, Clusters of Allen subclass 268 (CS-PRNr-DR En1 Sox2 Gaba) with at least a quarter of their GABAergic neurons of the five sections inside the tVTA, three mice pooled; Sox14 read with the Allen labels obtained without Sox14; MERFISH atlas, cells of the cluster on the atlas sections of the tVTA. b, Rows of Fig. 4f; proportion and ratio on our sections (main detection line); Allen 10x cells and MERFISH atlas, percentages as in Fig. 4f.

**Supplementary Table 10 | Properties of the sixteen VTA neurons in which the light-evoked IPSC was analyzed.**

| Neuron | Latency (ms) | Jitter (ms) | Rise (ms) | Amplitude (pA) | Decay τw (ms) | Cm (pF) | Rm (MΩ) | First spike delay (ms) | Action potential duration (ms) | TH | Location |
|---|---|---|---|---|---|---|---|---|---|---|---|
| P1 | 2.08 | 0.044 | 1.20 | 258 | 7.80 | 44 | 222 | 169 | 2.50 | + | VTA or SNc |
| P2 | 2.05 | 0.038 | 1.00 | 152 | 12.70 | 35 | 373 | 124 | 2.30 | n.d. | VTA or SNc |
| P3 | 2.17 | 0.033 | 1.00 | 255 | 8.20 | 35 | 365 | 124 | 2.70 | n.d. | VTA or SNc |
| P4 | 2.30 | 0.080 | 1.70 | 706 | 10.23 | 39 | 476 | 815 | 3.80 | + | VTA |
| P5 | 2.27 | 0.057 | 1.07 | 268 | 6.50 | 71 | 154 | 444 | 2.55 | n.d. | VTA |
| P6 | 2.70 | 0.190 | 1.40 | 93 | 9.90 | 29 | 445 | 1120 | 4.00 | - | VTA |
| P7 | 2.00 | 0.060 | 0.80 | 283 | 7.00 | 43 | 767 | 999 | 4.00 | n.d. | n.d. |
| P8 | 1.70 | 0.070 | 1.50 | 111 | 15.70 | 38 | 546 | 318 | 3.16 | n.d. | n.d. |
| P9 | 2.00 | 0.050 | 1.40 | 327 | 12.40 | 43 | 236 | 1160 | 4.32 | + | VTA |
| P10 | 2.00 | 0.046 | 0.80 | 477 | 7.00 | 43 | 268 | 1027 | 4.18 | + | VTA |
| P11 | 2.40 | 0.120 | 1.20 | 157 | 10.00 | 25 | 193 | 630 | 2.70 | - | VTA |
| P12 | 2.11 | 0.050 | 1.12 | 188 | 14.43 | 62 | 155 | n.d. | n.d. | - | VTA (uncertain) |
| P13 | 2.20 | 0.120 | 1.20 | 52 | 16.50 | 32 | 82 | n.d. | 6.25 | + | VTA |
| P14 | 1.96 | 0.380 | 1.75 | 278 | n.d. | 46 | 309 | n.d. | n.d. | + | VTA |
| P15 | 3.20 | 0.070 | 1.00 | 348 | 9.65 | 49 | 320 | 225 | 3.14 | + | VTA |
| P16 | 3.51 | 0.110 | 0.84 | 174 | 4.80 | 39 | 250 | 885 | 3.00 | + | VTA |
| mean ± s.e.m. (n) | 2.29 ± 0.12 (16) | 0.095 ± 0.022 (16) | 1.19 ± 0.07 (16) | 258 ± 40 (16) | 10.19 ± 0.91 (15) | 42 ± 3 (16) | 323 ± 43 (16) | 618 ± 111 (13) | 3.47 ± 0.28 (14) | 8 + / 3 - / 5 n.d. | |

Jitter, s.d. of the latency across the sweeps of that neuron; decay, weighted time constant; TH, tyrosine hydroxylase immunodetection; location, post hoc site of the recording; n.d., not determined. Sixteen of the twenty-two responsive neurons; six were excluded from the analysis of the IPSC kinetics because the recording was unstable, the IPSC too small or composite.

**Supplementary Table 11a | Statistical analyses of the ex vivo electrophysiology.**

| Figure panel | Measure | Groups and n | Test and tails | Statistic and exact P | Effect size [95% CI] | Post-hoc |
|---|---|---|---|---|---|---|
| Fig. 5e | First IPSC amplitude, control vs DAMGO (1 µM) | n = 6 neurons<br>unit: neuron | Wilcoxon signed rank, paired, two-tailed, exact | W = 0, P = 0.03125<br>6 non-zero pairs of 6, smallest attainable P = 0.03125, reached | First IPSC from 312 ± 95 to 89 ± 35 pA, 26 ± 7% of control; reduction mean 223 pA, s.e.m. 75, 95% CI 29 to 417 pA<br>Reduced in 6 of 6 neurons; rank-biserial r = 1.00 | - |
| Fig. 5e | Paired pulse ratio, control vs DAMGO (1 µM) | n = 6 neurons<br>unit: neuron | Wilcoxon signed rank, paired, two-tailed, exact | W = 0, P = 0.03125<br>6 non-zero pairs of 6, smallest attainable P = 0.03125, reached | PPR from 0.83 ± 0.03 to 1.15 ± 0.11; increase mean 0.32, s.e.m. 0.09, 95% CI 0.09 to 0.54<br>Increased in 6 of 6 neurons; rank-biserial r = 1.00 | - |
| Fig. 5f | Interspike interval, ISI1 vs ISI2, single 1 ms flash | n = 8 neurons<br>unit: neuron, each value the mean of at least 20 episodes | Wilcoxon signed rank, paired, two-tailed, exact | W = 0, P = 0.0078125<br>8 non-zero pairs of 8, smallest attainable P = 0.0078125, reached | ISI2 minus ISI1: mean 177 ms, s.e.m. 50, 95% CI 60 to 294 ms<br>Lengthened in 8 of 8 neurons; rank-biserial r = 1.00 | - |
| Fig. 5g | Interspike interval, ISI1 vs ISI2, ten 1 ms flashes at 20 Hz | n = 8 neurons<br>unit: neuron, each value the mean of at least 20 episodes | Wilcoxon signed rank, paired, two-tailed, exact | W = 0, P = 0.0078125<br>8 non-zero pairs of 8, smallest attainable P = 0.0078125, reached | ISI2 minus ISI1: mean 636 ms, s.e.m. 97, 95% CI 406 to 865 ms<br>Lengthened in 8 of 8 neurons; rank-biserial r = 1.00 | - |

One row per test, unit named in the third column; a dash, not applicable. Alpha 0.05; every test paired within neurons and two-tailed; exact Wilcoxon signed-rank test computed with SciPy 1.18.1 under Python 3.13; values are mean ± s.e.m.; no correction for multiple comparisons was applied.

### Supplementary Table 11b | Statistical analyses of the in vivo electrophysiology and fiber photometry.

**b | Figure 6 and Supplementary Figure 11**

| Figure panel | Measure | Groups and n | Test and tails | Statistic and exact P | Effect size [95% CI] | Post-hoc |
|---|---|---|---|---|---|---|
| Fig. 6c | Photoidentification: response above the chance level set by each neuron's own spontaneous rate | n = 9 neurons, 5 mice<br>unit: pulse<br>80 trains, 800 pulses | Exact binomial, one-tailed | 28/A P = 1.48e-42; 30/C P = 3.79e-130; 30/D P = 1.45e-150; 31/C P < 1e-300; 31/D P = 1.60e-284; 31/H P < 1e-300; 36/A P = 3.20e-301; 36/C P = 5.77e-183; 37/A P < 1e-300 | Fidelity, median 0.955, range 0.771 to 0.990 | - |
| Fig. 6d | Firing rate, before vs light, tVTA neurons | n = 9 neurons, 5 mice<br>unit: train | Wilcoxon signed rank, paired, two-tailed | 28/A P = 0.25, n.t., three trains, smallest attainable P = 0.25<br>30/C P = 1.95e-03; 30/D P = 1.95e-03; 31/C P = 9.77e-04; 31/D P = 1.56e-02; 31/H P = 4.88e-04; 36/A P = 7.81e-03; 36/C P = 1.56e-02; 37/A P = 4.88e-04 | Spikes per light pulse 0.77 to 2.91<br>Rank-biserial r = 1.00 in the nine neurons<br>Per-neuron rates in Supplementary Table 13 | - |
| Fig. 6d | Firing rate, before vs after the train, tVTA neurons | n = 9 neurons, 5 mice<br>unit: train | Wilcoxon signed rank, paired, two-tailed | 28/A P = 0.750, n.t., 3 non-zero pairs, smallest attainable P = 0.25<br>30/C P = 7.81e-03; 30/D P = 7.81e-03, 8 non-zero pairs of 10 each<br>The six neurons silent before the light have no non-zero pair and are not testable | 16.40 to 9.40 Hz and 9.20 to 5.40 Hz<br>The two spontaneously active neurons return below their own baseline | - |
| No panel (Results) | Fidelity of the light response, 30 vs 50 Hz | n = 3 neurons, 1 mouse<br>unit: train | Mann-Whitney U, two-tailed | 31/C P = 0.059; 31/D P = 1.000; 31/H P = 1.000<br>Smallest attainable two-sided P, 2 / C(n30 + n50, n50): 1.6e-04, 1.2e-03 and 2.2e-02<br>The three tests could each have reached significance; two of them compare values that are all tied at 100 % | Fidelity 100.0 vs 96.7, 100.0 vs 100.0, 100.0 vs 100.0 %<br>Rank-biserial r = -0.33, 0.00, 0.00<br>138 of the 140 pulses delivered at 50 Hz are answered, and 42 of the 44 trains of this analysis answer every pulse | - |
| Supplementary Fig. 11a | Trend of fidelity along the 957 ms trains of thirty pulses, planned linear contrast of the six blocks of five pulses, one contrast per train | n = 3 neurons, 2 mice<br>unit: train<br>1109 pulses | Wilcoxon signed rank on the planned linear contrast, two-tailed | 30/C P = 0.0031, 11 non-zero contrasts of 20, smallest attainable P = 0.0010<br>30/D P = 1.000, n.t., 1 non-zero contrast of 10, smallest attainable P = 1.000<br>31/H P = 0.0156, 7 non-zero contrasts of 7, smallest attainable P = 0.0156<br>Block by block against the first block, the non-zero pairs of 31/H number 1 to 4, so the smallest attainable P of any post-hoc is 0.125 | Median contrast +0.6, 0.0 and -1.0, so 30/C rises and 31/H falls<br>Fidelity by block of five pulses: 91 to 100 %, 98 to 100 %, 97 to 74 % | None possible |
| Fig. 6f | Firing rate, before vs light, VTA putative DA neurons | n = 3 neurons, 1 mouse<br>unit: train | Wilcoxon signed rank, paired, two-tailed | 30/C P = 1.953e-03; 30/E P = 5.79e-05; 30/F P = 7.83e-09 | Inhibition -93.5, -100 and -94.9 % of the before rate<br>Rank-biserial r = -1.00, -1.00 and -0.99 | - |
| Fig. 6f | Firing rate, light vs after, VTA putative DA neurons | n = 3 neurons, 1 mouse<br>unit: train | Wilcoxon signed rank, paired, two-tailed | 30/C P = 1.953e-03; 30/E P = 3.81e-05; 30/F P = 7.20e-10 | After rate 11.6, 2.2 and 12.8 Hz against 0.60, 0.00 and 0.42 Hz under the light<br>Rank-biserial r = 1.00 in the three neurons | - |
| Fig. 6f | Firing rate, before vs after the train, VTA putative DA neurons | n = 3 neurons, 1 mouse<br>unit: train | Wilcoxon signed rank, paired, two-tailed | 30/C P = 9.38e-02, n.s., 9 non-zero pairs of 10, smallest attainable P = 3.91e-03<br>30/E P = 1.29e-02, 10 non-zero pairs of 20<br>30/F P = 4.88e-07, 45 non-zero pairs of 50 | Before 9.20, 3.20 and 8.28 Hz against after 11.60, 2.20 and 12.80 Hz<br>Rank-biserial r = 0.67, -0.84 and 0.86<br>Firing does not return to the same level in all three: it ends above baseline in 30/F, below it in 30/E, and does not differ in 30/C | - |
| Fig. 6g | Accumbal dLight signal, mean z over 0 to +1 s vs the matched window -1 to 0 s, 30 Hz | n = 4 mice, 10 trials each<br>unit: trial | Wilcoxon signed rank, paired, two-tailed<br>No group test: with four animals the smallest attainable P is 0.125 | W = 0, 0, 4, 0<br>P = 1.953e-03, 1.953e-03, 1.367e-02, 1.953e-03 | Mean difference in z: -3.32, -2.05, -1.32, -5.33<br>Rank-biserial \|r\| = 1.00, 1.00, 0.86 and 1.00, all falls | - |
| Fig. 6h | Effect of stimulation frequency on End (ms), over 10, 30 and 50 Hz | n = 4 mice<br>unit: animal | One-way repeated-measures ANOVA | F(2,6) = 54.018, uncorrected P = 1.457e-04<br>epsilon = 0.597, corrected P = 2.564e-03 | Partial eta squared = 0.947, 95% CI 0.665 to 0.968 | none |
| Fig. 6h | Effect of stimulation frequency on Trough after light offset (ms), over 10, 30 and 50 Hz; descriptive measure, defined after inspection of the traces | n = 4 mice<br>unit: animal | One-way repeated-measures ANOVA | F(2,6) = 4.880, uncorrected P = 5.518e-02<br>epsilon = 0.524, corrected P = 1.102e-01 | Partial eta squared = 0.619, 95% CI 0.000 to 0.775 | none |
| Fig. 6h | Effect of stimulation frequency on Fraction of trough at light offset, over 10, 30 and 50 Hz; descriptive measure, defined after inspection of the traces | n = 4 mice<br>unit: animal | One-way repeated-measures ANOVA | F(2,6) = 45.553, uncorrected P = 2.359e-04<br>epsilon = 0.621, corrected P = 2.916e-03 | Partial eta squared = 0.938, 95% CI 0.616 to 0.962 | none |
| Supplementary Fig. 11b | Effect of stimulation frequency on Trough (z), over 10, 30 and 50 Hz | n = 4 mice<br>unit: animal | One-way repeated-measures ANOVA | F(2,6) = 4.436, uncorrected P = 6.566e-02<br>epsilon = 0.513, corrected P = 1.236e-01 | Partial eta squared = 0.597, 95% CI 0.000 to 0.762 | none |
| Supplementary Fig. 11c | Response duration against train duration, within each animal | n = 4 mice, 4 or 5 frequencies each<br>unit: frequency | Least-squares linear regression, two-tailed | BM30 P = 2.002e-02; BM31 P = 3.072e-05; BM36 P = 7.293e-04; BM37 P = 9.334e-04 | Slopes 0.526, 0.868, 0.968, 0.602; r = 0.934, 0.999, 0.999, 0.992 | - |

| Figure panel | Measure | Groups and n | Test and tails | Statistic and exact P | Effect size [95% CI] | Post-hoc |
|---|---|---|---|---|---|---|
| | | | The nineteen points are not independent, so the fits are not combined | | 95% CI of r: 0.29 to 1.00, 0.98 to 1.00, 0.95 to 1.00, 0.88 to 1.00 Intercepts 683, 408, 205 and 727 ms, below the shortest train measured, 180 ms | |
| Supplementary Fig. 11c | Effect of stimulation frequency on Onset (ms), over 10, 30 and 50 Hz | n = 4 mice unit: animal | One-way repeated-measures ANOVA | F(2,6) = 0.345, uncorrected P = 7.215e-01 epsilon = 0.574, corrected P = 6.222e-01 | Partial eta squared = 0.103, 95% CI 0.000 to 0.398 | none |
| Supplementary Fig. 11c | Effect of stimulation frequency on Duration (ms), over 10, 30 and 50 Hz | n = 4 mice unit: animal | One-way repeated-measures ANOVA | F(2,6) = 39.726, uncorrected P = 3.462e-04 epsilon = 0.593, corrected P = 4.449e-03 | Partial eta squared = 0.930, 95% CI 0.573 to 0.957 | none |
| Supplementary Fig. 11d | Isosbestic control, 465 vs 405 nm over 0 to +1 s, both channels through identical processing | n = 4 mice, 10 trains each unit: train | Wilcoxon signed rank, paired, two-tailed | BM30 P = 1.953e-03; BM31 P = 1.953e-03; BM36 P = 9.766e-03; BM37 P = 1.953e-03 | Difference 465 minus 405 in z: -3.57, -2.26, -1.54, -5.20 Rank-biserial \|r\| = 1.00, 1.00, 0.89 and 1.00, the 465 nm channel below 405 nm in every animal | - |
| Methods | Return to baseline, each successive 2 s window against the 2 s preceding the train, five windows | n = 4 mice, 10 trains each unit: train | Wilcoxon signed rank, paired, two-tailed | 0 to 2 s: BM30 P = 1.953e-03 **; BM31 P = 2.324e-01; BM36 P = 1.055e-01; BM37 P = 1.953e-03 ** 2 to 4 s: BM30 P = 2.754e-01; BM31 P = 9.219e-01; BM36 P = 5.566e-01; BM37 P = 1.934e-01 4 to 6 s: BM30 P = 2.734e-02 *; BM31 P = 3.711e-02 *; BM36 P = 7.695e-01; BM37 P = 6.250e-01 6 to 8 s: BM30 P = 2.734e-02 *; BM31 P = 3.750e-01; BM36 P = 4.922e-01; BM37 P = 6.250e-01 8 to 10 s: BM30 P = 1.055e-01; BM31 P = 4.922e-01; BM36 P = 5.566e-01; BM37 P = 3.750e-01 | 0 to 2 s: Mean level in z: -1.97, -0.38, -0.51, -1.97 Rank-biserial \|r\| = 1.00, 0.45, 0.60, 1.00 2 to 4 s: Mean level in z: -0.25, +0.07, -0.07, +0.25 Rank-biserial \|r\| = 0.42, 0.05, 0.24, 0.49 4 to 6 s: Mean level in z: -0.63, -0.43, -0.04, -0.24 Rank-biserial \|r\| = 0.78, 0.75, 0.13, 0.20 6 to 8 s: Mean level in z: -0.67, -0.15, +0.20, +0.05 Rank-biserial \|r\| = 0.78, 0.35, 0.27, 0.20 8 to 10 s: Mean level in z: -0.49, -0.20, -0.11, +0.20 Rank-biserial \|r\| = 0.60, 0.27, 0.24, 0.35 | - |
| Methods | Independence of successive trains: mean ΔF/F of the 2 s preceding each train against the rank of the train | n = 4 mice, 10 trains each unit: train | Least-squares linear regression, two-tailed | BM30 P = 4.899e-01; BM31 P = 2.388e-01; BM36 P = 9.115e-01; BM37 P = 1.819e-01 | Slopes 1.28e-04, -9.75e-04, -1.13e-04, 3.97e-04 per train, two positive and two negative | - |

b, one row per test; a dash, not applicable. Alpha 0.05; Wilcoxon and Mann-Whitney tests two-tailed; repeated-measures ANOVA on 10, 30 and 50 Hz, Greenhouse-Geisser corrected P as the value quoted; |r|, magnitude of the rank-biserial correlation only.

### Supplementary Table 11c | Statistical analyses of the connectome.

**c | Figure 8 and Supplementary Figure 13**

| Figure panel | Measure | Groups and n | Test and tails | Statistic and exact P | Effect size [95% CI] | Post-hoc |
|---|---|---|---|---|---|---|
| Fig. 8b | Asymmetry of the unidirectional regions, afferent-only against efferent-only | 54 unidirectional regions of 124<br>unit: region | Exact binomial against 0.5, two-tailed | 14 of 54 regions afferent-only<br>P = 5.35e-04 | Proportion 0.259, 95% CI 0.150 to 0.397 | - |
| Fig. 8e | Ipsilateral bias of the input distribution | 84 afferent regions<br>unit: region | Wilcoxon signed-rank against 0, two-tailed | W = 375<br>P = 3.21e-10 | Median +0.113, 95% CI +0.081 to +0.175 (20,000 bootstrap resamples); r = 0.69 | - |
| Fig. 8e | Ipsilateral bias of the output distribution | 110 efferent regions<br>unit: region | Wilcoxon signed-rank against 0, two-tailed | W = 11<br>P = 1.18e-19 | Median +0.536, 95% CI +0.432 to +0.645 (20,000 bootstrap resamples); r = 0.86 | - |
| Supplementary Fig. 13a | Inter-animal agreement of the afferent map, log10 ipsilateral density | n = 2 mice, 74 subcortical regions<br>unit: region<br>Regions with a non-zero density in both mice | Pearson correlation, two-tailed | r = 0.9627, exact P = 1.28e-42 | r = 0.9627, 95% CI 0.941 to 0.976 | - |

c, one row per test, unit named in the third column; a dash, not applicable. Alpha 0.05; two-tailed tests; the two Wilcoxon tests are one-sample tests of each distribution against zero, inputs and outputs never being compared to each other; regions are not independent units, so these are tests of central tendency.

### Supplementary Table 11d | Statistical analyses of the behavioral experiments.

**d | Figures 9 and 10, Supplementary Figures 17 and 18**

| Figure panel | Measure | Groups and n | Test and tails | Statistic and exact P | Effect size [95% CI] | Post-hoc |
|---|---|---|---|---|---|---|
| Fig. 9b | Time in stim chamber (s), RTPA 20 min session | Ctrl 6, LH-tVTA 4, tVTA 7; Sox14-Cre | Two-way RM ANOVA | Day x group F(2,14) = 14.41, P = 0.0004; Day F(1,14) = 0.22, P = 0.65; Group F(2,14) = 12.97, P = 0.0006 | 0.67 [0.25, 0.79] | Sidak, pretest vs test. LH-tVTA P = 0.012; tVTA P = 0.0029; Ctrl n.s. |
| Fig. 9c | Locomotor activity in stim chamber (% time), RTPA 20 min session | Ctrl 6, LH-tVTA 4, tVTA 7; Sox14-Cre | Two-way RM ANOVA | Day x group F(2,14) = 20.75, P < 0.0001; Day F(1,14) = 23.11, P = 0.0003; Group F(2,14) = 13.05, P = 0.0006 | 0.75 [0.38, 0.84] | Sidak, pretest vs test. tVTA P < 0.0001; others n.s. |
| Fig. 9e | Locomotor activity (% time), open-field procedure 1 | Ctrl 11, LH-tVTA 10, tVTA 7; Sox14-Cre and C57BL/6J | Two-way RM ANOVA | Time x group F(10,125) = 5.812, P < 0.0001; Time F(3.027,75.68) = 18.45, P < 0.0001; Group F(2,25) = 5.326, P = 0.012 | 0.32 [0.14, 0.39] | Sidak, at each minute. Ctrl vs LH-tVTA min 6 P = 0.0280; Ctrl vs tVTA min 3 and 4 P < 0.0001; LH-tVTA vs tVTA min 3 P = 0.0049, min 4 P = 0.0481; others n.s. |
| Fig. 9f | Locomotor activity (% baseline), open-field procedure 2 | Ctrl 11, LH-tVTA 10, tVTA 7; Sox14-Cre and C57BL/6J | Two-way RM ANOVA | Time x group F(6,75) = 11.36, P < 0.0001; Time F(2.694,67.36) = 21.43, P < 0.0001; Group F(2,25) = 20.93, P < 0.0001 | 0.48 [0.27, 0.57] | Sidak, at each stimulation. Ctrl vs tVTA 30 Hz (5 s on/15 s off) P = 0.0105, 30 Hz and 60 Hz continuous P < 0.0001; LH-tVTA vs tVTA 30 Hz (5 s on/15 s off) P = 0.0257, 30 Hz continuous P = 0.0032, 60 Hz continuous P = 0.0002; others n.s. |
| Fig. 9g | Freezing time (s), open-field procedure 2 | Ctrl 11, LH-tVTA 10, tVTA 7; Sox14-Cre and C57BL/6J | Two-way RM ANOVA | Day x group F(6,75) = 12.94, P < 0.0001; Day F(1.584,39.61) = 29.62, P < 0.0001; Group F(2,25) = 34.62, P < 0.0001 | 0.51 [0.31, 0.59] | Sidak, at each stimulation. Ctrl vs tVTA 30 Hz (5 s on/15 s off) P = 0.0139, 30 Hz continuous P = 0.0032, 60 Hz continuous P = 0.0038; LH-tVTA vs tVTA 30 Hz (5 s on/15 s off) P = 0.0071, 30 Hz continuous P = 0.0121, 60 Hz continuous P = 0.0041; others n.s. |
| Fig. 9i | Time in stim chamber (% of pretest), repeated RTPA 10 min sessions | Ctrl 11, LH-tVTA 10, tVTA 6; Sox14-Cre and C57BL/6J | Two-way RM ANOVA | StimDay x group F(6,72) = 16.55, P < 0.0001; StimDay F(2.705,64.92) = 34.21, P < 0.0001; Group F(2,24) = 20.48, P < 0.0001 | 0.58 [0.39, 0.66] | Sidak, at each StimDay. Ctrl vs LH-tVTA StimDay1 P = 0.0019, StimDay2 and 3 P < 0.0001; Ctrl vs tVTA StimDay2 P = 0.0096, StimDay3 P = 0.0005; LH-tVTA vs tVTA StimDay1 P = 0.0107; others n.s. |
| Fig. 9j | Locomotor activity in stim chamber (% time), repeated RTPA 10 min sessions | Ctrl 11, LH-tVTA 10, tVTA 6; Sox14-Cre and C57BL/6J | Two-way RM ANOVA | StimDay x group F(6,72) = 11.31, P < 0.0001; StimDay F(2.625,63.00) = 41.54, P < 0.0001; Group F(2,24) = 36.15, P < 0.0001 | 0.49 [0.27, 0.58] | Sidak, at each day. Ctrl vs tVTA and LH-tVTA vs tVTA StimDay1 to 3 P < 0.0001; others n.s. |
| Fig. 10b | Time in stim chamber (% pretest time), repeated RTPA 10 min sessions | tVTA 4, VTA/SNc 5; Sox14-Cre | Two-way RM ANOVA | StimDay x group F(5,35) = 0.49, P = 0.78; StimDay F(2.040,14.28) = 37.67, P < 0.0001; Group F(1,7) = 4.77, P = 0.065 | 0.07 [0.00, 0.14] | Dunnett vs pretest. StimDay1 P = 0.0025; StimDay3 P = 0.0003; StimDay4 P < 0.0001; test P = 0.0009; StimDay2 n.s. |
| Fig. 10c | Locomotor activity in stim chamber (% time), repeated RTPA 10 min sessions | tVTA 4, VTA/SNc 5; Sox14-Cre | Two-way RM ANOVA | StimDay x group F(5,35) = 0.99, P = 0.44; StimDay F(3.308,23.15) = 138.2, P < 0.0001; Group F(1,7) = 1.548, P = 0.25 | 0.12 [0.00, 0.23] | Dunnett vs pretest. StimDay1 to 4 P < 0.0001; test n.s. |
| Fig. 10e | Locomotor activity (% time), open-field procedure 1 | tVTA 4, VTA/SNc 5; Sox14-Cre | Two-way RM ANOVA | Day x group F(5,35) = 0.74, P = 0.60; Day F(2.348,16.44) = 68.16, P < 0.0001; Group F(1,7) = 0.03, P = 0.87 | 0.10 [0.00, 0.19] | Sidak, pairwise across minutes 1 to 6. 1 vs 3, 1 vs 4, 2 vs 3 and 2 vs 4 P < 0.0001; 3 vs 5 P = 0.0002; 3 vs 6 P = 0.0004; 4 vs 5 P = 0.0002; 4 vs 6 P = 0.0003; others n.s. |
| Fig. 10g | Time in stim chamber (% pretest time), repeated RTPA 10 min sessions | Ctrl 6, VTA 6, SNc 7; Sox14-Cre | Two-way RM ANOVA | StimDay x group F(10,80) = 2.867, P = 0.0042; StimDay F(3.48,55.68) = 16.17, P < 0.0001; Group F(2,16) = 5.40, P = 0.0162 | 0.26 [0.03, 0.33] | Sidak, at each day. Ctrl vs VTA StimDay4 P = 0.0008, test P = 0.0286; Ctrl vs SNc StimDay4 P = 0.0023; others n.s. |
| No panel (Results) | Freezing time in stim chamber (s), repeated RTPA 10 min sessions | Ctrl 6, VTA 6, SNc 7; Sox14-Cre | Two-way RM ANOVA | StimDay x group F(10,80) = 0.70, P = 0.72; StimDay F(1.723,27.57) = 0.72, P = 0.48; Group F(2,16) = 4.153, P = 0.035 | 0.08 [0.00, 0.09] | Sidak, at each day. No significant comparison; n.d. for several days |
| Fig. 10i | Locomotor activity (% time), open-field procedure 1 | Ctrl 6, VTA 6, SNc 7; Sox14-Cre | Two-way RM ANOVA | Time x group F(10,80) = 0.82, P = 0.61; Time F(3.736,59.77) = 12.42, P < 0.0001; Group F(2,16) = 0.38, P = 0.69 | 0.09 [0.00, 0.12] | Sidak, pairwise across minutes 1 to 6. 1 vs 2 P = 0.0003; 1 vs 3 P = 0.015; 1 vs 4, 1 vs 5 and 1 vs 6 P < 0.0001; others n.s. |
| Suppl. Fig. 17b | Time in stim chamber (s), RTPA 20 min session | Ctrl 5, LH-tVTA 6; C57BL/6J | Two-way RM ANOVA | Day x group F(1,9) = 44.63, P < 0.0001; Day F(1,9) = 46.52, P < 0.0001; Group F(1,9) = 30.69, P = 0.0004 | 0.83 [0.45, 0.91] | Sidak, pretest vs test. LH-tVTA P < 0.0001; Ctrl n.s. |
| Suppl. Fig. 18a | Average speed in stim chamber (m/min), RTPA 20 min session | Ctrl 6, LH-tVTA 4, tVTA 7; Sox14-Cre | Two-way RM ANOVA | Day x group F(2,14) = 11.77, P = 0.0010; Day F(1,14) = 20.30, P = 0.0005; Group F(2,14) = 4.91, P = 0.0242 | 0.63 [0.19, 0.76] | Sidak, pretest vs test. tVTA P < 0.0001; others n.s. |
| Suppl. Fig. 18b | Average speed in no-stim chamber (m/min), RTPA 20 min session | Ctrl 6, LH-tVTA 4, tVTA 7; Sox14-Cre | One-way ANOVA | F(2,14) = 0.2089, P = 0.814 | 0.03 [0.00, 0.21] | - |
| Suppl. Fig. 18c | Locomotor activity in no-stim chamber (% time), RTPA 20 min session | Ctrl 6, LH-tVTA 4, tVTA 7; Sox14-Cre | One-way ANOVA | F(2,14) = 0.0078, P = 0.9922 | 0.00 [0.00, 0.00] | - |
| Suppl. Fig. 18d | Freezing time in stim chamber (% time), RTPA 20 min session | Ctrl 6, LH-tVTA 4, tVTA 7; Sox14-Cre | Two-way RM ANOVA | Day x group F(2,14) = 3.57, P = 0.06; Day F(1,14) = 1.52, P = 0.24; Group F(2,14) = 1.12, P = 0.35 | 0.34 [0.00, 0.56] | - |
| Suppl. Fig. 18e | Locomotor activity without stim (% baseline), open-field procedure 2 | Ctrl 11, LH-tVTA 10, tVTA 7; Sox14-Cre and C57BL/6J | Two-way RM ANOVA | Day x group F(6,75) = 1.59, P = 0.16; Day F(2.35,58.88) = 5.86, P = 0.0030; Group F(2,25) = 1.033, P = 0.37 | 0.11 [0.00, 0.19] | - |
| Suppl. Fig. 18f | Freezing time without stim (s), open-field procedure 2 | Ctrl 11, LH-tVTA 10, tVTA 7; Sox14-Cre and C57BL/6J | Two-way RM ANOVA | Day x group F(6,75) = 1.13, P = 0.35; Day F(1.68,42.11) = 0.67, P = 0.49; Group F(2,25) = 0.54, P = 0.59 | 0.08 [0.00, 0.15] | - |

d, one row per test; a dash, not applicable; n.d., not determined. Alpha 0.05, two-tailed. Repeated-measures ANOVA with Greenhouse-Geisser correction on the repeated factor, GraphPad Prism 8.4.2. Effect size, partial eta squared with its 95% confidence interval, for the interaction in two-way ANOVA and for the group effect in one-way ANOVA. Post-hoc, significant comparisons with their P value; n.s., not significant.

**Supplementary Table 12 | Characterization of the twelve neurons recorded in vivo.**

| Neuron | Group | Trains | Pulses | Fid. (%) | Lat. (ms) | Jit. (ms) | Spont. (Hz) | First phase (ms) | Total duration (ms) |
|---|---|---|---|---|---|---|---|---|---|
| 28/A | tVTA | 3 | 30 | 93.3 | 2.11 | 0.238 | 10.077 | 1.045 | 2.655 |
| 30/C | tVTA | 10 | 100 | 97.0 | 2.31 | 0.240 | 16.095 | 0.767 | 2.330 |
| 30/D | tVTA | 10 | 100 | 99.0 | 2.91 | 0.287 | 11.426 | 1.030 | 3.380 |
| 31/C | tVTA | 11 | 110 | 95.5 | 2.96 | 0.251 | < 0.010 | 0.750 | 2.700 |
| 31/D | tVTA | 7 | 70 | 97.1 | 2.71 | 0.223 | < 0.023 | 0.600 | 2.550 |
| 31/H | tVTA | 12 | 120 | 94.2 | 3.01 | 0.263 | < 0.032 | 0.550 | 2.700 |
| 36/A | tVTA | 8 | 80 | 93.8 | 3.31 | 0.222 | < 0.030 | 0.300 | 2.850 |
| 36/C | tVTA | 7 | 70 | 77.1 | 3.76 | 0.179 | 0.084 | 0.300 | 2.850 |
| 37/A | tVTA | 12 | 120 | 95.8 | 3.61 | 0.275 | < 0.010 | 0.300 | 3.300 |
| 30/C DA | DA | 10 | - | - | - | - | 6.677 | 1.375 | 2.540 |
| 30/E | DA | 20 | - | - | - | - | 2.140 | 1.200 | 3.900 |
| 30/F | DA | 50 | - | - | - | - | 7.993 | 1.100 | 3.150 |

Fid., fidelity; Lat., median latency; Jit., jitter, s.d. of the peak latencies; Spont., spontaneous rate, "<" marking the neurons with no spontaneous spike, for which the rate is an upper Poisson limit; first phase and total duration, of the average action potential. The light response columns do not apply to the dopamine neurons.

**Supplementary Table 13 | Firing rate before, during and after the light, in the tVTA and the VTA neurons.**

**a | tVTA neurons**

| Neuron | Trains | Before (Hz) | Light (Hz) | After (Hz) | Spikes / pulse |
|---|---|---|---|---|---|
| 28/A | 3 | 9.33 ± 0.67 | 32.03 ± 3.61 | 10.67 ± 1.33 | 1.07 |
| 30/C | 10 | 16.40 ± 2.08 | 38.74 ± 1.38 | 9.40 ± 1.33 | 1.29 |
| 30/D | 10 | 9.20 ± 1.50 | 87.39 ± 2.95 | 5.40 ± 0.95 | 2.91 |
| 31/C | 11 | 0.00 ± 0.00 | 28.67 ± 0.47 | 0.00 ± 0.00 | 0.95 |
| 31/D | 7 | 0.00 ± 0.00 | 29.17 ± 0.55 | 0.00 ± 0.00 | 0.97 |
| 31/H | 12 | 0.00 ± 0.00 | 28.28 ± 0.58 | 0.00 ± 0.00 | 0.94 |
| 36/A | 8 | 0.00 ± 0.00 | 28.15 ± 0.79 | 0.00 ± 0.00 | 0.94 |
| 36/C | 7 | 0.00 ± 0.00 | 23.17 ± 2.34 | 0.00 ± 0.00 | 0.77 |
| 37/A | 12 | 0.00 ± 0.00 | 28.78 ± 0.58 | 0.00 ± 0.00 | 0.96 |

**b | VTA putative dopamine neurons**

| Neuron | Trains | Before (Hz) | Light (Hz) | After (Hz) | Inhibition (%) | Silent trains |
|---|---|---|---|---|---|---|
| 30/C | 10 | 9.20 ± 1.67 | 0.60 ± 0.40 | 11.60 ± 1.90 | -93.5 | 8/10 |
| 30/E | 20 | 3.20 ± 0.27 | 0.00 ± 0.00 | 2.20 ± 0.20 | -100 | 20/20 |
| 30/F | 50 | 8.28 ± 0.75 | 0.42 ± 0.15 | 12.80 ± 0.79 | -94.9 | 43/50 |

a, tVTA neurons; b, VTA putative dopamine neurons. Windows of Fig. 6d, f; mean ± s.e.m. across the trains of that neuron; silent trains, trains without any spike under the light.

**Supplementary Table 14 | Inter-animal reproducibility of the regional densities of the Sox14+ tVTA connectome.**

| Dataset | Mice | Regions | Inter-animal agreement | Median CV (%) | IQR of CV (%) | Mean CV (%) | Regions with CV ≤ 20 % |
|---|---|---|---|---|---|---|---|
| Afferent, labeled-cell density | 2 | 84 | Pearson r = 0.9627 on log10 density, 74 subcortical regions, P = 1.28e-42 | 15.2 | 6.7 to 29.7 | 19.2 | 50 of 84 |
| Efferent, fiber area | 4 | 110 | Mean pairwise Pearson r = 0.766 on log10 signal, 6 pairs, r from 0.67 to 0.85 | 62.1 | 51.5 to 80.0 | 65.2 | 2 of 110 |

CV, coefficient of variation per region, s.d. divided by the mean across animals, on the ipsilateral signal; median, interquartile range and mean taken over the regions. Afferent agreement on the 74 subcortical regions (Supplementary Fig. 13a); efferent agreement pairwise, on the regions above threshold in both animals of a pair (Supplementary Fig. 13c). No test compares the two datasets, whose units differ.

### Supplementary Table 15 | Complete connectome of the Sox14+ tVTA, 124 brain regions.

| Region | Allen full name | Super-group | Connectivity class | Rank aff. | Rank eff. | Afferent density (mean ± s.e.m., cells/mm²) | Efferent fiber area (mean ± s.e.m., % of region area, ipsilateral) |
|---|---|---|---|---|---|---|---|
| ACA | Anterior cingulate area | Cortex | Afferent-only | 83 | - | 0.6 ± 0.0 | - |
| AI | Agranular insular area | Cortex | Reciprocal (weak) | 81 | 105 | 1.0 ± 0.1 | 0.052 ± 0.013 |
| AON | Anterior olfactory nucleus | Cortex | Afferent-only | 79 | - | 1.5 ± 0.3 | - |
| CLA | Claustrum | Cortex | Reciprocal (weak) | 66 | 82 | 8.4 ± 1.2 | 0.123 ± 0.062 |
| DP | Dorsal peduncular area | Cortex | Afferent-only | 72 | - | 6.3 ± 2.4 | - |
| FRP | Frontal pole, cerebral cortex | Cortex | Efferent-only | - | 96 | - | 0.083 ± 0.044 |
| ILA | Infralimbic area | Cortex | Afferent-only | 76 | - | 3.1 ± 0.8 | - |
| LA | Lateral amygdalar nucleus | Cortex | Efferent-only | - | 107 | - | 0.044 ± 0.007 |
| NLOT | Nucleus of the lateral olfactory tract | Cortex | Efferent-only | - | 92 | - | 0.096 ± 0.034 |
| ORB | Orbital area | Cortex | Reciprocal (weak) | 78 | 90 | 1.6 ± 0.3 | 0.098 ± 0.036 |
| PIR | Piriform area | Cortex | Afferent-only | 84 | - | 0.2 ± 0.0 | - |
| PL | Prelimbic area | Cortex | Afferent-only | 77 | - | 1.9 ± 0.5 | - |
| VISC | Visceral area | Cortex | Afferent-only | 82 | - | 0.8 ± 0.3 | - |
| ACB | Nucleus accumbens | Striatum | Reciprocal (weak) | 49 | 87 | 20.0 ± 3.2 | 0.104 ± 0.025 |
| CP | Caudoputamen | Striatum | Efferent-only | - | 80 | - | 0.126 ± 0.038 |
| FS | Fundus of striatum | Striatum | Reciprocal (weak) | 69 | 65 | 7.2 ± 1.3 | 0.188 ± 0.031 |
| LS | Lateral septal nucleus | Striatum | Reciprocal (weak) | 80 | 55 | 1.3 ± 0.1 | 0.238 ± 0.061 |
| OT | Olfactory tubercle | Striatum | Efferent-only | - | 108 | - | 0.041 ± 0.007 |
| SF | Septofimbrial nucleus | Striatum | Efferent-only | - | 63 | - | 0.206 ± 0.027 |
| SH | Septohippocampal nucleus | Striatum | Efferent-only | - | 10 | - | 1.239 ± 0.390 |
| BST | Bed nuclei of the stria terminalis | Pallidum | Reciprocal (weak) | 38 | 84 | 35.8 ± 1.5 | 0.113 ± 0.029 |
| GPe | Globus pallidus, external segment | Pallidum | Efferent-only | - | 61 | - | 0.211 ± 0.081 |
| GPi | Globus pallidus, internal segment | Pallidum | Reciprocal (weak) | 70 | 13 | 6.8 ± 0.0 | 0.926 ± 0.124 |
| MA | Magnocellular nucleus | Pallidum | Reciprocal (weak) | 41 | 37 | 33.6 ± 0.2 | 0.376 ± 0.097 |
| MSC | Medial septal complex | Pallidum | Efferent-only | - | 65 | - | 0.188 ± 0.074 |
| SI | Substantia innominata | Pallidum | Reciprocal (weak) | 51 | 41 | 19.0 ± 1.2 | 0.340 ± 0.086 |
| AAA | Anterior amygdalar area | Amygdala | Reciprocal (weak) | 60 | 76 | 13.6 ± 3.4 | 0.139 ± 0.050 |
| CEA | Central amygdalar nucleus | Amygdala | Reciprocal (weak) | 62 | 60 | 10.9 ± 0.4 | 0.217 ± 0.080 |
| IA | Intercalated amygdalar nucleus | Amygdala | Efferent-only | - | 103 | - | 0.060 ± 0.028 |
| MEA | Medial amygdalar nucleus | Amygdala | Reciprocal (weak) | 75 | 89 | 4.4 ± 0.9 | 0.100 ± 0.033 |
| CL | Central lateral nucleus of the thalamus | Thalamus | Efferent-only | - | 68 | - | 0.178 ± 0.083 |
| CM | Central medial nucleus of the thalamus | Thalamus | Efferent-only | - | 66 | - | 0.185 ± 0.067 |
| Eth | Ethmoid nucleus of the thalamus | Thalamus | Efferent-only | - | 98 | - | 0.071 ± 0.009 |
| IAM | Interanteromedial nucleus of the thalamus | Thalamus | Efferent-only | - | 85 | - | 0.106 ± 0.043 |
| IMD | Intermediodorsal nucleus of the thalamus | Thalamus | Efferent-only | - | 20 | - | 0.675 ± 0.360 |
| LH | Lateral habenula | Thalamus | Reciprocal hub | 2 | 33 | 499.1 ± 4.7 | 0.423 ± 0.142 |
| MD | Mediodorsal nucleus of thalamus | Thalamus | Efferent-only | - | 52 | - | 0.270 ± 0.094 |
| MH | Medial habenula | Thalamus | Reciprocal (weak) | 58 | 102 | 16.3 ± 3.4 | 0.064 ± 0.010 |
| PF | Parafascicular nucleus | Thalamus | Reciprocal (weak) | 61 | 31 | 12.0 ± 0.2 | 0.449 ± 0.140 |
| PIL | Posterior intralaminar thalamic nucleus | Thalamus | Reciprocal (weak) | 67 | 62 | 8.2 ± 0.5 | 0.208 ± 0.080 |
| POL | Posterior limiting nucleus of the thalamus | Thalamus | Efferent-only | - | 79 | - | 0.128 ± 0.060 |
| PoT | Posterior triangular thalamic nucleus | Thalamus | Efferent-only | - | 70 | - | 0.162 ± 0.094 |
| PP | Peripeduncular nucleus | Thalamus | Efferent-only | - | 97 | - | 0.082 ± 0.025 |
| PR | Perireunensis nucleus | Thalamus | Efferent-only | - | 51 | - | 0.278 ± 0.129 |
| PT | Parataenial nucleus | Thalamus | Efferent-only | - | 75 | - | 0.140 ± 0.045 |
| PVT | Paraventricular nucleus of the thalamus | Thalamus | Efferent-only | - | 59 | - | 0.221 ± 0.105 |
| RE | Nucleus of reuniens | Thalamus | Efferent-only | - | 50 | - | 0.280 ± 0.135 |
| RH | Rhomboid nucleus | Thalamus | Efferent-only | - | 91 | - | 0.098 ± 0.028 |
| RT | Reticular nucleus of the thalamus | Thalamus | Efferent-only | - | 100 | - | 0.070 ± 0.034 |
| SMT | Submedial nucleus of the thalamus | Thalamus | Efferent-only | - | 67 | - | 0.184 ± 0.066 |
| SPA | Subparafascicular area | Thalamus | Reciprocal (weak) | 48 | 29 | 20.4 ± 6.2 | 0.486 ± 0.223 |
| SPF | Subparafascicular nucleus | Thalamus | Reciprocal (weak) | 71 | 44 | 6.3 ± 0.3 | 0.306 ± 0.080 |
| VAL | Ventral anterior-lateral complex of the thalamus | Thalamus | Efferent-only | - | 88 | - | 0.103 ± 0.034 |
| Xi | Xiphoid thalamic nucleus | Thalamus | Efferent-only | - | 57 | - | 0.226 ± 0.094 |
| ADP | Anterodorsal preoptic nucleus | Hypothalamus | Reciprocal (weak) | 25 | 53 | 78.4 ± 3.7 | 0.247 ± 0.040 |

| Region | Allen full name | Super-group | Connectivity class | Rank aff. | Rank eff. | Afferent density (mean ± s.e.m., cells/mm²) | Efferent fiber area (mean ± s.e.m., % of region area, ipsilateral) |
|---|---|---|---|---|---|---|---|
| AHN | Anterior hypothalamic nucleus | Hypothalamus | Reciprocal (weak) | 59 | 78 | 15.6 ± 5.7 | 0.132 ± 0.049 |
| ARH | Arcuate hypothalamic nucleus | Hypothalamus | Afferent-only | 56 | - | 17.1 ± 1.8 | - |
| AVP | Anteroventral preoptic nucleus | Hypothalamus | Reciprocal hub | 14 | 25 | 120.7 ± 11.4 | 0.537 ± 0.154 |
| AVPV | Anteroventral periventricular nucleus | Hypothalamus | Reciprocal (weak) | 42 | 104 | 31.4 ± 3.0 | 0.056 ± 0.016 |
| DMH | Dorsomedial nucleus of the hypothalamus | Hypothalamus | Reciprocal (weak) | 43 | 49 | 31.0 ± 2.4 | 0.282 ± 0.084 |
| LHA | Lateral hypothalamic area | Hypothalamus | Reciprocal (weak) | 27 | 6 | 73.4 ± 7.5 | 1.897 ± 0.562 |
| LM | Lateral mammillary nucleus | Hypothalamus | Reciprocal hub | 23 | 17 | 79.3 ± 13.4 | 0.734 ± 0.126 |
| LPO | Lateral preoptic area | Hypothalamus | Reciprocal hub | 18 | 9 | 107.1 ± 23.6 | 1.337 ± 0.635 |
| MEPO | Median preoptic nucleus | Hypothalamus | Reciprocal (weak) | 37 | 101 | 41.1 ± 11.8 | 0.069 ± 0.013 |
| MM | Medial mammillary nucleus | Hypothalamus | Reciprocal (weak) | 19 | 73 | 105.7 ± 12.1 | 0.151 ± 0.037 |
| MPN | Medial preoptic nucleus | Hypothalamus | Afferent-only | 32 | - | 58.1 ± 3.6 | - |
| MPO | Medial preoptic area | Hypothalamus | Reciprocal (weak) | 10 | 36 | 125.8 ± 22.4 | 0.380 ± 0.111 |
| PeF | Perifornical nucleus | Hypothalamus | Reciprocal (weak) | 53 | 32 | 18.4 ± 5.0 | 0.427 ± 0.091 |
| PH | Posterior hypothalamic nucleus | Hypothalamus | Reciprocal (weak) | 30 | 22 | 65.5 ± 0.2 | 0.638 ± 0.298 |
| PMd | Dorsal premammillary nucleus | Hypothalamus | Reciprocal hub | 16 | 16 | 109.5 ± 1.6 | 0.761 ± 0.351 |
| PMv | Ventral premammillary nucleus | Hypothalamus | Reciprocal (weak) | 20 | 93 | 93.1 ± 2.1 | 0.095 ± 0.063 |
| PS | Parastrial nucleus | Hypothalamus | Reciprocal hub | 5 | 28 | 248.6 ± 26.8 | 0.505 ± 0.130 |
| PSTN | Parasubthalamic nucleus | Hypothalamus | Reciprocal (weak) | 35 | 4 | 43.0 ± 4.9 | 4.662 ± 1.772 |
| PVH | Paraventricular hypothalamic nucleus | Hypothalamus | Reciprocal (weak) | 63 | 71 | 10.7 ± 2.2 | 0.157 ± 0.045 |
| PVHd | Paraventricular hypothalamic nucleus, descending division | Hypothalamus | Reciprocal (weak) | 39 | 42 | 35.5 ± 8.7 | 0.332 ± 0.111 |
| PVp | Periventricular hypothalamic nucleus, posterior part | Hypothalamus | Afferent-only | 45 | - | 22.9 ± 0.4 | - |
| RCH | Retrochiasmatic area | Hypothalamus | Efferent-only | - | 83 | - | 0.119 ± 0.036 |
| SBPV | Subparaventricular zone | Hypothalamus | Efferent-only | - | 69 | - | 0.174 ± 0.053 |
| SO | Supraoptic nucleus | Hypothalamus | Efferent-only | - | 39 | - | 0.369 ± 0.164 |
| STN | Subthalamic nucleus | Hypothalamus | Efferent-only | - | 8 | - | 1.363 ± 0.416 |
| SUM | Supramammillary nucleus | Hypothalamus | Reciprocal hub | 21 | 14 | 92.8 ± 10.8 | 0.814 ± 0.245 |
| TM | Tuberomammillary nucleus | Hypothalamus | Reciprocal (weak) | 28 | 48 | 72.9 ± 6.0 | 0.292 ± 0.115 |
| TU | Tuberal nucleus | Hypothalamus | Reciprocal (weak) | 52 | 74 | 18.9 ± 3.4 | 0.140 ± 0.025 |
| VLPO | Ventrolateral preoptic nucleus | Hypothalamus | Efferent-only | - | 64 | - | 0.194 ± 0.033 |
| VMH | Ventromedial hypothalamic nucleus | Hypothalamus | Afferent-only | 73 | - | 5.5 ± 2.2 | - |
| VMPO | Ventromedial preoptic nucleus | Hypothalamus | Reciprocal (weak) | 34 | 72 | 47.8 ± 1.2 | 0.151 ± 0.080 |
| ZI | Zona incerta | Hypothalamus | Reciprocal (weak) | 44 | 26 | 27.0 ± 3.1 | 0.508 ± 0.139 |
| CUN | Cuneiform nucleus | Midbrain | Reciprocal (weak) | 36 | 94 | 41.6 ± 9.0 | 0.090 ± 0.030 |
| DR | Dorsal nucleus raphe | Midbrain | Reciprocal hub | 1 | 3 | 1126.2 ± 88.5 | 4.859 ± 1.349 |
| IPN | Interpeduncular nucleus | Midbrain | Reciprocal hub | 6 | 27 | 233.7 ± 16.1 | 0.508 ± 0.291 |
| MRN | Midbrain reticular nucleus | Midbrain | Reciprocal (weak) | 33 | 7 | 49.3 ± 2.2 | 1.551 ± 0.314 |
| PAG | Periaqueductal gray | Midbrain | Reciprocal (weak) | 31 | 21 | 64.2 ± 3.4 | 0.664 ± 0.281 |
| PPN | Pedunculopontine nucleus | Midbrain | Reciprocal hub | 26 | 5 | 77.2 ± 1.5 | 2.203 ± 0.103 |
| PRT | Pretectal region | Midbrain | Reciprocal (weak) | 74 | 77 | 5.2 ± 0.1 | 0.136 ± 0.091 |
| SCm | Superior colliculus, motor related | Midbrain | Afferent-only | 54 | - | 18.3 ± 0.8 | - |
| SCO | Subcommissural organ | Midbrain | Efferent-only | - | 106 | - | 0.046 ± 0.008 |
| SNc | Substantia nigra, compact part | Midbrain | Reciprocal hub | 4 | 2 | 272.3 ± 2.2 | 20.039 ± 4.277 |
| SNr | Substantia nigra, reticular part | Midbrain | Reciprocal (weak) | 47 | 11 | 21.0 ± 2.8 | 1.178 ± 0.401 |
| VTA-cx | VTA complex | Midbrain | Reciprocal hub | 11 | 1 | 124.9 ± 30.2 | 26.369 ± 6.391 |
| VTN | Ventral tegmental nucleus | Midbrain | Efferent-only | - | 46 | - | 0.299 ± 0.184 |
| Acs5 | Accessory trigeminal nucleus | Pons | Efferent-only | - | 35 | - | 0.396 ± 0.337 |
| CS | Superior central nucleus raphe | Pons | Reciprocal hub | 3 | 12 | 309.4 ± 25.8 | 1.066 ± 0.684 |
| DTN | Dorsal tegmental nucleus | Pons | Reciprocal (weak) | 24 | 54 | 79.3 ± 27.3 | 0.241 ± 0.153 |
| LC | Locus coeruleus | Pons | Reciprocal hub | 17 | 19 | 109.1 ± 31.3 | 0.708 ± 0.315 |
| LDT | Laterodorsal tegmental nucleus | Pons | Reciprocal hub | 8 | 18 | 168.2 ± 1.1 | 0.726 ± 0.317 |
| NI | Nucleus incertus | Pons | Reciprocal (weak) | 15 | 58 | 118.4 ± 29.2 | 0.225 ± 0.074 |
| PB | Parabrachial nucleus | Pons | Reciprocal hub | 13 | 15 | 120.9 ± 1.2 | 0.806 ± 0.244 |
| PC5 | Parvicellular motor 5 nucleus | Pons | Efferent-only | - | 99 | - | 0.071 ± 0.023 |
| PCG | Pontine central gray | Pons | Reciprocal (weak) | 9 | 38 | 128.7 ± 11.3 | 0.369 ± 0.140 |
| PRNc | Pontine reticular nucleus, caudal part | Pons | Reciprocal (weak) | 40 | 30 | 33.8 ± 0.7 | 0.468 ± 0.120 |

| Region | Allen full name | Super-group | Connectivity class | Rank aff. | Rank eff. | Afferent density (mean ± s.e.m., cells/mm²) | Efferent fiber area (mean ± s.e.m., % of region area, ipsilateral) |
|---|---|---|---|---|---|---|---|
| PRNr | Pontine reticular nucleus | Pons | Reciprocal (weak) | 29 | 24 | 67.1 ± 13.2 | 0.553 ± 0.195 |
| RPO | Nucleus raphe pontis | Pons | Reciprocal (weak) | 7 | 34 | 227.1 ± 18.0 | 0.410 ± 0.138 |
| SG | Supragenual nucleus | Pons | Efferent-only | - | 56 | - | 0.235 ± 0.044 |
| SLD | Sublaterodorsal nucleus | Pons | Reciprocal (weak) | 22 | 40 | 86.1 ± 8.7 | 0.348 ± 0.122 |
| TRN | Tegmental reticular nucleus | Pons | Reciprocal (weak) | 65 | 43 | 8.7 ± 2.9 | 0.309 ± 0.136 |
| GRN | Gigantocellular reticular nucleus | Medulla | Reciprocal (weak) | 46 | 45 | 21.1 ± 4.0 | 0.305 ± 0.124 |
| IRN | Intermediate reticular nucleus | Medulla | Reciprocal (weak) | 68 | 86 | 7.5 ± 1.8 | 0.104 ± 0.037 |
| MARN | Magnocellular reticular nucleus | Medulla | Reciprocal (weak) | 55 | 47 | 18.1 ± 1.9 | 0.296 ± 0.113 |
| RaM | Medullary raphe nuclei | Medulla | Reciprocal hub | 12 | 23 | 121.1 ± 18.0 | 0.622 ± 0.253 |
| MV | Medial vestibular nucleus | Medulla | Reciprocal (weak) | 50 | 95 | 19.0 ± 2.6 | 0.086 ± 0.038 |
| NTB | Nucleus of the trapezoid body | Medulla | Efferent-only | - | 109 | - | 0.039 ± 0.003 |
| PGRN | Paragigantocellular reticular nucleus | Medulla | Afferent-only | 64 | - | 10.5 ± 1.9 | - |
| PPY | Parapyramidal nucleus | Medulla | Efferent-only | - | 81 | - | 0.125 ± 0.079 |
| PRP | Nucleus prepositus | Medulla | Afferent-only | 57 | - | 16.9 ± 0.0 | - |

Regions in the order of Fig. 8a, super-group by super-group and alphabetically within each. Ranks among the 84 afferent and the 110 efferent regions, 1 the densest. Reciprocal hub, top 30 % of both lists (afferent rank ≤ 26, efferent rank ≤ 33); weak-reciprocal, detected in both datasets below that threshold. Mean ± s.e.m. across animals, n = 2 (afferent) and n = 4 (efferent); a dash, not detected in that dataset. The intra-tVTA labeling of Fig. 8a is not listed. RaM, medullary raphe nuclei (magnus, obscurus and pallidus merged).

**Supplementary Table 16 | Connectivity composition of the Sox14+ tVTA connectome per anatomical super-group.**

| Super-group | n | Afferent-only | Reciprocal hubs | Weak-reciprocal | Efferent-only | Reciprocal (%) | Hubs |
|---|---|---|---|---|---|---|---|
| Cortex | 13 | 7 | 0 | 3 | 3 | 23.1 | - |
| Striatum | 7 | 0 | 0 | 3 | 4 | 42.9 | - |
| Pallidum | 6 | 0 | 0 | 4 | 2 | 66.7 | - |
| Amygdala | 4 | 0 | 0 | 3 | 1 | 75.0 | - |
| Thalamus | 24 | 0 | 1 | 5 | 18 | 25.0 | LH |
| Hypothalamus | 33 | 4 | 6 | 18 | 5 | 72.7 | AVP, LM, LPO, PMd, PS, SUM |
| Midbrain | 13 | 1 | 5 | 5 | 2 | 76.9 | DR, IPN, PPN, SNc, VTA-cx |
| Pons | 15 | 0 | 4 | 8 | 3 | 80.0 | CS, LC, LDT, PB |
| Medulla | 9 | 2 | 1 | 4 | 2 | 55.6 | RaM |
| Total | 124 | 14 | 17 | 53 | 40 | 56.5 | 17 hubs |

Counts of Fig. 8b. Reciprocal (%), proportion of the regions of the super-group detected in both directions, hubs and weak-reciprocal together.

**Supplementary Table 17 | Hemispheric lateralization of the Sox14+ tVTA connectome per anatomical super-group.**

| Super-group | Dataset | Regions | Strongly contralateral | Contra-dominant | Bilateral | Ipsi-dominant | Strongly ipsilateral |
|---|---|---|---|---|---|---|---|
| Cortex | Afferent | 10 | 0 | 0 | 5 | 4 | 1 |
| Cortex | Efferent | 6 | 0 | 0 | 0 | 0 | 6 |
| Striatum | Afferent | 3 | 0 | 0 | 1 | 2 | 0 |
| Striatum | Efferent | 7 | 0 | 0 | 0 | 0 | 7 |
| Pallidum | Afferent | 4 | 0 | 0 | 1 | 2 | 1 |
| Pallidum | Efferent | 6 | 0 | 0 | 0 | 0 | 6 |
| Amygdala | Afferent | 3 | 0 | 0 | 1 | 2 | 0 |
| Amygdala | Efferent | 4 | 0 | 0 | 0 | 0 | 4 |
| Thalamus | Afferent | 6 | 0 | 0 | 2 | 4 | 0 |
| Thalamus | Efferent | 24 | 0 | 0 | 5 | 8 | 11 |
| Hypothalamus | Afferent | 28 | 0 | 1 | 17 | 9 | 1 |
| Hypothalamus | Efferent | 29 | 0 | 0 | 3 | 10 | 16 |
| Midbrain | Afferent | 11 | 0 | 0 | 11 | 0 | 0 |
| Midbrain | Efferent | 12 | 0 | 0 | 3 | 5 | 4 |
| Pons | Afferent | 12 | 0 | 1 | 9 | 2 | 0 |
| Pons | Efferent | 15 | 0 | 0 | 2 | 8 | 5 |
| Medulla | Afferent | 7 | 0 | 0 | 6 | 1 | 0 |
| Medulla | Efferent | 7 | 0 | 0 | 1 | 6 | 0 |
| All super-groups | Afferent | 84 | 0 | 2 | 53 | 26 | 3 |
| All super-groups | Efferent | 110 | 0 | 0 | 14 | 37 | 59 |

Number of regions per class, super-groups in the order of Fig. 8a. LI as in Fig. 8e; bilateral, $|LI| \leq 0.20$; dominant, $0.20 < |LI| \leq 0.50$; strongly lateralized, $|LI| > 0.50$. Afferent, n = 2 mice; efferent, n = 4 mice; no test on these counts.
